# Multi-behavioral phenotyping in early-life-stage zebrafish for identifying disruptors of non-associative learning

**DOI:** 10.1101/2024.09.25.613874

**Authors:** David Leuthold, Nadia K. Herold, Jana Nerlich, Kristina Bartmann, Ilka Scharkin, Stefan J. Hallermann, Nicole Schweiger, Ellen Fritsche, Tamara Tal

## Abstract

**Background:** The vertebrate nervous system is vulnerable to chemical toxicity and the widespread release of chemicals into the environment outstrips the capacity to assess their safety. The zebrafish (*Danio rerio*) is a powerful vertebrate model that can bridge the gap between *in vitro* and mammalian-based *in vivo* studies. However, the behavior-rich repertoire of larval zebrafish, a 3R-compliant model amenable to higher-throughput chemical screens, has yet to be fully deployed to identify and characterize chemical compounds that cause neurotoxicity.

**Objective:** We sought to establish a multi-behavioral phenotyping approach in larval zebrafish to identify and mechanistically elucidate neuroactive chemicals, with particular focus on chemical compounds that affect non-associative habituation learning.

**Methods:** We devised a battery of automated behavior assays in larval zebrafish. The battery captures stereotypical visual and acoustic behaviors including habituation, a form of non-associative learning. To elucidate mechanisms underlying exposure-induced behavioral alterations in zebrafish, *in silico* target predictions, pharmacological interventions, patch-clamp recordings in cultured mouse cortical neurons, and human multi-neurotransmitter (hMNR) assay in 3D BrainSpheres were used.

**Results:** Known pharmacological modulators of habituation in zebrafish evoked distinct behavioral patterns. By screening chemicals positive for *ex vivo N*-methyl-D-aspartate receptor (NMDAR) modulation, we identified chlorophene, a biocide that caused sedation, paradoxical excitation, and reduced habituation in zebrafish. Using *in silico* target predictions and pharmacological interventions, we discovered that chlorophene acts via gamma-aminobutyric acid A receptors (GABA_A_Rs), a previously unknown target site. Orthogonal validation in cultured mouse cortical neurons and human stem cell-derived BrainSpheres confirmed chlorophene’s interaction with GABA_A_Rs. Chlorophene’s behavioral profile resembled that of flupirtine, a Kv7 potassium channel (M-current) activator, suggesting that habituation deficits stem from M-current rather than GABA_A_R modulation.

**Conclusions:** These studies combined a series of behavior assays in a phenotypically rich, rapid, and inexpensive non-mammalian vertebrate test system to screen chemicals for neurotoxicity. Together with *in silico* target predictions and mouse- and human-based models, our findings establish multi-behavioral phenotyping in zebrafish as a powerful toolkit for neurotoxicity testing and mechanism identification, with relevance for humans.

## Introduction

The nervous system is one of the most vulnerable organ systems to toxic injury due to the interdependence of neurodevelopmental processes and where perturbations can result in profound structural and/or functional consequences^1^. Epidemiological studies indicate that exposure to pesticides (e.g., rotenone, paraquat, chlorpyrifos), metals (e.g., lead, mercury), neuroactive drugs (e.g. benzodiazepines, anticholinergics), or industrial chemicals (e.g., polychlorinated biphenyls, brominated flame retardants, phthalates) contributes to neurobehavioral pathologies such as Parkinson’s disease^2,3^, Alzheimer’s disease^4–6^, attention deficit hyperactivity disorder^7,8^, and other neurodevelopmental deficits^9^. These associations are supported by experimental findings in mice^10–14^, rats^15,16^, non-human primates^17,18^, and other, non-mammalian models^19,20^. Despite these associations, most neurotoxicants remain unknown^9^. This is highlighted by the fact that, to date, fewer than 200 chemicals^21^, out of a predicted global inventory of registered chemicals and mixtures of more than 350,000 substances^22^, have been tested according to guideline neurotoxicity studies including acute (28 d), sub-chronic (90 d), and chronic (≥1 year) adult neurotoxicity^23^ and developmental neurotoxicity (DNT) studies^24,25^, and are publicly available in the US EPA ToxRef database^26^. This lack of information exists because DNT *in vivo* studies are typically conducted in rats^24,25^, costly (∼$1,000,000 per chemical^27^), laborious (∼1 year per chemical^27,28^), and raise ethical concerns^27,29^.

As untested chemicals should not be presumed safe, a new testing paradigm is needed to assess the ever-expanding universe of potentially neurotoxic chemicals^9,27^. To fill this gap, an integrated testing strategy that combines *in silico* models, *in vitro* techniques, and alternative animal models, collectively referred to as New Approach Methods (NAMs), has been proposed^27,30,31^. *In silico* methods (e.g. grouping and read-across, quantitative structure-activity relationships) can be used to predict ADME (Absorption, Distribution, Metabolism, Excretion) processes (e.g. blood-brain-barrier penetration), to model physiologically-based toxicokinetics, and to predict mechanisms based on relationships between chemical structure/properties and biological activity^32^. *In silico* approaches rely on experimental data. International efforts to enhance neurotoxicity testing involve the establishment of a DNT *in vitro* testing battery that depicts key neurodevelopmental processes including proliferation, migration, differentiation, apoptosis, neurite outgrowth, synaptogenesis, and functional network formation in a suite of rodent- and human-based cell models^33,34^. The underlying assumption of this approach is that the alteration of any of these processes is indicative of DNT^33^. Despite advances in depicting the processes that orchestrate nervous system development and function *in vitro*, more complex functional endpoints at the organismal level, including complex behaviors, remain elusive. In particular, *in vitro* assays fail to capture behavior endpoints assessed in guideline DNT studies including motor activity, motor and sensory function, learning, and memory^24^.

The zebrafish is a powerful vertebrate model that can bridge the gap between *in vitro* and mammalian-based *in vivo* studies. Although zebrafish are phylogenetically more distant from humans than rodents, 70% of human genes and 82% of disease-associated human genes have a zebrafish orthologue^35^. Embryo-larval stages of zebrafish up to 5 d post fertilization (dpf) are non-protected^36,37^ and therefore considered to be an alternative to mammalian testing^38^. The high reproduction rate, rapid *ex utero* development, and small size of embryo-larval stages make zebrafish amenable to medium-to-high-throughput screens. Early life-stage zebrafish provide a diverse repertoire of behaviors such as spontaneous tail contractions^39^, touch-evoked responses^40,41^, visual and acoustic startle responses^42^, phototaxis^43^, learning and memory^44,45^. While multi-behavioral phenotyping in zebrafish has been widely applied in drug discovery^46,47^, molecular target identification^48,49^, and to study neurological disorders^50–52^, the behavior-rich repertoire of zebrafish has yet to be fully deployed to identify and characterize chemicals that cause neurotoxicity.

The aim of this study is to identify and mechanistically evaluate chemical compounds that cause deficits in learning and memory. First, we developed a multi-behavioral phenotyping battery in early-life-stage zebrafish to quantify various visual and acoustic behaviors including non-associative learning. Second, we validated the assay battery using known pharmacological modulators of habituation learning. Third, we performed a targeted chemical screen to identify chemical compounds that affect learning and memory in zebrafish. We then combined pharmacological interventions, phenocopy strategies, and electrophysiology in mouse- and human-based models to identify mechanisms underlying phenotypic changes and to demonstrate cross-species relevance. Overall, the zebrafish-based multi-behavioral phenotyping approach represents an efficient tool for the identification and mechanistic elucidation of neurotoxic environmental chemicals with relevance across taxa, including humans.

## Methods

### Zebrafish maintenance and embryo collection

All protocols and procedures involving zebrafish were conducted according to German and European animal protection standards and were approved by the Government of Saxony (Landesdirektion Leipzig, file number 75-9185.64). Adult zebrafish (*Danio rerio*) of the UFZ-OBI/WIK strain, an outcross between the in-house UFZ-OBI strain and the WIK (Wild Indian Karyotype) strain, were used in this study. The UFZ-OBI strain originated from a local hardware store, while the WIK strain, a wild-type zebrafish lineage collected in India, was obtained from the European Zebrafish Resource Center (EZRC) at the Karlsruhe Institute of Technology (KIT). Fish were maintained on a 14-hour light, 10-hour dark cycle at 28 °C and were fed twice daily on weekdays and once daily on weekends with live brine shrimp and/or SDS 400 dry feed (Special Diets Services). Fertilized eggs were collected from group breeds containing 120-140 fish/120 L water. To minimize microbial burden and reduce potential developmental disruptions, we opted to bleach embryos for 5 min with 0.05% NaOCl at 3-4 h post-fertilization (hpf) following established protocols^53^. Embryos were raised in crystallization dishes at a density of 50 embryos per 100 mL standard dilution water (ISO 7346-3; 80 mM CaCl_2_·2H_2_O, 20 mM MgSO_4_·7H_2_O, 31 mM NaHCO_3_, 3.1 mM KCl, pH 7.4) until 4 dpf. Viable, hatched, and morphologically normal larvae with an inflated posterior swim bladder were individually distributed into single wells of clear polystyrene flat-bottom 96-square-well plates (Cytiva, Marlborough, MA, USA) filled with 200 µL standard dilution water. Plates were covered with microseal ‘A’ film (Bio-Rad Laboratories, Hercules, CA, USA) and microplate lids (Cytiva, Marlborough, MA, USA). Plates were stored at 28 °C on a 14-h light to 10-h dark cycle prior chemical exposure at 5 dpf. To control for potential circadian influences on behavior, we used two independent, time-shifted spawning units to ensure that all larvae were tested at 5 dpf at the same circadian time relative to their native diurnal rhythm, thereby minimizing variability across replicates.

### Zebrafish chemical exposure

Chemical compounds are listed in **Table S1**. To validate our behavioral assay battery, we selected pharmacological modulators previously shown to influence habituation in larval zebrafish, primarily based on landmark studies by Best et al.^45^, Wolman et al.^44^, and Roberts et al.^54^ (**Table S2**). Selection criteria also included representation of diverse mechanisms of action to capture a broad range of neurobehavioral effects. The selected compounds were dissolved in standard dilution water before administration. Environmentally-relevant chemical compounds, selected from *ex vivo* studies, were dissolved in dimethyl sulfoxide (DMSO, Merck, 1.02952.1000, ≥99.9% purity). 100 mM chemical stock aliquots, prepared in DMSO, were stored at −80 °C. Serial dilutions were prepared from 200 µM chemical stock solutions in standard dilution water (OECD Test Guideline 236), adjusted to pH 7.4, ensuring consistent ionic strength across the study within the zebrafish pH tolerance range (5-10)^55^ and minimizing potential ion-related confounding effects on behavioral assays and chemical exposure outcomes. At 5 dpf, larvae were exposed to 200 µL of 2× exposure solutions for a final volume of 400 µL per well. Each compound was tested at 7-10 concentrations ranging from 0.1-100 µM, unless otherwise indicated. This range was selected to account for potential differences in uptake kinetics and to ensure adequate bioavailability, particularly for more hydrophobic compounds (logK_OW_>3). When DMSO was used, the control group was exposed to an equal volume for a final concentration of 0.01% DMSO. Compounds were tested in 8-24 replicate wells across 2-4 replicate plates using larvae from two independent spawning events. To detect acute, receptor-mediated effects, larvae were exposed for 40 min (28 °C, light on) prior to behavior analysis, a duration chosen based on previous studies demonstrating its effectiveness in capturing neuroactive effects^47,56–58^. For behavior phenotype rescue experiments, larvae in 200 µL standard dilution water were pre-treated with 100 µL 3× picrotoxin solution at a concentration of 64.6 µM (21.5 µM final concentration) or vehicle for 15 min, followed by addition of either 100 µL 4× chlorophene solution at 27.3 µM (6.8 µM final concentration), 21.5 µM picrotoxin, a mixture of chlorophene and picrotoxin (6.8 µM chlorophene, 21.5 µM picrotoxin final concentration), or vehicle. The final concentration of DMSO was 0.01% for all groups. Plates were incubated for an additional 25 min before behavior acquisition.

### Zebrafish automated behavior assay battery and motor activity quantification

96-square well plates containing a single 5 dpf larva per well were loaded into a ZebraBox behavior apparatus (ViewPoint) and the measurement was initiated. The assay commenced with an initial 21 min acclimation period (1 min 100% light, 20 min 0% light). The temperature was maintained at 28°C for the duration of the experiment. Videos were recorded at 25 frames per second and ZebraLab video tracking software (ViewPoint) in ‘quantization mode’ (sensitivity threshold = 15) to quantify locomotor activity using pixel intensity changes per second for each of the 96 evenly-spaced, well-specific regions. In a battery of 11 behavioral assays, sequential variations in stimulus modalities (**Table S3**) were applied to stimulate changes in locomotor activity, that were optimized considering visual (light- and dark-cycle durations) and acoustic stimulus parameters (sound frequency, volume, inter-stimulus interval (ISI), inter-bout interval), and developmental stage (**Fig. S1** and **S2**). Infrared light was used to continuously illuminate the measuring chamber and visual light at 100% nominal intensity was determined to be 7.5 klux (LI-250 light meter, LI-COR). Acoustic stimuli with a frequency of 300 Hz were applied at nominal volumes of 75% (65 dB for ASR1) and 100% (75 dB for ASR2, ASR3, ASH1-5).

Motor activity data (Δpixel/second/well) was analyzed using R scripts for data processing (dplyr), distance-based analysis and clustering (concaveman, parallel), multidimensional scaling (MDS; ggforce), statistical analysis (scales), and data visualization (ggplot2, RColorBrewer, ComplexHeatmap, circlize, grid). To quantify motor activity, cumulative pixel intensity changes per second were calculated per well for each assay. The average motor activity per second during the final minute of the acclimation period was used to quantify baseline activity (assay BSL). Motor activity following dark-to-light transitions was divided into two periods reflecting rapid visual startle responses lasting 1 s (assay VSR1) and low basal activity in the following 9:59 min (assay VMR1). Subsequent changes in motor activity following light-to-dark transitions were binned into three periods including a one-second visual startle response (assay VSR2), a gradual increase in motor activity in the following 3:59 min (assay VMR2), and a transient decrease in motor activity to baseline level within the subsequent 16 min of the dark period (assay VMR3). Then, acoustic startle responses to five low-intensity (assay ASR1) and high-intensity (assay ASR2) 1-s acoustic stimuli, separated by 1-min ISIs, were averaged across five trials. Motor responses to 30 high-intensity acoustic stimuli (1-s with 1-s ISI) were condensed into a single metric reflecting acoustic startle habituation (assay ASH1) comprising the cumulative motor activity during the final ten taps scaled to the overall activity during the initial and final ten taps to account for individual differences in baseline startle activity^59^. Out of five repetitive habituation bouts with an inter-bout interval of 1 min, the ratio of motor activity during taps in the final bout (ASH5) relative to the sum of activities in the initial (ASH1) and final bout (ASH5) was calculated to quantify potentiation of habituation (ASH1/5). Habituation trials were followed by a three-minute rest period. Then, five high-intensity acoustic stimuli (1-s, 1-min ISI; ASR3) that were averaged across trials and scaled to the overall responses summed across ASR2 and ASR3 assays to quantify memory retention (ASR2/3). Following each experiment, dead larvae, identified upon visual inspection, were excluded from behavior data analyses.

### Statistics, behavioral profiling, hierarchical clustering, and multidimensional scaling for zebrafish behavior data

Statistical analyses were performed on 11 behavior metrics including baseline activity (BSL), visual startle responses (VSR1 and VSR2), visual motor responses (VMR1-VMR3), acoustic startle responses (ASR1 and ASR2), acoustic startle habituation (ASH1), potentiation of habituation (ASH1/5), and memory retention (ASR2/3). Given the diverse, mostly nonparametric probability distributions of data across different behavior metrics, we chose a universally applicable, nonparametric bootstrapping method that is not limited by a model assumption^60^. First, the absolute value (two-sided test) of the difference in medians between untreated control and each treatment was calculated as a test statistic. Then, a *N* (number of observations to sample) times *B* (number of bootstrap samples; set to 10,000) matrix was created, where each column represented a bootstrap resample (with replacement) of the observed data. Third, the bootstrap test statistic (absolute value of the difference in medians) was calculated for each bootstrap resample per column, considering the number of observations per group. Finally, the p-value was calculated reflecting the proportion of bootstrap test statistics that were equal or greater than the observed test statistic^61,62^. For multiple comparisons, the Kruskal-Wallis rank sum test was used and the R package “agricolae”^63^ was used to report results of all pairwise comparisons using compact letter displays. A Benjamini-Hochberg correction was used to adjust for multiple comparisons in order to control for family-wise Type I error rates. Results were considered significant if p_adjusted_<0.05. To identify modulators of non-associative habituation learning, we defined a ‘hit’ as a compound that resulted in a statistically significant higher or lower acoustic startle habituation response (ASH1) in at least one tested concentration. To calculate 95% confidence intervals of the median for each one-second time bin, 1,000 bootstraps (with replacement) were sampled from the observed data and the percentile method was used to determine the lower (2.5^th^ percentile) and upper (97.5^th^ percentile) confidence interval bounds^64^. As a measure of effect size, strictly standardized median difference (SSMD) values were calculated as follows: SSMD=x^∼^_treatment_-x^∼^_control_/(MAD^2^ + MAD^2^)^1/2^, where x^∼^ and MAD represent median and median absolute deviation^65^. SSMD calculations were performed on each of the 11 behavior metrics, collectively representing a compound’s behavioral profile at a given concentration. Hierarchical clustering was performed on behavioral profiles using k-means clustering in the R package ComplexHeatmap^66^. For MDS, two-dimensional coordinates for each behavioral profile were computed in R using metric MDS on Euclidean distance matrices.

### Identification of candidate compounds for zebrafish behavior-based screening

The US EPA CompTox Chemicals Dashboard (https://comptox.epa.gov/dashboard/) was searched for assay endpoint names and genes using the term ‘NMDA,’ with data collected between July and August 2021. Two assays were identified, including “NVS_LGIC_rGluNMDA_Agonist” and “NVS_LGIC_rGluNMDA_MK801_Agonist”, both representing tissue-based, cell-free radioligand binding assays on extracted gene-proteins from rat forebrain membranes relating to the gene *grin1*. Detailed assay information can be found in **Table S4**. Among 81 unique chemical compounds present in at least one of the two datasets, 26 compounds were annotated as hits (**Table S5**). Positive hits were included in the zebrafish behavior screen if their AC_50_ (active concentration causing 50% loss in scintillation count) values were below the lower bound of cytotoxicity^67^ or the predicted 96-h LC_50_ in fish (ECOSAR v2.0, U.S. EPA) (**Table S6**). Additionally, two compounds annotated as inactive, including cypermethrin and ascorbate, were selected as negative reference compounds. Chemical exposures were performed as described above.

### Molecular target prediction

The Similarity Ensemble Approach (SEA^68^) was used to predict molecular target sites for MK-801 (**Table S7**) and chlorophene (**Table S8**).

### Mouse cortical cultures

Neocortical neuronal cultures from P0 to P1 mice were prepared as previously described^69^. Briefly, mice were decapitated and cerebral cortices were removed, dissected and enzymatically digested with papain (Sigma, CAS 9001-73-4) or trypsine (Sigma, CAS 9007-07-7) in the presence of DNAse (Sigma, CAS 9003-98-9), followed by mechanical dissociation and centrifugation through a cushion of 4% bovine serum albumin (Sigma, CAS 9048-46-8). These steps were completed using Hibernate medium (ThermoFisher). Cells were then plated onto Poly-L-Lysine (Sigma, CAS 9001-73-4) coated coverslips in 24 well plates. For each coverslip, 25-30 k cells were allowed to settle in a 40 μL drop for approximately 30 min and then each well was filled with 500 μL of NeurobasalA/B27 growth medium (Invitrogen) supplemented with GlutaMax (0.25%, Invitrogen), glutamine (0.25-0.5 mM, Sigma), penicillin/streptomycin (1:100, ThermoFisher), and heat-inactivated fetal calf serum (10%, Sigma, CAS 1943609-65-1). Media was partially exchanged on day 3 (800 μL) and day 14 (500 μL) with fresh maintenance medium comprised of BrainPhys (StemCell), B27 (2%, Invitrogen), GlutaMax (0.25%, Invitrogen), and penicillin/streptomycin (1%, ThermoFisher). Cultures were maintained for up to 2-3 weeks at 37°C and 5% CO_2_ until use.

### Electrophysiology

Postsynaptic patch-clamp recordings were performed at mouse cortical neurons [days *in vitro* (DIV) 15-16] using a HEKA EPC10 amplifier (HEKA Elektronik). Pipette solution for voltage-clamp recordings contained 120 mM CsCl, 20 mM TEA-Cl, 10 mM HEPES, 5 mM EGTA, 3 mM Mg-ATP, 0.3 mM Na-GTP, 5 mM Na-Phosphocreatin, and 3 mM QX314-Cl. The pH was adjusted with CsOH to 7.31 and osmolarity was adjusted with sucrose to 298 mOsm. Series resistance (Rs) was, on average, 9.5 ± 2.3 MΩ and was compensated to remaining Rs of 5.1 ± 0.8 MΩ. Pipettes were pulled from borosilicate glass (Science Products) with a DMZ Universal Electrode (Zeitz Instruments) with a resistance of 3-4 MΩ. Recordings were performed at room temperature. Postsynaptic currents and potentials were analyzed with the NeuroMatic plug-in (Version 3) for Igor Pro (WaveMetrics, Lake Oswego, OR, USA; Version 9). Obtained parameters were presented and statistically compared with Sigma Plot 11 (Systat Software).

Recordings of the holding current (I_hold_) and spontaneous inhibitory postsynaptic currents (sIPSCs) were performed at a holding potential of −60 mV in extracellular Tyrode’s salt solution containing 145 mM NaCl, 2.5 mM KCl, 1.2 mM MgCl_2_, 2 mM CaCl_2_, 10 mM HEPES, and 10 mM glucose. The pH was adjusted by NaOH to 7.4. To isolate GABA_A_R-mediated currents, the extracellular solution was supplemented by 10 µM 2,3-Dihydroxy-6-nitro-7-sulfamoyl-benzo[*f*]chinoxalin-2,3-dion (NBQX; Biotechne, CAS 479347-86-9) to block AMPA receptors and 50 µM (2*R*)-2-Amino-5-phosphonopentanoic acid (APV; Tocris, CAS 79055-68-8) to block *N*-methyl-D-aspartate receptor (NMDARs). Voltages were not corrected for a calculated liquid junction potential of 2 mV. After establishing a baseline recording period (before condition ∼3 min), 10 µM chlorophene was washed in through the perfusion system and I_hold_ was monitored. After four min, 50 µM picrotoxin (Sigma, CAS 124-87-8) and 20 µM SR95531 (Sigma, CAS 104104-50-9) were bath-applied to block GABA_A_R-mediated chloride conductance. To compare I_hold_ before and during the application of chlorophene and after application of the GABA_A_R blockers, the mean I_hold_ was calculated for (i) the before period (I_hold_ ctr), (ii) 2 min after application of chlorophene (I_hold_ +chloro), and (iii) 1 min after application of picrotoxin and SR95531 (I_hold_ +PTX/SR) for each cell. The main effect of drug perfusion was tested with repeated measures ANOVA followed by pairwise comparisons (Bonferroni t-test, n=6 neurons).

To evaluate the effect of chlorophene on NMDA-evoked currents, NMDA currents were pharmacologically isolated by supplementing the extracelluar solution with 50 µM picrotoxin to block GABA_A_ receptors and 10 µM NBQX to block AMPA receptors. After establishing a stable baseline recording period, NMDA (1 µM, 10 µM or 20 µM, Merck, CAS 6384-92-5) was washed in through the perfusion system and the change of I_hold_, representing a NMDA evoked current, was monitored. After reaching a stable NMDA evoked current (∼3 min) chlorophene was bath applied. In a subset of cells, 50 µM APV was applied at the end of the experiment to block NMDA evoked currents. To compare the effect of chlorophene and APV exposure on NMDA currents, I_hold_ was calculated (i) 3 min after NMDA application (I_hold_ +NMDA), (ii) 2 min after chlorophene application (I_hold_ +chloro) and (iii) 1 min after APV application (I_hold_ +APV). The effect of chlorophene on NMDA currents was compared by a paired t-test (I_hold_ +NMDA vs. I_hold_ +chloro) and for the subset of cells with APV application by repeated measures ANOVA followed by pairwise comparisons (Bonferroni t-test).

The pipette solution for current clamp recordings contained 115 mM KMeSO_3_, 5 mM NaCl, 10 mM HEPES, 0.1 mM EGTA, 4 mM Mg-ATP, 0.3 mM Na-GTP, 20 mM K_2_-Phosphocreatin. The pH was adjusted with KOH to 7.33 and osmolarity was adjusted with sucrose to 298 mOsm. Voltages were not corrected for a calculated liquid junction potential of 9.6 mV. To test the effect of chlorophene on basal neuronal properties and neuronal firing the internal chloride concentration was set to 5 mM to mimic physiological chloride gradients and hyperpolarizing GABA_A_ receptor-mediated conductance. To isolate the effect of chlorophene via GABA_A_ receptors, the extracelluar solution was supplemented by 10 µM NBQX and 50 µM APV to block AMPA and NMDA receptors. The built-in bridge compensation was adjusted to 100% of the series resistance determined in voltage-clamp. Passive and active neuronal properties of single neurons were measured before (control) and after chlorophene application, respectively. The input resistance (R_in_) was calculated from alternating subthreshold current injections from −50 to 20 pA (10 pA steps, 200 ms duration). Action potential firing was evoked by suprathreshold depolarizing current injections. The current threshold required for eliciting an action potential was denoted as rheobase. The action potential frequency was calculated from the number of action potentials in response to a 200 ms current injection with an amplitude twice as large as the rheobase. The resting membrane potential (V_rest_) was measured ∼1 min after obtaining whole-cell access as the membrane voltage with zero current in current clamp mode. For statistical comparison of neuronal properties under control and chlorophene condition, p values were calculated by paired t-tests.

### Human multi-neurotransmitter receptor (hMNR) assay

The human multi-neurotransmitter (hMNR) assay used 3D BrainSpheres derived from human-induced pluripotent stem cell (hiPSC)-based neural progenitor cells (hiNPCs). The BrainSpheres were generated from the hiPSC line iPS(IMR90)-4, derived from IMR90 fibroblasts and reprogrammed using lentiviral transfection of OCT4, SOX2, NANOG, and LIN28 (WiCell, WISCi004-B)^70^. The cultivation of the hiPSC line IMR90 (clone 4, WiCell, Madison, WI, USA), including neural induction, was carried out according to a previously described 2D-NIM protocol^71^. Briefly, hiPSC colonies were singularized and cultured in defined medium, including dual SMAD inhibitors SB-431542 and LDN-193189 for 21 d in 2D. On day 21 of neural induction, hiNPCs were cryopreserved in single-cell suspension. For sphere generation, hiNPCs were thawed, transferred to 6-well plates, and cultivated in a gyrical shaking incubator (140 rpm, 12.5 mm diameter) in a proliferation medium^71^. To start the 3D neural differentiation, hiNPC spheres (0.3 mm in diameter) were cultivated in CINDA+ differentiation medium^71,72^ for 3 weeks, and half of the medium was replaced twice weekly. After 3D differentiation, BrainSpheres were plated on pre-coated (Poly-L-Ornithine, Laminin LN521) 96-well microelectrode arrays (MEA, #M768-tMEA-96B, Axion Biosystems) in CINDA+ differentiation medium (1 sphere per well) and fed twice per week by replacing half of the medium. After 4 weeks of differentiation on the MEAs, spontaneous electrical baseline activity was recorded with the Axion Maestro Pro MEA system. For initial unit identification, BrainSpheres were acutely exposed to the respective neurotransmitter agonists including glutamate (50 µM), and γ-aminobutyric acid (GABA; 10 µM). After electrical activity was measured, neurotransmitter antagonists were added to each well (10 µM bicuculline, 50 µM/50 µM D-APV/NBQX), network activity was recorded and substances were removed with a complete exchange of the medium. After a 3 h washout, the second baseline was recorded and chlorophene was gradually added at sequentially increasing concentrations (0.25, 0.74, 2.22, 6.67, 20 µM). Extracellular recording of electrical activity was performed for 15 min at 37°C and 5% CO_2_ in the Axion Maestro Pro System (Axion Biosystems), in line with previous work^71^. For further analysis of detected spikes, recorded .spk files were concatenated in the order of measurement and converted into a single .nex file with a MATLAB (R2021b, R2022b, MathWorks, Natick, MA, USA) script. Spike sorting of the .nex file was performed using the Offline Sorter (OFS, version 4.4, Plexon, Dallas, TX, USA) software, applying the automatic clustering T-Distribution EM method with 10 degrees of freedom (DOF) and an initial number of units of 20. The sorted units were exported as per-unit and per-waveform data. BrainSphere spiking data were analyzed using R scripts for statistical analysis and data visualization with the ggplot2 package. Only recordings with a spike count of ≥10 per 15 min in both the initial (recording 1) and the second baseline recording (recording 4) were included in the analysis. To determine neuronal subtype-specific responses, units were classified as either glutamatergic or GABAergic based on their spiking activity under neurotransmitter and receptor antagonist treatments. A unit was classified as glutamatergic (glutamate + APV/NBQX condition) if spiking increased in recording 2 (glutamate treatment) relative to recording 1 (baseline) and decreased in recording 3 (APV/NBQX treatment) relative to recording 2. GABAergic units (GABA + bicuculline condition) were identified by a decrease in spiking in recording 2 (GABA treatment) relative to recording 1 (baseline) and an increase in recording 3 (bicuculline treatment) relative to recording 2. For both neuronal subtypes, chlorophene exposure conditions were mapped to corresponding recording segments (recordings 4-9) with assigned concentrations ranging from 0-20 µM. To account for variability in baseline spiking activity across units, spike counts were normalized to the total spike count per unit across all analyzed recording segments, ensuring comparability across conditions. Wilcoxon signed-rank tests for paired data were performed to compare normalized spike counts at each chlorophene concentration against baseline (0 µM) within the same unit. Benjamini-Hochberg correction was applied to adjust for multiple comparisons.

### Availability of data and code

All data (DOI: 10.6084/m9.figshare.26819890) and code (DOI: 10.6084/m9.figshare.26819914) were uploaded to FigShare and will be made public upon paper acceptance.

## Results

### A multi-behavioral phenotyping approach to quantify visual and acoustic behaviors in larval zebrafish

To identify chemicals that modulate learning and memory in larval zebrafish, we devised a battery of 11 automated, sequential behavioral assays called the Visual and Acoustic Motor Response (VAMR) assay battery. The battery included assays which quantified baseline (BSL) activity, visual startle responses (VSR1,2), visual motor responses (VMR1-3), acoustic startle responses (ASR1,2), acoustic startle habituation (ASH1), potentiation of habituation (ASH1/ASH5), and memory retention (ASR3/ASR2) (**Fig. 1A**). Overall, four stimulus modalities were used. This included visual light at intensities of 0 lux (dark condition) and 7.5 klux (light condition). Additionally, 5 dpf larvae were exposed to acoustic stimuli at 300 Hz, with low (65 dB) and high (75 dB) volumes (**Fig. S1**). These stimuli provoked stereotyped motor activity patterns in larval zebrafish that were quantified as pixel change per second for each individual per well (**Fig. 1A**). First, in the BSL assay, zebrafish showed low average baseline activity per second. In the VSR assays, transitions from dark-to-light (**Fig. 1B**) or light-to-dark (**Fig. 1D**) conditions elicited rapid motor responses lasting 1 s. During the VMR assays, visual light caused low basal activity (**Fig. 1C**), whereas a lack of visual light gradually increased motor activity to a maximum after approximately 4 min which was followed by an asymptotic decrease to baseline (**Fig. 1E**). In the ASR1 and ASR2 assays, five low-or high-volume acoustic stimuli, with an inter-stimulus interval of 1 min, evoked weak or strong motor responses, respectively (**Fig. 1F,G**). Next, a series of 30 high-intensity acoustic stimuli with an inter-stimulus interval of 1 s induced robust habituation (**Fig. 1H,I**). Repetitive application of habituation trials with an inter-bout interval of 1 min gradually decreased motor activity (**Fig. 1J**), consistent with potentiation of habituation. Following a 3 min rest period, five high-volume acoustic stimuli elicited motor responses significantly lower than in the ASR2 assay (**Fig. 1K**), consistent with memory retention. Additional data (**Fig. S2**) show that the phenotyping battery meets key criteria for non-associative habituation learning. These findings support the potential of the VAMR battery to identify compounds that influence visual and acoustic behaviors in larval zebrafish, including those related to learning and memory.

**Figure 1.**
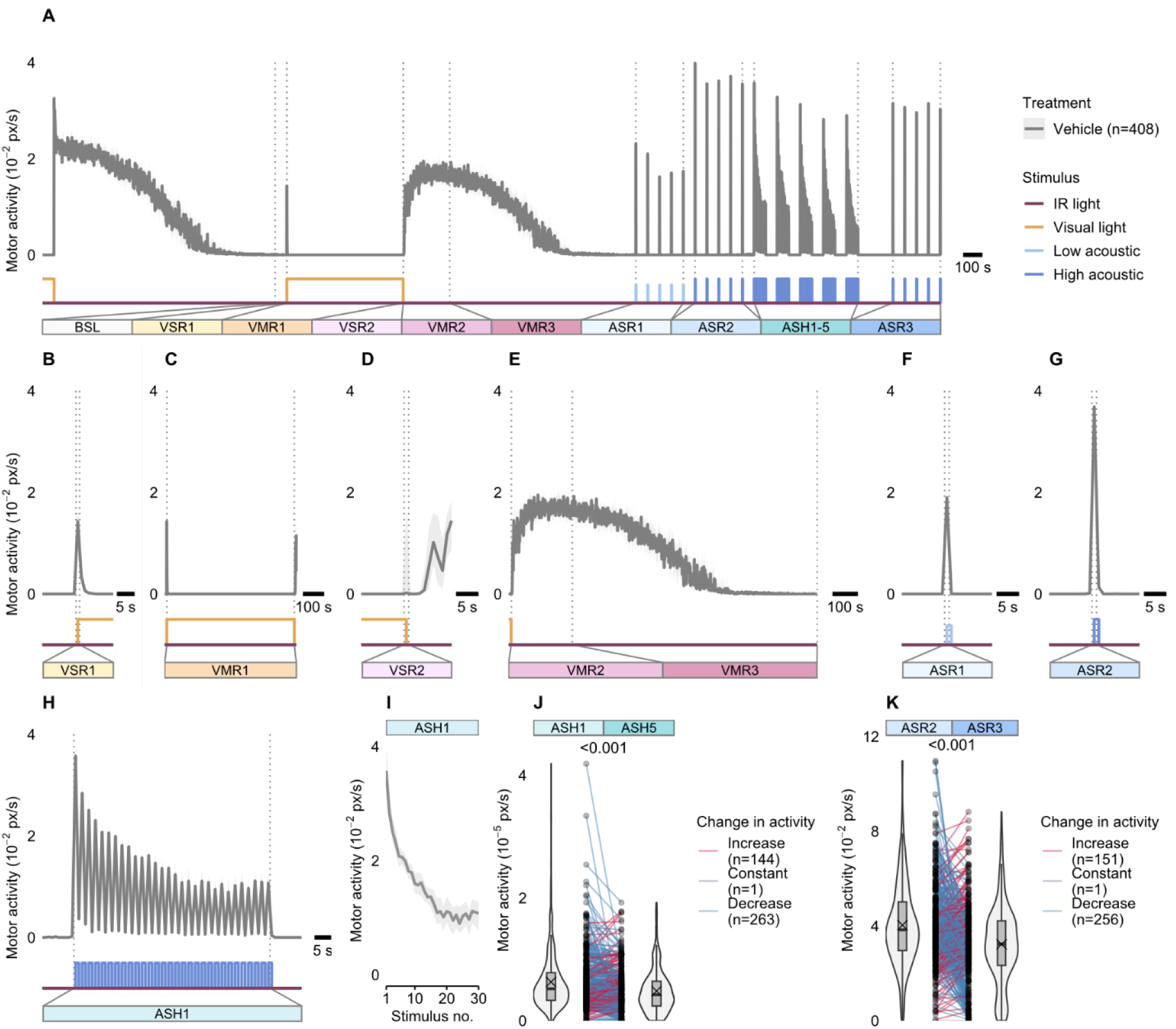
A battery of visual and acoustic assays and resulting behaviors in larval zebrafish. (**A**) Motor activity (y-axis) of zebrafish larvae over time and across various stimulus modalities (x-axis). Data are median ± 95% CI for *n* = 408 larvae obtained from 28 x 96-well plates, each containing 12-16 control individuals. (**B**) A rapid dark-to-light transition caused a visual startle response (VSR1). (**C**) Subsequent exposure to visual light caused low motor activity in zebrafish larvae (VMR1). (**D**) Light-to-dark transition caused an additional visual startle response (VSR2). (**E**) Motor activity upon light-to-dark transition reached a maximum after approximately 4 min (VMR2) followed by an asymptotic decrease to baseline (VMR3). (**F**) A series of five low- and (**G**) high-intensity acoustic stimuli (1 s) provokes acoustic startle responses with lower or higher amplitude, respectively. (**H**) A sequence of 30 high-intensity acoustic stimuli (inter stimulus interval, ISI = 1 s) induced an exponential response decrement, indicating habituation. (**I**) Representative plot of data used for **H**, excluding ISIs. (**J**) Decrease in responsiveness during first (ASH1), relative to fifth habituation bout (ASH5), indicates potentiation of habituation. (**K**) Responsiveness to non-habituating high-intensity acoustic stimuli is significantly decreased comparing pre-(ASR2) and post-habituation (ASR3) assays, indicating memory retention. Numbers above the split rainclouds in **J** and **K** represent p-values (one-sided Wilcoxon signed-rank test). *n* = 408. Summary data can be found in Excel Tables S1-S5. ASH, acoustic startle habituation; ASR, acoustic startle response; BSL, baseline motor activity; VMR, visual motor response; VSR, visual startle response.

### Multi-behavioral phenotyping of known modulators of habituation learning in larval zebrafish

To evaluate the assay battery, we first tested a reference set of compounds that act through mechanisms previously reported to modulate habituation learning in zebrafish^44,45^ (**Table S2**). Selected compounds included those reported to reduce short-term habituation including two *N*-methyl-D-aspartate receptor (NMDAR) antagonists ((2*R*)-2-Amino-5-phosphonopentanoic acid (APV), MK-801) and donepezil (acetylcholinesterase inhibitor) and compounds that reportedly enhance short-term habituation including citalopram (selective serotonin reuptake inhibitor), haloperidol (dopamine D2 receptor antagonist), picrotoxin (gamma-aminobutyric acid A receptor (GABA_A_R) antagonist), and yohimbine (α-2 adrenoreceptor antagonist). Larval zebrafish were exposed to a series of 7-10 concentrations per compound for 1 h before quantification of motor activity. To quantify habituation behavior, the initial habituation bout (ASH1) was used to calculate a ‘habituation score,’ achieved by dividing the motor activity during the final ten taps by the cumulative responses during the initial and final ten acoustic stimuli. On average, the untreated vehicle control exhibited a habituation score of 0.32. Compounds that reduced short-term habituation therefore generated values greater than the control group. In contrast, compounds that accelerated the habituation response yielded values lower than the control. In line with previous reports, the prototypical NMDAR antagonists APV and MK-801 significantly reduced non-associative habituation learning in a concentration-dependent manner (**Fig. 2A,D**), whereas citalopram (21.5, 100 µM) and haloperidol (1, 4.6 µM) significantly enhanced habituation (**Fig. 2B,C**). In addition, MK-801 significantly enhanced habituation at the highest test concentration (**Fig. 2D**). Non-significant alterations in habituation were determined in zebrafish exposed to picrotoxin, donepezil, or yohimbine (**Fig. S3**). These results demonstrate concentration-dependent effects of NMDAR antagonists and other compounds on habituation behavior in zebrafish.

**Figure 2.**
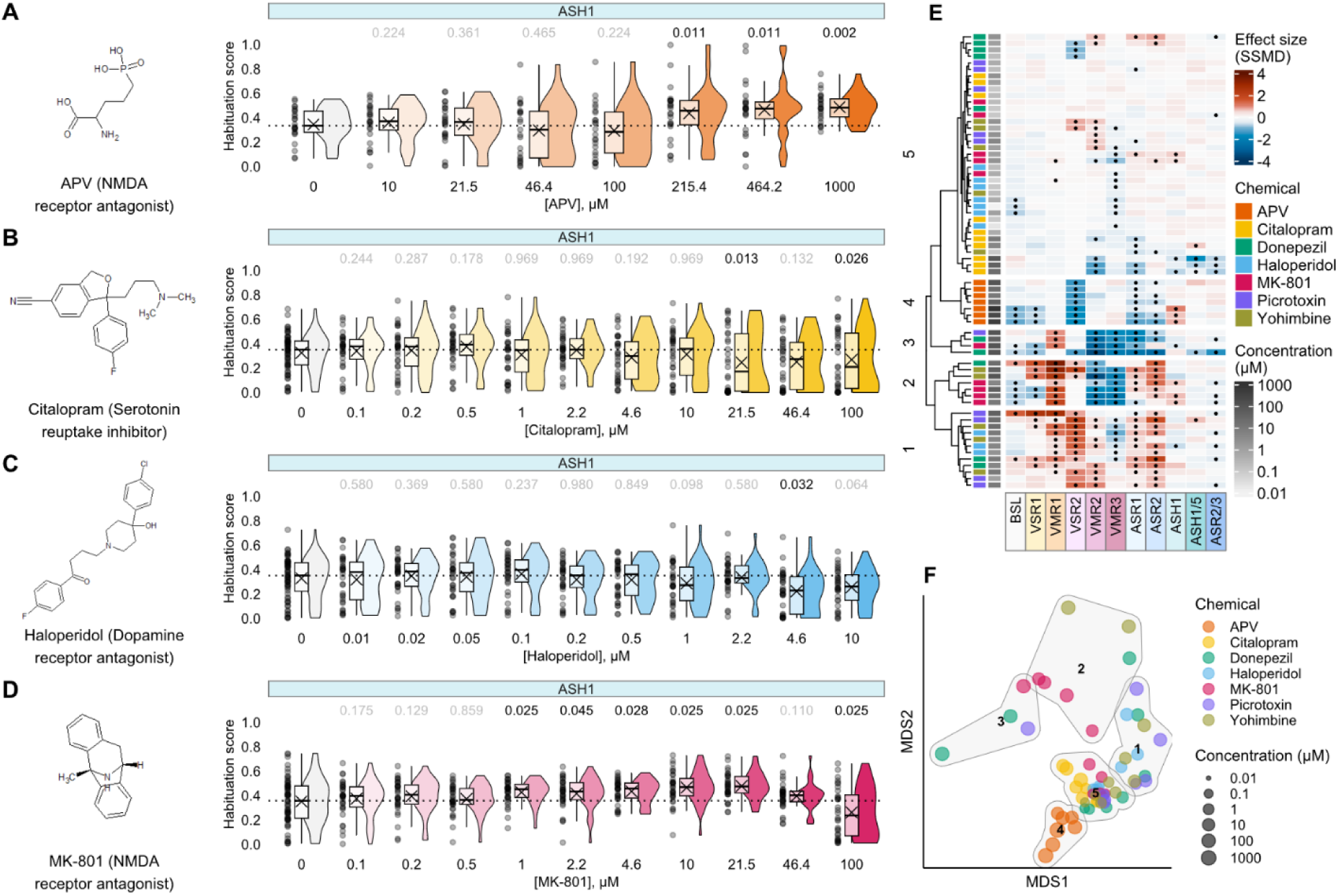
Behavioral phenotypes following acute exposure to known modulators of habituation learning. (**A-D**) Structures, modes of action, and habituation scores of reference modulators of habituation. Raincloud plots quantify acoustic startle habituation as habituation scores (y-axis) across indicated compounds and concentrations (x-axis). Horizontal dotted lines indicate the median habituation score of untreated larvae. Numbers above the rainclouds represent adjusted p-values (grey: p ≥ 0.05, black: p < 0.05, two-sample bootstrapping test, *n*_control_ = 24-64, *n*_treatment_ = 24-32). Habituation scores significantly greater than negative control indicate reduced habituation whereas reduced values reflect enhanced habituation. (**E**) Hierarchical clustering of compound- and concentration-specific profiles (y-axis), across the entire assay battery (**Fig. S4**), based on effect sizes (SSMD, strictly standardized median difference) across behavioral endpoints (x-axis). Heatmap colors indicate deviation from the median control phenotype where red and blue reflect higher and lower activity, respectively. Statistically significant changes in behavioral phenotypes are marked with a dot (p<0.05, two-sample bootstrapping test). (**F**) Multidimensional scaling (MDS) representation of pairwise Euclidean distances between behavioral profiles based on data shown in **E**. Contour labels (grey) and numbers indicate clusters shown in **E**. Summary data can be found in Excel Tables S6-S11. APV, (2*R*)-2-Amino-5-phosphonopentanoic acid; ASH, acoustic startle habituation; ASR, acoustic startle response; BSL, baseline motor activity; VMR, visual motor response; VSR, visual startle response.

To determine whether the reference compounds selectively modulated short-term habituation learning, we simplified motor activity patterns by condensing compound- and concentration-dependent responses into behavioral profiles reflecting effect size relative to vehicle control (**Fig. 2E, Fig. S4**). Except for habituation (described above), habituation potentiation, and memory retention endpoints, the cumulative motor activity per assay for each individual was calculated. The ‘potentiation score’ (ASH1/5) reflects the ratio of the scaled sum of motor responses during taps in the final habituation bout (ASH5) to the sum of responses in the initial (ASH1) and final (ASH5) habituation bouts. Motor activity in the ASR3 assay, relative to an individual’s overall responses summed across ASR2 and ASR3 assays, was used to quantify memory retention (ASR2/3). Subsequently, we determined effect sizes relative to vehicle control for each of the eleven endpoints, yielding compound- and concentration-specific behavioral profiles. To determine phenotypic similarities between behavioral profiles, we used two complementary approaches, hierarchical clustering (**Fig. 2E**) and multidimensional scaling (MDS; **Fig. 2F**). While the majority of compounds caused slight behavioral changes in the lower concentration range (cluster 5), exposure to higher concentrations provoked diverse multi-behavioral phenotypes (clusters 1-4) (**Fig. 2E**). Within these clusters, chemical exposure caused different combinations of behavioral alterations, such as enhanced visual and acoustic startle responses (cluster 1), hyper- and hypolocomotion in response to light or dark conditions (cluster 2), or reduced visual and acoustic startle activity (clusters 3 and 4) (**Fig. 2E**). Behavioral profiles of the NMDAR antagonist APV (cluster 4) were segregated from all other profiles including concentrations that significantly reduced habituation (**Fig. 2E,F**). While both NMDAR antagonists (APV and MK-801) blocked habituation learning and reduced dark-to-light visual startle responses in zebrafish, disparate phenotypes were also observed. Specific to APV, exposure also reduced all other visual and acoustic startle responses (**Fig. 2E, Fig. S4**). In contrast, MK-801 exposure enhanced acoustic startle responses, caused light-sensitive hyperlocomotion, and resulted in lower responsiveness under dark conditions (**Fig. 2E, Fig. S4**). At concentrations that resulted in lower habituation, behavioral profiles of MK-801 differed from APV and clustered with profiles of picrotoxin (cluster 3), donepezil (clusters 2 and 3), and yohimbine (cluster 2) (**Fig. 2E,F**). *In silico* target predictions (**Table S7**) indicated that MK-801 may interact with GABA_A_ and/or acetylcholine receptors in zebrafish. In summary, while the assay battery shows that known habituation modulators do not act exclusively on habituation behavior, it enables the identification and differentiation of chemical modes of action.

### Targeted screen for chemical compounds that affect learning and memory

Based on these results, we hypothesized that the assay battery could be used to identify chemical compounds that affect learning and memory in zebrafish. Given the well-characterized involvement of NMDARs in synaptic plasticity and memory function^73–75^, we conducted a targeted selection of candidate compounds. The EPA CompTox Chemicals Dashboard, which contains bioactivity data on thousands of chemicals tested across numerous high-throughput screening assays^76^, was mined for relevant assays. We identified two tissue-based, cell-free radioligand binding assays that used extracted proteins from rat forebrain membranes to determine changes in NMDAR agonist activity (**Fig. 3A**, **Table S4**). In total, these datasets contained 81 unique chemicals of which 26 were annotated as hits (**Table S5**). To enrich the set of hits for compounds that exhibited higher specificity towards NMDARs relative to non-specific general toxicity, we used *in vitro* cytotoxicity and baseline toxicity data (**Table S6**). These data were used to rank order compounds according to their specificity for NMDAR interaction relative to non-specific toxicity effects (**Fig. 3A-C**). Rankings predicted 11 compounds with higher potency for NMDAR modulation relative to cytotoxicity and baseline toxicity levels (**Fig. 3B,C**). For example, besonprodil and MK-968, two preclinical drugs that act as NMDAR antagonists, and methadone, a µ-opioid receptor antagonist used as a painkiller, were among the predicted hits that appeared in both rankings (**Fig. 3B,C**). Ten compounds were purchased and screened in our behavior assay battery. In addition, we selected two chemicals (cypermethrin, ascorbic acid) as negative reference compounds for NMDAR modulation. Cypermethrin is a pyrethroid insecticide that prolongs the opening of sodium channels^77^. From a structural perspective, it is closely related to deltamethrin, a developmental neurotoxicant^78^. In contrast, ascorbic acid was proposed as a negative chemical for developmental neurotoxicity^21^.

**Figure 3.**
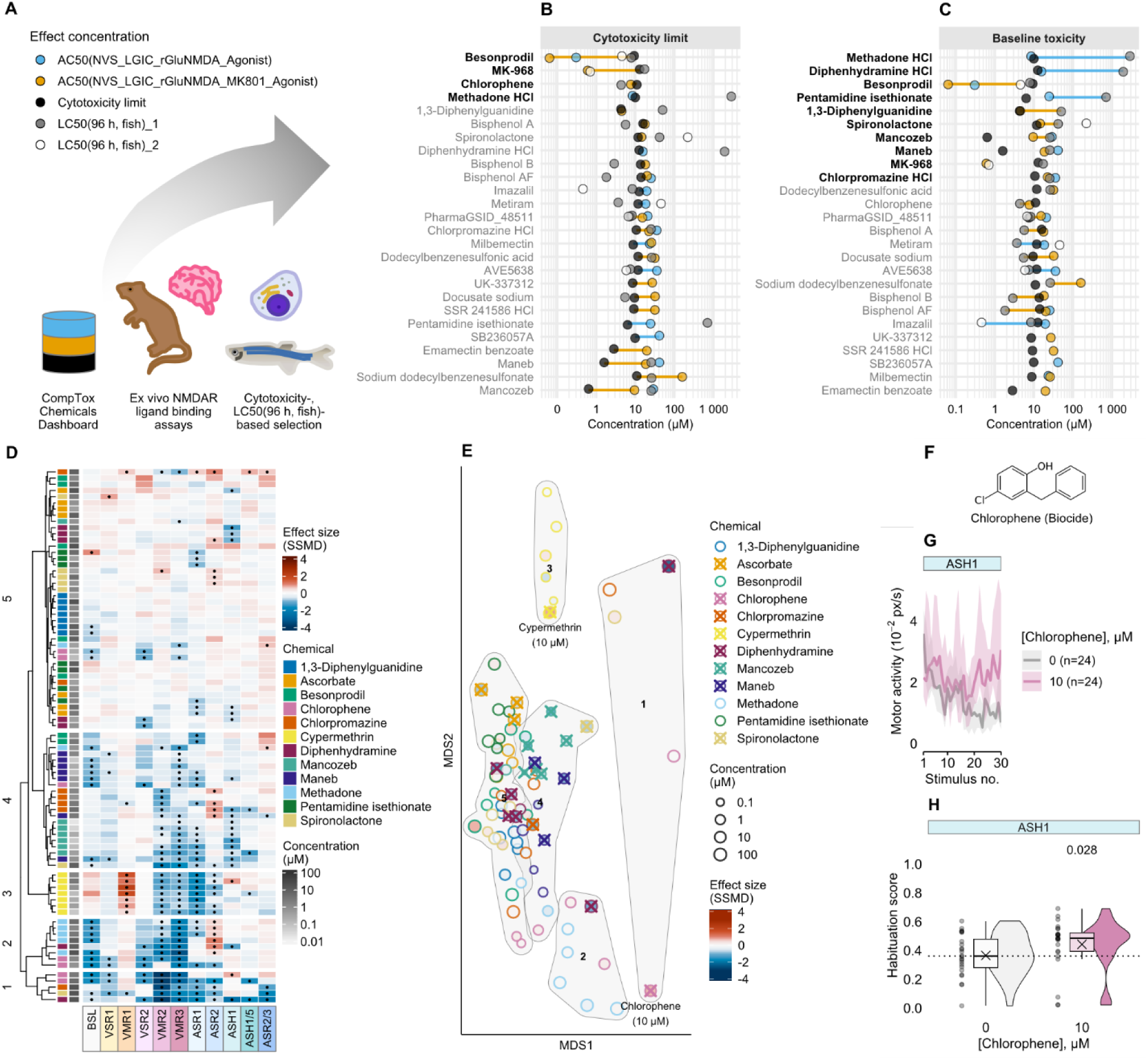
Prediction and evaluation of chemical compounds that interfere with *N*-methyl-D-aspartate receptor (NMDAR)-dependent signaling. (**A**) Data on *ex vivo* rat NMDAR radioligand binding (AC_50_: blue, orange), *in vitro* cytotoxicity limit (lower bound on the estimate of the cytotoxicity threshold; black), and predicted baseline toxicity in fish (LC_50_; grey, white) were gathered from the US EPA CompTox Chemical Dashboard and ECOSAR. (**B**) Cytotoxicity- and (**C**) baseline toxicity-based rankings of 26 *ex vivo* positive chemicals (y-axis) predict 11 unique candidate compounds (bold black) that alter NMDAR agonist activity with higher potency relative to cytotoxicity and predicted baseline toxicity levels. (**D**) Hierarchical clustering of behavioral profiles for 10 predicted candidate compounds and two negative reference chemicals (ascorbate, cypermethrin) (y-axis) across behavioral endpoints (x-axis). Heatmap colors based on effect sizes (SSMD, strictly standardized median difference) indicate deviation from the median control phenotype. Red and blue reflect higher and lower activity, respectively (*n* = 12-24 larvae per condition). Statistically significant changes in behavioral phenotypes are marked with a dot (p<0.05, two-sample bootstrapping test). (**E**) Multidimensional scaling (MDS) representation of pairwise distances between behavioral profiles based on data shown in **D**. Fill color indicates deviation from the median control phenotype in the acoustic startle habituation (ASH1) assay where red and blue reflect reduced and enhanced habituation, respectively. Contour labels (grey) and numbers indicate clusters shown in **D**. Crosses mark statistically significant changes in the acoustic startle response (ASH1) assay (p<0.05, two-sample bootstrapping test). (**F**) Structure of the positive hit chlorophene. (**G**) Median motor activity (y-axis) of vehicle control (0.01% DMSO, grey) and chlorophene-exposed (10 µM, pink) zebrafish larvae over 30 sequential (inter stimulus interval = 1 s) high-intensity acoustic stimuli (x-axis) (shaded envelope represents the 95% confidence interval). (**H**) Raincloud plot showing the habituation score (y-axis) for zebrafish exposed to 0.01% DMSO (vehicle control; grey) or chlorophene (10 µM, pink) (horizontal dotted line; median control habituation score). The number above the raincloud represents the adjusted p-value (two-sample bootstrapping test). Summary data can be found in Excel Tables S12-S17. ASH, acoustic startle habituation; ASR, acoustic startle response; BSL, baseline motor activity; VMR, visual motor response; VSR, visual startle response.

To determine whether exposure to the selected compounds affected behavior, we treated zebrafish for 1 h with multiple concentrations of each test chemical and translated the 11 quantitative features into behavioral profiles (**Fig. S5**). Hierarchical clustering segregated compound- and concentration-dependent profiles into five main clusters (**Fig. 3D**). Cluster 1 was primarily characterized by hypoactivity across the majority of assays and included compounds such as chlorophene (biocide) and spironolactone (diuretic) (**Fig. 3D**). A second cluster (cluster 2) exhibited lower baseline and dark-phase motor activity (VMR2, VMR3) while often displaying enhanced responsiveness to high-volume acoustic stimuli (ASR2) (**Fig. 3D**). This cluster included methadone, diphenhydramine (antihistamine), and chlorophene. Cluster 3 exclusively contained cypermethrin, which provoked light-sensitive hyperlocomotion (VMR1) in a concentration-dependent manner and reduced motor activity across multiple visual- and acoustic-based endpoints (**Fig. 3D**). In cluster 4, compounds were associated with reduced dark-phase motor activity (VMR2, VMR3), lower responsiveness to low-intensity acoustic stimuli (ASR1), and enhanced acoustic startle habituation (ASH1) (**Fig. 3D**). This group contained chlorpromazine (neuroepileptic) as well as the structurally related fungicides mancozeb and maneb. Cluster 5 displayed relatively subtle behavioral changes. Notably, this group included several compounds that were exclusively represented in this cluster, such as 1,3-diphenylguanidine (vulcanization accelerator), ascorbate (antioxidant), and the antibiotic pentamidine isethionate (**Fig. 3D**). To visualize chemical modulators of habituation learning among the variety of phenotypes, we projected all multi-behavioral profiles into two dimensions using MDS and shaded each data point with the effect size determined in the ASH1 assay (**Fig. 3E**). We found that behavioral profiles with enhanced or reduced startle habituation spread across the two-dimensional space, reflecting diverse multi-behavioral phenotypes (**Fig. 3E**). Chlorpromazine (21.5 µM), diphenhydramine (1.4 µM, 4.6-100 µM), mancozeb (0.02-1 µM), maneb (1-2.2 µM, 10 µM), and spironolactone (46.4 µM) significantly enhanced acoustic startle habituation (**Fig. 3D,E**, **Fig. S6**, p<0.05, two-sample bootstrapping test). In contrast, besonprodil (100 µM, cluster 2), cypermethrin (10 µM, cluster 3), and chlorophene (10 µM, cluster 4), reduced habituation (**Fig. 3D,E**). Cypermethrin (p = 0.018) and chlorophene (p = 0.028) both caused statistically significant reductions in habituation (**Fig. 3F-H**, **Fig. S6**, p<0.05, two-sample bootstrapping test). Overall, six out of ten of the predicted hits were active in our ASH1 assay. Notably, both negative reference chemicals, cypermethrin and ascorbate, showed unexpected activity. While ascorbate (10-21.5 µM, 100 µM) significantly enhanced acoustic startle habituation, cypermethrin (10 µM) significantly reduced it (**Fig. S6**). However, cypermethrin’s effect is likely a false positive, as it severely impaired zebrafish responsiveness to acoustic stimuli, leading to an artifact in the habituation score calculation (**Fig. S7**). These findings indicate that the VAMR assay battery may help identify compounds with potential effects on NMDAR-mediated signaling. The zebrafish assay may provide insights into diverse phenotypic outcomes that could be further investigated mechanistically.

### Evaluation of NMDARs and glutamatergic signaling as putative target sites of chlorophene

Given its wide use as a biocide, preservative, and disinfectant^79,80^, we investigated chlorophene in more detail. Because chlorophene exposure impaired habituation in zebrafish (**Fig. 3G,H**) and altered NMDAR agonist activity in rat *ex vivo* preparations (**Fig. 3B**), we explored whether chlorophene might influence habituation learning through effects on NMDARs. We further examined whether the underlying molecular mechanisms might be conserved across taxa, including mice and humans. To evaluate the translational relevance of these findings, we used whole-cell patch-clamp recordings in cultured mouse cortical neurons (**Fig. 4A**). We found that, while the NMDAR antagonist APV effectively inhibited the current response provoked by NMDA application, chlorophene application failed to do so (**Fig. 4B-D**). As a second approach to determine whether chlorophene targeted glutamatergic signaling, the compound was evaluated in a microelectrode array (MEA) assay using human-induced pluripotent stem cell-derived (hiPSC) 3D BrainSpheres. In this *in vitro* model, chlorophene exposure reduced the normalized spiking count of glutamatergic units in a concentration-dependent manner (**Fig. 4E**). Together, these findings indicate that while chlorophene might influence glutamatergic signaling, they do not provide evidence for direct NMDAR antagonism as the mechanism underlying reduced habituation in zebrafish.

**Figure 4.**
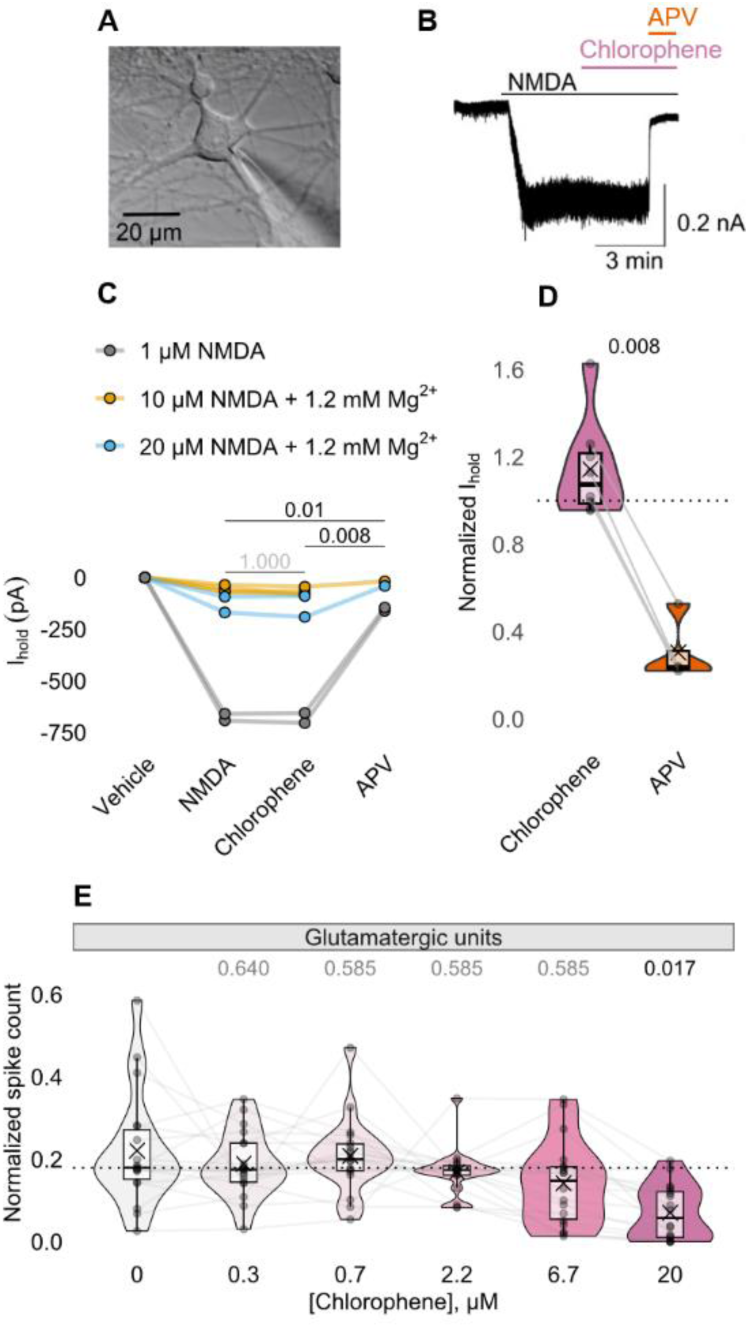
Interaction of chlorophene with *N*-methyl-D-aspartate receptors (NMDARs) in cultured mouse cortical neurons and effect of chlorophene exposure on the normalized spike count of glutamatergic units in human stem cell-derived 3D BrainSpheres. (**A**) Representative infrared differential interference contrast image of a patch-clamped cultured mouse cortical neuron. (**B**) Representative current response to NMDA (1 µM) remained unchanged during chlorophene (10 µM) application, but was inhibited by the NMDAR antagonist (2*R*)-2-Amino-5-phosphonopentanoic acid (APV; 50 µM). (**C**) Individual holding currents (I_hold_) before (vehicle) and during NMDA administration were not affected by chlorophene (10 µM, *n* = 8), but inhibited by APV (50 µM, *n* = 4). (**D**) I_hold_ during chlorophene (10 µM, *n* = 8) and APV (50 µM, *n* = 4) application normalized to I_hold_ evoked by respective NMDA exposure. (**E**) Normalized spike count (y-axis) of human stem cell-derived 3D-differentiated BrainSpheres exposed to increasing concentrations of chlorophene (x-axis). After the first baseline recording, the neurotransmitter glutamate, followed by the antagonists APV/NBQX (2,3-dihydroxy-6-nitro-7-sulfamoyl-benzo[*f*]quinoxaline), were applied to identify glutamatergic units (paired Wilcoxon signed-rank test, *n* = 18). Summary data can be found in Excel Tables S18-S20.

### Orthogonal evaluation of GABA_A_Rs as putative target sites of chlorophene in zebrafish, mouse, and human models

To determine whether chlorophene acted on targets other than NMDARs, we used the Similarity Ensemble Approach^68^ to predict protein targets based on structural similarity to their ligands. Among 107 predicted targets were three GABA_A_R subunits including α2, α3, and α5 (**Table S8**). Certain agonists and positive allosteric modulators of GABA_A_Rs have been reported to cause sedation and paradoxical excitation in zebrafish^49^, two phenotypes observed in chlorophene-exposed larvae (**Fig. 3G,H and Fig. 5A,B**). We therefore hypothesized that sedation in visually-mediated responses and enhanced responsiveness to repeated acoustic stimuli, were related to a GABAergic mode of action in chlorophene-exposed zebrafish. To determine whether chlorophene exposure caused these phenomena via GABA_A_R agonism, we pretreated zebrafish with the GABA_A_R antagonist picrotoxin (**Fig. 5C**). Pharmacological intervention rescued chlorophene-induced sedation, restoring visual motor responses (**Fig. 5C**, **Fig. S8**) whereas picrotoxin pretreatment did not counteract the habituation deficit caused by chlorophene exposure (**Fig. 5D**). Together, these observations indicate that chlorophene exposure was associated with sedation and paradoxical excitation, but not defective habituation, through GABA_A_Rs in zebrafish.

**Figure 5.**
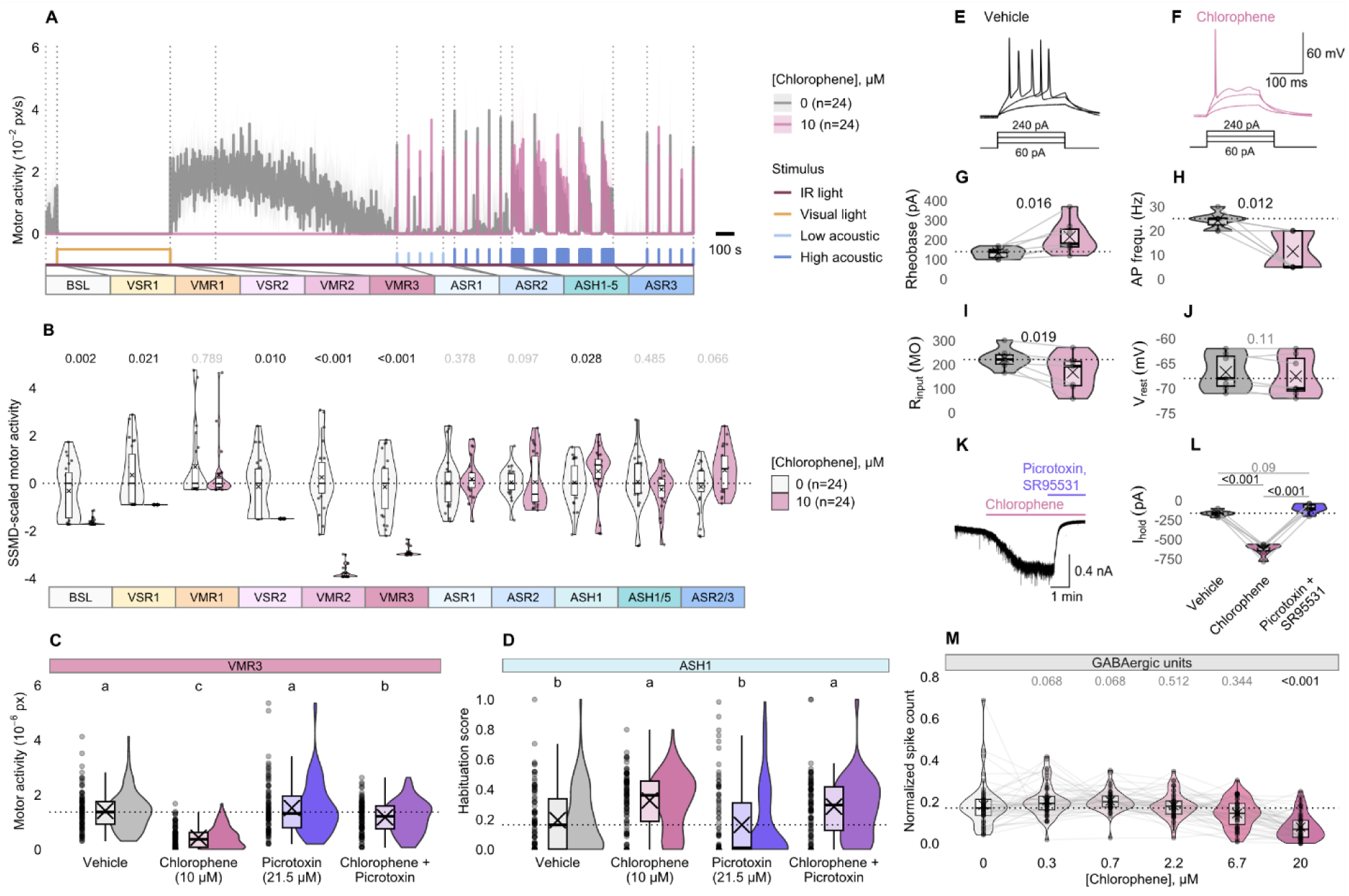
Effects of chlorophene exposure on zebrafish behavior, and GABAergic signaling in cultured mouse cortical neurons and human stem cell-derived 3D BrainSpheres. (**A**) Motor activity (y-axis) of zebrafish treated with chlorophene (10 µM, red) or 0.01% DMSO (grey) over time and across various stimulus modalities (x-axis). Data represents median ± 95% CI. (**B**) Split raincloud plots show the effect of chlorophene exposure on zebrafish motor activity (y-axis) across 11 assay endpoints (x-axis). Motor activity is expressed as effect size (SSMD, strictly standardized median difference) to represent different assay endpoints on a common scale. Data points are equally shifted to set the median control activity per assay endpoint to zero. Numbers above the rainclouds represent adjusted p-values (grey: p ≥ 0.05, black: p < 0.05, two-sample bootstrapping test). (**C**) Representative raincloud plot showing the cumulative motor activity (y-axis) of zebrafish exposed to the indicated compounds (x-axis) in response to dark conditions (VMR3) (Kruskal-Wallis test, *n* = 94-96 larvae). (**D**) Habituation scores (y-axis) of zebrafish treated with the indicated compounds (x-axis). A sequence of 30 high-intensity acoustic stimuli with 1-s inter-stimulus interval (ISI) was used to induce acoustic startle habituation (ASH1) (Kruskal-Wallis test, *n* = 94-96 larvae). (**E**) Representative response of a cultured mouse cortical neuron to depolarizing current injections under the control condition (black) and (**F**) during chlorophene application (10 µM, pink). (**G-J**) Passive membrane and intrinsic properties of cultured mouse cortical neurons without and during application of chlorophene including (**G**) rheobase, (**H**) action potential (AP) firing frequency at 2x rheobase, (**I**) input resistance (R_input_), and (**J**) resting membrane potential (V_rest_) (paired t-tests, *n* = 7). (**K**) Representative current response of a cultured mouse cortical neuron to chlorophene (10 µM) was inhibited by co-exposure to the gamma-aminobutyric acid A receptor (GABA_A_R) antagonists picrotoxin (50 µM) and SR95531 (10 µM). (**L**) Individual and median holding current (I_hold_) under control condition, during chlorophene application and GABA_A_R blockade by picrotoxin and SR95531 (repeated measures ANOVA, Bonferroni t-test, *n* = 6). (**M**) Normalized spike count (y-axis) of 3D BrainSpheres treated with increasing concentrations of chlorophene (x-axis). After the first baseline recording, the neurotransmitter GABA followed by the antagonist bicuculline were applied to identify GABAergic units (two-sample bootstrapping test, *n* = 46). Summary data can be found in Excel Tables S21-S27. ASH, acoustic startle habituation; ASR, acoustic startle response; BSL, baseline motor activity; VMR, visual motor response; VSR, visual startle response.

We next sought to evaluate whether chlorophene acts as a GABA_A_R agonist in mouse- and human-based models. To determine whether chlorophene acts on GABA_A_Rs we first measured currents in cultured mouse cortical neurons. Chlorophene exposure induced a slow tonic chloride current that was blocked by the GABA_A_R antagonists picrotoxin and SR95531 (**Fig. 5K,L**). This suggests that chlorophene activates rodent GABA_A_Rs. To investigate the effect of GABA_A_R activation by chlorophene on neuronal action potential firing, we measured the membrane potential during current injections while blocking glutamatergic receptors with APV and 2,3-Dihydroxy-6-nitro-7-sulfamoyl-benzo[*f*]chinoxalin-2,3-dion (NBQX) (**Fig. 5E,F**). Chlorophene exposure increased the minimum current needed to generate an action potential (i.e. the rheobase; **Fig. 5G**), decreased the action potential firing frequency and the input resistance (**Fig. 5H and I**), and did not change the resting membrane potential (**Fig. 5J**). These data suggest that chlorophene-induced GABA_A_R-mediated chloride conductance shunts neuronal action potential spiking by lowering the input resistance and increasing the current threshold required for neuronal spiking. To determine whether chlorophene acts as a GABA_A_R modulator in humans, we used the 3D BrainSphere MEA assay where the normalized spike count of GABAergic units was significantly decreased in chlorophene-treated (20 µM) 3D BrainSpheres (**Fig. 5M**). In summary, these data show that chlorophene exposure was associated with similar effects on GABA_A_Rs in zebrafish, mouse, and human-based models.

### Evaluation of M-current activation as putative mechanism causing defective habituation in chlorophene-exposed zebrafish larvae

Certain anesthetic GABA_A_R ligands, such as etomidate and propofol, produce sedation and paradoxical excitation in zebrafish^49^. M-current (Kv7 potassium channel) ligands have been shown to modify etomidate-induced paradoxical excitation in zebrafish^49^. To test whether the M-current plays a role in regulating sedation, paradoxical excitation, and potentially habituation, we examined the effects of the M-current activator flupirtine^81,82^ in our battery of behavior assays (**Fig. S9**). We found that flupirtine (21.5-46.4 µM) impaired habituation in zebrafish (**Fig. 6A**). Additionally, flupirtine induced both sedation and paradoxical excitation (**Fig. 6B**). To determine phenotypic similarities and dissimilarities, we compared behavioral profiles of chemical compounds identified as impairing habituation including APV, chlorophene, cypermethrin, flupirtine, and MK-801, using hierarchical clustering and MDS. With regard to profiles associated with impaired habituation, hierarchical clustering revealed that flupirtine (21.5-46.4 µM) was phenotypically related to chlorophene (10 µM) and MK-801 (21.5-46.4 µM) (cluster 1, **Fig. 6C**). In contrast, cypermethrin (0.1-10 µM) formed a distinct, compound-specific cluster (cluster 2, **Fig. 6C**). APV (10-1000 µM) solely populated cluster 3, which also included chlorophene (0.2-4.6 µM, 21.5 µM) and flupirtine (1-10 µM) profiles that did not affect habituation, as well as lower concentrations of MK-801 (0.1-4.6 µM) (Fig. **6C**). MDS revealed that flupirtine (46.6 µM) and chlorophene (10 µM) clustered more closely together than with other profiles, indicating high phenotypic similarity (**Fig. 6D**). Together, these data show that M-current activation is associated with sedation, paradoxical excitation, and habituation deficits in zebrafish, supporting its potential involvement in chlorophene-induced effects.

**Figure 6.**
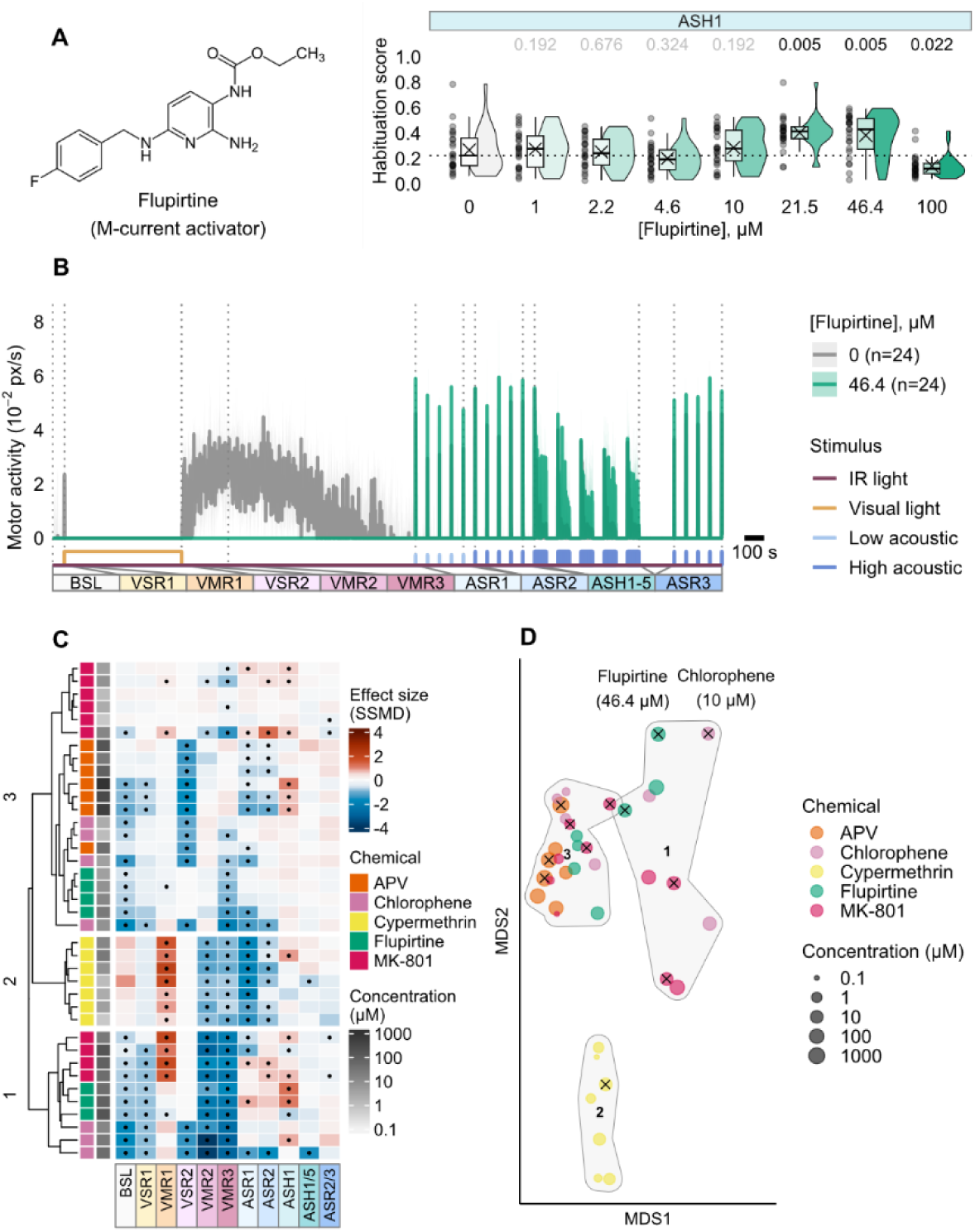
Effects of M-current activator flupirtine on zebrafish behavior and comparison with behavioral phenotypes of chemical compounds determined to reduce habituation in this study. (**A**) Structure of flupirtine and habituation scores following acute exposure. Raincloud plots quantify acoustic startle habituation as habituation scores (y-axis) across multiple flupirtine concentrations (x-axis). Horizontal dotted lines indicate the median habituation score of untreated larvae. Numbers above the rainclouds represent adjusted p-values (grey: p ≥ 0.05, black: p < 0.05, two-sample bootstrapping test, *n* = 24). Habituation scores significantly greater than negative control indicate reduced habituation whereas reduced values reflect enhanced habituation. (**B**) Motor activity (y-axis) of zebrafish treated with flupirtine (46.4 µM, green) or 0.01% DMSO (grey) over time and across various stimulus modalities (x-axis). Data represents median ± 95% CI. (**C**) Hierarchical clustering of behavioral profiles for the five chemical compounds identified to reduce habituation in this study (y-axis) across behavioral endpoints (x-axis). Heatmap colors based on effect sizes (SSMD, strictly standardized median difference) indicate deviation from the median control phenotype. Red and blue reflect higher and lower activity, respectively (*n* = 24-64 larvae per condition). Statistically significant changes in behavioral phenotypes are marked with a dot (p<0.05, two-sample bootstrapping test). (**D**) Multidimensional scaling (MDS) representation of pairwise Euclidian distances between behavioral profiles based on data shown in **C**. Contour labels (grey) and numbers indicate clusters shown in **C**. Crosses mark statistically significant reductions in habituation in the acoustic startle response (ASH1) assay (p<0.05, two-sample bootstrapping test). Summary data can be found in Excel Tables S28-S31. APV, (2*R*)-2-Amino-5-phosphonopentanoic acid; ASH, acoustic startle habituation; ASR, acoustic startle response; BSL, baseline motor activity; VMR, visual motor response; VSR, visual startle response.

## Discussion

Given the ubiquitous exposure to diverse environmental chemicals across lifespan and the lack of information on the hazards potentially presented by these chemical compounds, there is an unmet need to efficiently assess their potential adverse effects on neurodevelopment and function. Here we describe a multi-behavioral phenotyping approach in early life-stage zebrafish that enables (i) the rapid identification of neuroactive compounds and (ii) the elucidation of mechanisms underlying alterations in neurobehavior. While current behavior-based toxicity tests in larval zebrafish, such as the light-dark transition test^58,83–85^, largely focus on a small number of visual motor behaviors, we merged this widely used assay with additional, well-described paradigms, broadly applied in large-scale behavioral drug screens, including acoustic startle responses^46,49^, habituation learning^44,45,47^, and other memory-related endpoints that have yet to be applied to toxicity testing.

The Visual and Acoustic Motor Response (VAMR) battery builds upon established zebrafish behavioral assays by integrating multiple stimulus-driven responses to assess sensorimotor function, learning, and memory. While previous studies have developed various methods to analyze zebrafish behavior, many have either focused on specific behavioral paradigms^43–45^ or prioritized compound classification^48,57,86^, often without a strong emphasis on learning and memory.

Sensorimotor and learning-based paradigms have provided key insights into zebrafish behavior and cognition. Visual-motor responses have been widely studied as measures of sensorimotor adaptation. Burgess & Granato^43^ showed that zebrafish adjust locomotor activity in response to illumination changes, increasing movement following dark transitions and exhibiting sustained reductions under prolonged light exposure. Best et al.^45^ demonstrated non-associative learning in larval zebrafish through acoustic startle habituation, fulfilling key criteria such as spontaneous recovery and dishabituation. Wolman et al.^87^ extended this work by automating the assessment of short- and long-term habituation and conducting a high-throughput screen that identified novel cognitive modulators. Together, these foundational assays have been instrumental in characterizing both sensorimotor processing and experience-dependent behavioral modifications.

Building on these paradigms, multi-behavioral assay batteries have been developed to classify neuroactive compounds based on their effects across multiple behavioral domains. Bruni et al.^57^ introduced a battery of ten assays incorporating visual and acoustic stimuli of varying intensities and wavelengths, designed to generate detailed behavioral profiles for neuroactive compound classification. McCarroll et al.^49^ applied a subset of these assays to characterize central nervous system depressants, particularly focusing on motor suppression under excitatory light conditions and altered responses to low-intensity acoustic stimuli. Jordi et al.^47^ developed a high-throughput system to identify compounds that selectively influenced feeding behavior while minimizing off-target effects, combining feeding assays with locomotion, dark pulse responses, and habituation to mechanical stimuli.

Expanding on these approaches, the VAMR battery applies these rationales by evaluating the selectivity of habituation modulators while extending beyond more untargeted compound classification, where diverse stimuli are used without the intent to model specific phenotypes or rely on defined molecular mechanisms^57^. By integrating multiple stimulus-driven responses, including visual motor responses, visual and acoustic startle, habituation, and memory, within a structured framework, the VAMR battery enables a more targeted and mechanistically informed assessment of learning and memory, while also being applicable to other behavioral domains. This approach supports the evaluation of pharmacological agents as well as environmental chemicals.

This work focused on the identification of chemical modulators of habituation, which is a form of non-associative learning^88^. Defective habituation is associated with decrements in learning and memory^89,90^ and is prevalent in neuropsychiatric disorders such as autism spectrum disorder, schizophrenia, and attention deficit hyperactivity disorder^91^. Here, we demonstrated that the phenotyping battery fulfills hallmark criteria of non-associative learning^88^, a phenomenon conserved across phyla including zebrafish^45^, rodents^92^, and humans^93^. Confirmation of habituation characteristics (**Fig. S2**) and the reproducibility of phenotypes observed with known habituation modulators demonstrate that the VAMR battery quantifies non-associative learning. This includes a quantitative demonstration of dishabituation, which is defined as an increased response to a habituated stimulus following a second, brief stimulus using a different stimulus modality^88^. Dishabituation to acoustic stimuli has been quantitatively demonstrated 7 dpf zebrafish using a second, light-based stimulus^45^ while in 6 dpf larvae, dishabituation following a user-based tactile stimulus has been reported^44,94^. This study demonstrates dishabituation in 5 dpf zebrafish, using a light-based secondary stimulus (**Fig. S2**). While our findings confirm the reproducibility of habituation characteristics and the effects of certain known modulators, it is important to consider potential strain-specific variability. For instance, Best et al.^45^ found that while both AB and WIK strains exhibit habituation in the acoustic startle response (ASR) assay, WIK larvae display a stronger initial response and take longer to return to baseline, highlighting strain-dependent behavioral differences. Additionally, strain-dependent effects of chemical exposure have been widely reported^95–97^. Future studies should assess whether the key findings observed here are consistent across different, more commonly used wild-type strains to ensure broader applicability.

In line with previous studies, we found that acute administration of the prototypical NMDAR antagonists APV or MK-801 reduced acoustic startle habituation in zebrafish^44,54^. This observation is consistent with deficits in learning and memory reported in rodent-based test paradigms^98,99^. We found that different chemical modulators do not act with specificity to exclusively alter habituation. This may result from off-target activities or systemic compound interactions across multiple molecular targets that orchestrate behavior. Our findings indicate that compounds not traditionally classified as NMDAR modulators can still impact habituation, often producing behavioral phenotypes distinct from those of chemical compounds primarily known to modulate NMDARs with higher specificity (e.g., APV, MK-801). This highlights the VAMR assay’s ability to capture diverse mechanistic interactions beyond direct receptor antagonism, making it a valuable tool for investigating complex neuroactive compound effects. The mode(s) of action underlying behavioral alterations in zebrafish exposed to MK-801 for example, are not completely understood. *grin1* encodes the obligatory GluN1 subunit of NMDARs^100^. While behavioral defects in *grin1* double mutants are generally phenocopied by short-term MK-801 treatment^101^, off-target activities on acetylcholine receptors^102^ and voltage-gated potassium channels^103^ may explain the difference in phenotypes observed in zebrafish exposed to MK-801 and APV. Overall, our data indicate that zebrafish multi-behavioral phenotyping strategies can be used to identify complex, system-modulating compounds.

We found that exposure to chlorophene, a chlorinated phenolic compound, caused reduced habituation learning, sedation, and paradoxical excitation in zebrafish. Sedatives and general anesthetics (e.g. benzodiazepines, propofol, etomidate) can cause paradoxical excitation where an opposite, paradoxical reaction induces both sedation and increased brain and/or motor activity^104–106^. It is widely accepted that anesthetics induce neuronal suppression via GABAR agonism, NMDAR antagonism, or by membrane polarity modulation^107–111^. Several lines of evidence indicate that chlorophene also acts on GABA_A_Rs. First, among other proteins, SEA algorithms^68^ predicted multiple GABA_A_R subunits to be targeted chlorophene, including subunits α2, α3, and α5. Second, chemical intervention using the GABA_A_R antagonist picrotoxin counteracted sedation of visual motor behaviors in zebrafish. Third, tonic inhibition against neuronal excitability in cultured mouse cortical neurons was blocked by the GABA_A_R antagonists picrotoxin and SR95531. Fourth, chlorophene exposure reduced the normalized spiking count of GABAergic units in hiPSC-derived 3D BrainSpheres.

Most GABA_A_R α subunits are expressed during zebrafish embryogenesis^112^. The amino acid sequences and electrophysiological properties of GABA_A_R subunits are largely conserved among vertebrates including zebrafish, mice, and humans^113^. Several studies suggest that α5-GABA_A_R-mediated tonic inhibition reduces learning and memory^114–120^. Under physiological conditions, tonic GABA_A_R currents cause hyperpolarization and decrease neuronal excitability. Synaptic input, therefore, becomes subthreshold and interferes with coordinated pre- and post-synaptic spiking activity, which is required for NMDA-dependent long-term plasticity^121^. Antagonizing GABAergic tone facilitates neuroplasticity in genetic mouse models for cognitive disabilities^122,123^. In contrast, the deficits in learning and memory in zebrafish described here were not rescued by the GABA_A_R antagonist picrotoxin. Our results indicate that reduced habituation may result from a mechanism distinct from functional GABA_A_R agonism. Alternatively, co-occurring phenotypes including sedation, paradoxical excitation, and defective learning, may causally relate to a common, previously unknown upstream regulator (see discussion below).

NMDARs are essential to mediate experience-dependent neuroplasticity underlying learning and memory^75,100^. In teleost fish, including zebrafish, the telencephalon, and in particular the medial and lateral pallium, are analogous to amygdala and hippocampus in mammals^124–126^ and represent the neuroanatomical structures responsible for learning and memory involving NMDAR activation^127–129^. However, our electrophysiological data indicates that chlorophene does not directly interfere with NMDAR-dependent signaling in mouse cortical neurons. This suggests that chlorophene indirectly affects NMDARs, phenocopying the lower habituation observed in zebrafish exposed to NAMDAR antagonists. A large-scale screen for chemical modulators of non-associative learning identified additional mechanisms reducing short-term habituation in larval zebrafish including potassium channel antagonism^44^. Interestingly, another zebrafish-based small molecule screen implicated that the M-current, a voltage-gated potassium channel that regulates neuronal excitability^130^, is related to sedation and paradoxical excitation^49^. Chlorophene’s behavioral profile, characterized by sedation, paradoxical excitation, and habituation deficits, closely resembled that of flupirtine, a non-opioid analgesic known to activate Kv7 potassium channels (M-current) and potentiate GABAergic transmission^82,131,132^. Given that Kv7 channels integrate excitatory and inhibitory signaling^82,131,132^, our findings reinforce the role of M-current modulation as a potential mechanism underlying chlorophene’s neurobehavioral effects. This interpretation is further supported by recent findings demonstrating that voltage-gated potassium channels responsible for generating the M-current can be directly activated by endogenous neurotransmitters, including GABA and related metabolites^133^. These findings suggest that chlorophene’s impact on habituation may not result solely from GABA_A_R modulation but rather from a combination of GABAergic, glutamatergic, and voltage-gated potassium channel interactions. By capturing polypharmacological effects involving GABA_A_R modulation and Kv7 channel activation, our study demonstrates that the VAMR assay extends beyond direct NMDAR antagonism to reveal a broader range of neuroactive mechanisms. Our screen confirmed that exposure to diphenhydramine, an antihistamine used in some hypnotic preparations, caused sedation and paradoxical excitation in early-life-stage zebrafish^134^. Surprisingly, as compared to chlorophene, diphenhydramine exposure had the opposite effect on non-associative learning, enhancing short-term habituation in zebrafish. Future studies should investigate whether M-current modulators counteract the sedative, paradoxical excitatory, and habituative learning effects of compounds such as chlorophene and diphenhydramine.

Chlorophene and its structural analogues, known as chlorophenes, represent a group of halogenated phenolic compounds that are globally used as antimicrobial agents and preservatives in personal care products and cosmetics^80,135^. Their environmental occurrence in surface waters and soils exclusively results from anthropogenic activities through emissions from sewage treatment plants and application of sewage sludge^135–137^. Based on a predicted environmental concentration (PEC) of 0.2 µg/L (0.9 nM) and a predicted no effect concentration (PNEC) of 0.058 µg/L (0.3 nM), a risk quotient (RQ=PEC/PNEC) of 3.4 has been calculated for the emission of chlorophene into surface waters, indicating that chlorophene may pose an unreasonable risk to the aquatic environment^135,138^. Consequently, chlorophene is included on international lists of environmental chemicals of emerging concern^135^. This, paired with the toxicity data reported here, suggests that (bio)monitoring in human and environmental matrices is justified.

Chlorophenes are hydrophobic (log K_OW_ 4.3-7.5), persistent (ultimate biodegradation half-life 1.9-4.9 years), low-to-moderately bioaccumulative in fish (BCF 55-400 L/kg), and toxic to aquatic organisms including fish, invertebrates, and algae^135,138^. In particular, chlorophene’s 30-day no observed effect concentration (NOEC_chronic_) for survival in zebrafish (Fish Early Life Stage Test, OECD TG 210^139^) of 0.58 mg/L (2.7 nM) indicates high chronic toxicity (acute-to-chronic ratio; ACR=LC50_acute_/NOEC_chronic_=569) and is consistent with a specific mode of action^79,135,138,140^. Our study suggests that chlorophene acts as both a functional agonist at GABA_A_Rs and an M-current activator, indicating a neurotoxic mode of action. Currently, adult or developmental neurotoxicity assessment is addressed under the Classification, Labeling and Packaging (CLP) Regulation^141,142^, REACH (Registration, Evaluation, Authorisation and Restriction of Chemicals), and the Biocidal Products Regulation (BPR). Under CLP, DNT studies are required if there is evidence that the nervous system is a target organ including any effect interfering with normal development until sexual maturation, including death, structural abnormality, altered growth, and functional deficits^143^, whereas both under REACH and BPR, adult neurotoxicity (functional or morphological alterations in the mature nervous system) triggers tests on developmental neurotoxicity^144,145^. Interestingly, an ECHA (European Chemicals Agency) assessment reports state that chlorophene has “…no structural similarity to […] known inducers of delayed neurotoxicity” and that “[…] studies in several species did not reveal the potential for neurotoxic effects […]”^79^. This discrepancy underscores limitations in current neurotoxicity assessment practices as this conclusion was reached without conducting regulatory adult or developmental neurotoxicity tests.

In addition to neurotoxicity, there is increasing evidence that chlorophene and related chemical compounds act as endocrine-disrupting chemicals (EDCs) that interfere with hormonal homeostasis. In cell-based luciferase reporter assays *in vitro*, chlorophene modulated androgen, estrogen, and thyroid receptor activity, and meets the criteria for an EDC^146^. A structurally related compound, bromochlorophene, was shown to act as an antagonist on androgen, estrogen, glucocorticoid, and thyroid receptors *in vitro*^146^. Given that GABA is widely distributed in endocrine tissues and serves as a hormone or trophic factor in neuroendocrine systems^147,148^, and considering that changes in hypothalamic GABA levels are involved in endocrine disorders^149,150^, a potential link between altered GABAergic signaling and endocrine disruption should be addressed in EDC-related studies.

In the future, zebrafish multi-behavioral assays might be utilized for the regulatory assessment of chemical-induced neurotoxicity. The European Union’s chemicals strategy (European Green Deal) aims to enhance innovation for safe and sustainable chemicals to better protect human health and the environment^151^. To improve public health and environmental protection under legislative instruments such as REACH and CLP, current risk assessment should prevent the use of chemicals that affect neurological systems^151^. Because current regulatory DNT testing is exclusively conducted in rodents (e.g., OECD Test No. 426: Developmental Neurotoxicity Study^24^), less than 0.06% of an estimated 350,000 commercially available chemicals^22^ have been evaluated for potential effects on the developing nervous system^21^. As a result, there is an urgent need for alternative, rapid, inexpensive, and 3R-compliant^36^ approaches to evaluate potential DNT effects. Compared to cell-based *in vitro* neurotoxicity methods^152,153^, multi-behavioral phenotyping in zebrafish represents a more holistic and untargeted approach to identify neurotoxicants and to infer their potential chemical mode(s) of action. In addition, unlike *in vitro*-based approaches, early-life-stage zebrafish capture the functional complexity of the vertebrate nervous system^154^, including its conserved developmental biology that is relevant to higher vertebrates including humans^155^. In addition, zebrafish are metabolically competent, enabling the identification and characterization of adverse effects derived from biotransformation products^156,157^. While the zebrafish phenotyping battery contains multiple learning- and memory-related readouts including habituation, potentiation of habituation^44^, and memory retention^44,54^ that serve as potential correlates to behavior-based endpoints in OECD rodent neurotoxicity test guidelines^24^, more work is needed to understand the direct relationship between rodent- and zebrafish-based behavior readouts.

In summary, we combined a series of behavior assays in a non-mammalian vertebrate model to screen chemicals for neurotoxicity in a phenotypically rich, rapid, and inexpensive test system. Together with *in silico* target predictions, and mouse- and human-based models, our findings establish multi-behavioral phenotyping as a powerful toolkit for neurotoxicity testing and mechanism identification, with cross-species relevance. More broadly, multi-behavioral phenotyping in zebrafish can accelerate the identification of neuroactive chemicals that should be prioritized for additional testing to better protect human health and the environment from chemicals that disrupt nervous system function.

## Acknowledgements

We thank Gi Mick Wu (UFZ) for statistical consultation. We also thank Janet Krüger (UFZ) for technical assistance in the lab. This work was funded by a W2 Helmholtz Association Grant (T.T.). This work was also carried out in the framework of the European Partnership for the Assessment of Risks from Chemicals (PARC) and has received funding from the European Union’s Horizon Europe research and innovation programme under Grant Agreement No 101057014 (T.T. and E.F.).

## Author contributions

D.L. designed and performed the research, analyzed the data, and wrote the manuscript. N.K.H., J.N., K.B., and I.S. performed experiments and analyzed data. T.T. designed the study and wrote the manuscript. All authors contributed to data interpretation and commented on the manuscript.

