## Supplemental material for "Multi-behavioral phenotyping in early-life-stage zebrafish for identifying disruptors of non-associative learning"

### 1 Supplemental Tables

**Table S1.** **Structures, identifiers, supplier, purity, molecular weight and LogD of chemical compounds investigated in this study.** Octanol-water partitioning coefficients (LogD) were calculated using the LogD module of ACD/Percepta (Advanced Chemistry Development Inc., ACD/Labs 2015 release, build 2726, 27 Nov 2014). MW, molecular weight.

| **Structure** | **Chemical** | **CAS number** | **DTXSID** | **SMILES** | **Supplier** | **Purity (%)** | **MW (g/mol)** | **LogD (pH=7.4)** |
| --- | --- | --- | --- | --- | --- | --- | --- | --- |
| 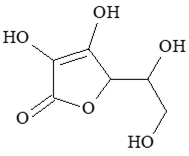 | (+)-Sodium L-ascorbate | 134-03-2 | DTXSID0020105 | C(C(C1C(=C(C(=O)O1)O)[O-])O)O.[Na+] | Sigma-Aldrich | ≥98 | 198.11 | 3.42 |
| 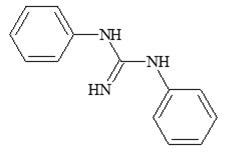 | 1,3-Diphenylguanidine | 102-06-7 | DTXSID3025178 | N=C(NC1=CC=CC=C1)NC1=CC=CC=C1 | Alfa Aesar | 99.5 | 211.26 | 1.27 |
| 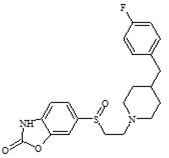 | Besonprodil | 253450-09-8 | DTXSID2047270 | FC1=CC=C(CC2CCN(CCS(=O)C3=CC=C4NC(=O)OC4=C3)CC2)C=C1 | Sigma-Aldrich | NA | 402.49 | 6.05 |
| 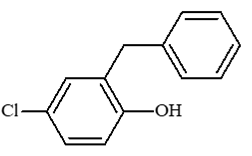 | Chlorophene | 120-32-1 | DTXSID5020154 | OC1=C(CC2=CC=CC=C2)C=C(Cl)C=C1 | Sigma-Aldrich | 95 | 218.68 | 2.34 |
| 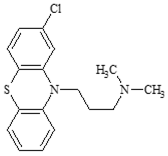 | Chlorpromazine hydrochloride | 69-09-0 | DTXSID7024827 | Cl.CN(C)CCCN1C2=C(SC3=C1C=C(Cl)C=C3)C=CC=C2 | Sigma-Aldrich | ≥98 | 355.33 | 2.68 |
| 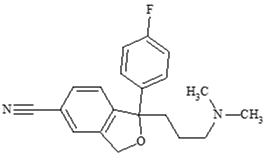 | Citalopram hydrobromide | 59729-32-7 | DTXSID40872344 | CN(C)CCCC1(C2=C(CO1)C=C(C=C2)C#N)C3=CC=C(C=C3)F.Br | Sigma-Aldrich | ≥98 | 405.30 | -5.97 |
| 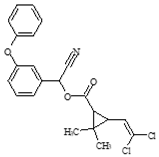 | Cypermethrin | 52315-07-8 | DTXSID1023998 | CC1(C(C1C(=O)OC(C#N)C2=CC(=CC=C2)OC3=CC=CC=C3)C=C(Cl)Cl)C | Supleco | ≥90 | 416.30 | 2.79 |
| 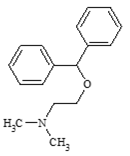 | Diphenhydramine hydrochloride | 147-24-0 | DTXSID4020537 | Cl.CN(C)CCOC(C1=CC=CC=C1)C1=CC=CC=C1 | Sigma-Aldrich | ≥98 | 291.81 | 2.65 |
| 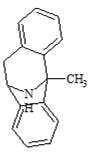 | Dizocilpin maleate (MK-801) | 77086-22-7 | DTXSID2045785 | CC12C3=CC=CC=C3CC(N1)C4=CC=CC=C24 | Alfa Aeser | ≥99 | 337.375 | -3.45 |
| 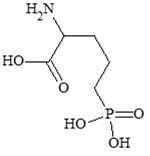 | DL-2-Amino-5-phosphonopentanoic acid (APV) | 76326-31-3 | DTXSID90893701 | C(CC(C(=O)O)N)CP(=O)(O)O | Sigma-Aldrich | ≥98 | 197.13 | -3.45 |
| 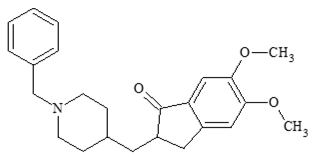 | Donepezil hydrochloride | 120011-70-3 | DTXSID0046698 | COC1=C(C=C2C(=C1)CC(C2=O)CC3CCN(CC3)CC4=CC=CC=C4)OC.Cl | Sigma-Aldrich | ≥98 | 415.95 | 2.8 |
| 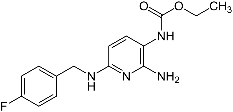 | Flupirtine maleate salt | 75507-68-5 | DTXSID8045771 | CCOC(=O)NC1=C(N=C(C=C1)NCC2=CC=C(C=C2)F)N | Sigma-Aldrich | ≥98 | 420.39 | 2.4 |
| 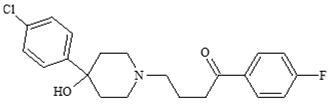 | Haloperidol | 52-86-8 | DTXSID4034150 | C1CN(CCC1(C2=CC=C(C=C2)Cl)O)CCCC(=O)C3=CC=C(C=C3)F | Sigma-Aldrich | ≥98 | 375.86 | -0.91 |
| 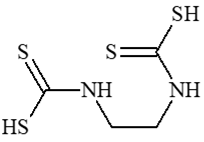 | Mancozeb | 8018-01-7 | DTXSID0034695 | C(CNC(=S)[S-])NC(=S)[S-].C(CNC(=S)[S-])NC(=S)[S-].[Mn+2].[Zn+2] | HPC Standards | PESTANAL®, analytical standard | 541.07 | -0.02 |
| 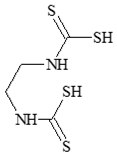 | Maneb | 12427-38-2 | DTXSID9020794 | [Mn++].[S-]C(=S)NCCNC([S-])=S | HPC Standards | PESTANAL®, analytical standard | 265.3 (as monomer) | 2.78 |
| 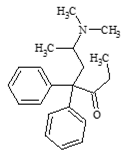 | Methadone hydrochloride | 1095-90-5 | DTXSID2020501 | Cl.CCC(=O)C(CC(C)N(C)C)(C1=CC=CC=C1)C1=CC=CC=C1 | Sigma-Aldrich | ≥98 | 345.91 | 1.58 |
| 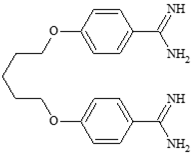 | Pentamidine isethionate | 140-64-7 | DTXSID5023796 | OCCS(O)(=O)=O.OCCS(O)(=O)=O.NC(=N)C1=CC=C(OCCCCCOC2=CC=C(C=C2)C(N)=N)C=C1 | Sigma-Aldrich | ≥98 | 592.68 | 3.42 |
| 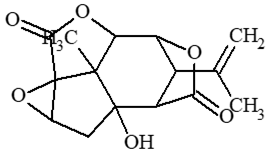 | Picrotoxin | 124-87-8 | DTXSID7045605 | CC(=C)C1C2C3C4(C(C1C(=O)O2)(CC5C4(O5)C(=O)O3)O)C.CC12C3C4C(C(C1(CC5C2(O5)C(=O)O3)O)C(=O)O4)C(C)(C)O | Sigma-Aldrich | ≥98 | 602.58 | 1.27 |
| 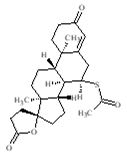 | Spironolactone | 52-01-7 | DTXSID6034186 | [H][C@@]12CC[C@@]3(CCC(=O)O3)[C@@]1(C)CC[C@@]1([H])[C@@]2([H])[C@@H](CC2=CC(=O)CC[C@]12C)SC(C)=O | Alfa Aesar | 97-103 | 416.57 | 6.05 |
| 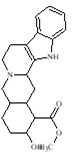 | Yohimbine hydrochloride | 65-19-0 | DTXSID40891272 | COC(=O)C1C(CCC2C1CC3C4=C(CCN3C2)C5=CC=CC=C5N4)O.Cl | Sigma-Aldrich | ≥98 | 390.90 | 2.34 |

**Table S2. Reference set of habituation modulators in larval zebrafish.**

| **Compound** | **Pathway** | **Target** | **Function** | **Habituation** | **Reference** |
| --- | --- | --- | --- | --- | --- |
| (2*R*)-amino-5-phosphonovaleric acid (APV) | Glutamatergic | NMDA receptor | Antagonist, competitive | Reduced | Roberts et al. 2011^2^ |
| Citalopram | Serotinergic | Selective serotonin reuptake | Inhibitor | Enhanced | Wolman et al. 2011^3^ |
| Donepezil | Cholinergic | Acetylcholinesterase | Inhibitor, non-esterifying | Reduced | Best et al. 2008^4^ |
| Haloperidol | Dopaminergic | Dopamine receptor | Antagonist | Enhanced | Wolman et al. 2011^3^ |
| MK-801 | Glutamatergic | NMDA receptor | Antagonist, non-competitive | Reduced | Wolman et al. 2011^3^ |
| Picrotoxin | GABAergic | GABAA receptor | Antagonist, ion channel blockage | Reduced | Wolman et al. 2011^3^ |
| Yohimbine | Adrenergic | α2-adrenoreceptor | Antagonist | Enhanced | Wolman et al. 2011^3^ |

**Table S3.** **Assay battery stimulus configuration.** BSL, baseline; VSR, visual startle response; VMR, visual motor response; ASR, acoustic startle response; ASH, acoustic startle habituation.

| **Assay order** | **Assay name** | **Stimulus name** | **Stimulus no.** | **Start (s)** | **End (s)** |
| --- | --- | --- | --- | --- | --- |
| NA | NA | Visual light (100%) | NA | 0 | 60 |
| NA | NA | IR light | NA | 60 | 1200 |
| 1 | BSL | IR light | NA | 1200 | 1260 |
| 2 | VSR1 | Visual light (100%) | NA | 1260 | 1261 |
| 3 | VMR1 | Visual light (100%) | NA | 1261 | 1860 |
| 4 | VSR2 | IR light | NA | 1860 | 1861 |
| 5 | VMR2 | IR light | NA | 1861 | 2101 |
| 6 | VMR3 | IR light | NA | 2101 | 3060 |
| 7 | ASR1 | IR light, low acoustic (75%) | 1 | 3060 | 3061 |
| 7 | ASR1 | IR light | NA | 3061 | 3121 |
| 7 | ASR1 | IR light, low acoustic (75%) | 2 | 3121 | 3122 |
| 7 | ASR1 | IR light | NA | 3122 | 3182 |
| 7 | ASR1 | IR light, low acoustic (75%) | 3 | 3182 | 3183 |
| 7 | ASR1 | IR light | NA | 3183 | 3243 |
| 7 | ASR1 | IR light, low acoustic (75%) | 4 | 3243 | 3244 |
| 7 | ASR1 | IR light | NA | 3244 | 3304 |
| 7 | ASR1 | IR light, low acoustic (75%) | 5 | 3304 | 3305 |
| NA | NA | IR light | NA | 3305 | 3365 |
| 8 | ASR2 | IR light, high acoustic (100%) | 1 | 3365 | 3366 |
| 8 | ASR2 | IR light | NA | 3366 | 3426 |
| 8 | ASR2 | IR light, high acoustic (100%) | 2 | 3426 | 3427 |
| 8 | ASR2 | IR light | NA | 3427 | 3487 |
| 8 | ASR2 | IR light, high acoustic (100%) | 3 | 3487 | 3488 |
| 8 | ASR2 | IR light | NA | 3488 | 3548 |
| 8 | ASR2 | IR light, high acoustic (100%) | 4 | 3548 | 3549 |
| 8 | ASR2 | IR light | NA | 3549 | 3609 |
| 8 | ASR2 | IR light, high acoustic (100%) | 5 | 3609 | 3610 |
| NA | NA | IR light | NA | 3610 | 3670 |
| 9 | ASH1 | IR light, high acoustic (100%) | 1 | 3670 | 3671 |
| 9 | ASH1 | IR light | NA | 3671 | 3672 |
| 9 | ASH1 | IR light, high acoustic (100%) | 2 | 3672 | 3673 |
| 9 | ASH1 | IR light | NA | 3673 | 3674 |
| 9 | ASH1 | IR light, high acoustic (100%) | 3 | 3674 | 3675 |
| 9 | ASH1 | IR light | NA | 3675 | 3676 |
| 9 | ASH1 | IR light, high acoustic (100%) | 4 | 3676 | 3677 |
| 9 | ASH1 | IR light | NA | 3677 | 3678 |
| 9 | ASH1 | IR light, high acoustic (100%) | 5 | 3678 | 3679 |
| 9 | ASH1 | IR light | NA | 3679 | 3680 |
| 9 | ASH1 | IR light, high acoustic (100%) | 6 | 3680 | 3681 |
| 9 | ASH1 | IR light | NA | 3681 | 3682 |
| 9 | ASH1 | IR light, high acoustic (100%) | 7 | 3682 | 3683 |
| 9 | ASH1 | IR light | NA | 3683 | 3684 |
| 9 | ASH1 | IR light, high acoustic (100%) | 8 | 3684 | 3685 |
| 9 | ASH1 | IR light | NA | 3685 | 3686 |
| 9 | ASH1 | IR light, high acoustic (100%) | 9 | 3686 | 3687 |
| 9 | ASH1 | IR light | NA | 3687 | 3688 |
| 9 | ASH1 | IR light, high acoustic (100%) | 10 | 3688 | 3689 |
| 9 | ASH1 | IR light | NA | 3689 | 3690 |
| 9 | ASH1 | IR light, high acoustic (100%) | 11 | 3690 | 3691 |
| 9 | ASH1 | IR light | NA | 3691 | 3692 |
| 9 | ASH1 | IR light, high acoustic (100%) | 12 | 3692 | 3693 |
| 9 | ASH1 | IR light | NA | 3693 | 3694 |
| 9 | ASH1 | IR light, high acoustic (100%) | 13 | 3694 | 3695 |
| 9 | ASH1 | IR light | NA | 3695 | 3696 |
| 9 | ASH1 | IR light, high acoustic (100%) | 14 | 3696 | 3697 |
| 9 | ASH1 | IR light | NA | 3697 | 3698 |
| 9 | ASH1 | IR light, high acoustic (100%) | 15 | 3698 | 3699 |
| 9 | ASH1 | IR light | NA | 3699 | 3700 |
| 9 | ASH1 | IR light, high acoustic (100%) | 16 | 3700 | 3701 |
| 9 | ASH1 | IR light | NA | 3701 | 3702 |
| 9 | ASH1 | IR light, high acoustic (100%) | 17 | 3702 | 3703 |
| 9 | ASH1 | IR light | NA | 3703 | 3704 |
| 9 | ASH1 | IR light, high acoustic (100%) | 18 | 3704 | 3705 |
| 9 | ASH1 | IR light | NA | 3705 | 3706 |
| 9 | ASH1 | IR light, high acoustic (100%) | 19 | 3706 | 3707 |
| 9 | ASH1 | IR light | NA | 3707 | 3708 |
| 9 | ASH1 | IR light, high acoustic (100%) | 20 | 3708 | 3709 |
| 9 | ASH1 | IR light | NA | 3709 | 3710 |
| 9 | ASH1 | IR light, high acoustic (100%) | 21 | 3710 | 3711 |
| 9 | ASH1 | IR light | NA | 3711 | 3712 |
| 9 | ASH1 | IR light, high acoustic (100%) | 22 | 3712 | 3713 |
| 9 | ASH1 | IR light | NA | 3713 | 3714 |
| 9 | ASH1 | IR light, high acoustic (100%) | 23 | 3714 | 3715 |
| 9 | ASH1 | IR light | NA | 3715 | 3716 |
| 9 | ASH1 | IR light, high acoustic (100%) | 24 | 3716 | 3717 |
| 9 | ASH1 | IR light | NA | 3717 | 3718 |
| 9 | ASH1 | IR light, high acoustic (100%) | 25 | 3718 | 3719 |
| 9 | ASH1 | IR light | NA | 3719 | 3720 |
| 9 | ASH1 | IR light, high acoustic (100%) | 26 | 3720 | 3721 |
| 9 | ASH1 | IR light | NA | 3721 | 3722 |
| 9 | ASH1 | IR light, high acoustic (100%) | 27 | 3722 | 3723 |
| 9 | ASH1 | IR light | NA | 3723 | 3724 |
| 9 | ASH1 | IR light, high acoustic (100%) | 28 | 3724 | 3725 |
| 9 | ASH1 | IR light | NA | 3725 | 3726 |
| 9 | ASH1 | IR light, high acoustic (100%) | 29 | 3726 | 3727 |
| 9 | ASH1 | IR light | NA | 3727 | 3728 |
| 9 | ASH1 | IR light, high acoustic (100%) | 30 | 3728 | 3729 |
| NA | NA | IR light | NA | 3729 | 3789 |
| 10 | ASH2 | IR light, high acoustic (100%) | 1 | 3789 | 3790 |
| 10 | ASH2 | IR light | NA | 3790 | 3791 |
| 10 | ASH2 | IR light, high acoustic (100%) | 2 | 3791 | 3792 |
| 10 | ASH2 | IR light | NA | 3792 | 3793 |
| 10 | ASH2 | IR light, high acoustic (100%) | 3 | 3793 | 3794 |
| 10 | ASH2 | IR light | NA | 3794 | 3795 |
| 10 | ASH2 | IR light, high acoustic (100%) | 4 | 3795 | 3796 |
| 10 | ASH2 | IR light | NA | 3796 | 3797 |
| 10 | ASH2 | IR light, high acoustic (100%) | 5 | 3797 | 3798 |
| 10 | ASH2 | IR light | NA | 3798 | 3799 |
| 10 | ASH2 | IR light, high acoustic (100%) | 6 | 3799 | 3800 |
| 10 | ASH2 | IR light | NA | 3800 | 3801 |
| 10 | ASH2 | IR light, high acoustic (100%) | 7 | 3801 | 3802 |
| 10 | ASH2 | IR light | NA | 3802 | 3803 |
| 10 | ASH2 | IR light, high acoustic (100%) | 8 | 3803 | 3804 |
| 10 | ASH2 | IR light | NA | 3804 | 3805 |
| 10 | ASH2 | IR light, high acoustic (100%) | 9 | 3805 | 3806 |
| 10 | ASH2 | IR light | NA | 3806 | 3807 |
| 10 | ASH2 | IR light, high acoustic (100%) | 10 | 3807 | 3808 |
| 10 | ASH2 | IR light | NA | 3808 | 3809 |
| 10 | ASH2 | IR light, high acoustic (100%) | 11 | 3809 | 3810 |
| 10 | ASH2 | IR light | NA | 3810 | 3811 |
| 10 | ASH2 | IR light, high acoustic (100%) | 12 | 3811 | 3812 |
| 10 | ASH2 | IR light | NA | 3812 | 3813 |
| 10 | ASH2 | IR light, high acoustic (100%) | 13 | 3813 | 3814 |
| 10 | ASH2 | IR light | NA | 3814 | 3815 |
| 10 | ASH2 | IR light, high acoustic (100%) | 14 | 3815 | 3816 |
| 10 | ASH2 | IR light | NA | 3816 | 3817 |
| 10 | ASH2 | IR light, high acoustic (100%) | 15 | 3817 | 3818 |
| 10 | ASH2 | IR light | NA | 3818 | 3819 |
| 10 | ASH2 | IR light, high acoustic (100%) | 16 | 3819 | 3820 |
| 10 | ASH2 | IR light | NA | 3820 | 3821 |
| 10 | ASH2 | IR light, high acoustic (100%) | 17 | 3821 | 3822 |
| 10 | ASH2 | IR light | NA | 3822 | 3823 |
| 10 | ASH2 | IR light, high acoustic (100%) | 18 | 3823 | 3824 |
| 10 | ASH2 | IR light | NA | 3824 | 3825 |
| 10 | ASH2 | IR light, high acoustic (100%) | 19 | 3825 | 3826 |
| 10 | ASH2 | IR light | NA | 3826 | 3827 |
| 10 | ASH2 | IR light, high acoustic (100%) | 20 | 3827 | 3828 |
| 10 | ASH2 | IR light | NA | 3828 | 3829 |
| 10 | ASH2 | IR light, high acoustic (100%) | 21 | 3829 | 3830 |
| 10 | ASH2 | IR light | NA | 3830 | 3831 |
| 10 | ASH2 | IR light, high acoustic (100%) | 22 | 3831 | 3832 |
| 10 | ASH2 | IR light | NA | 3832 | 3833 |
| 10 | ASH2 | IR light, high acoustic (100%) | 23 | 3833 | 3834 |
| 10 | ASH2 | IR light | NA | 3834 | 3835 |
| 10 | ASH2 | IR light, high acoustic (100%) | 24 | 3835 | 3836 |
| 10 | ASH2 | IR light | NA | 3836 | 3837 |
| 10 | ASH2 | IR light, high acoustic (100%) | 25 | 3837 | 3838 |
| 10 | ASH2 | IR light | NA | 3838 | 3839 |
| 10 | ASH2 | IR light, high acoustic (100%) | 26 | 3839 | 3840 |
| 10 | ASH2 | IR light | NA | 3840 | 3841 |
| 10 | ASH2 | IR light, high acoustic (100%) | 27 | 3841 | 3842 |
| 10 | ASH2 | IR light | NA | 3842 | 3843 |
| 10 | ASH2 | IR light, high acoustic (100%) | 28 | 3843 | 3844 |
| 10 | ASH2 | IR light | NA | 3844 | 3845 |
| 10 | ASH2 | IR light, high acoustic (100%) | 29 | 3845 | 3846 |
| 10 | ASH2 | IR light | NA | 3846 | 3847 |
| 10 | ASH2 | IR light, high acoustic (100%) | 30 | 3847 | 3848 |
| NA | NA | IR light | NA | 3848 | 3908 |
| 11 | ASH3 | IR light, high acoustic (100%) | 1 | 3908 | 3909 |
| 11 | ASH3 | IR light | NA | 3909 | 3910 |
| 11 | ASH3 | IR light, high acoustic (100%) | 2 | 3910 | 3911 |
| 11 | ASH3 | IR light | NA | 3911 | 3912 |
| 11 | ASH3 | IR light, high acoustic (100%) | 3 | 3912 | 3913 |
| 11 | ASH3 | IR light | NA | 3913 | 3914 |
| 11 | ASH3 | IR light, high acoustic (100%) | 4 | 3914 | 3915 |
| 11 | ASH3 | IR light | NA | 3915 | 3916 |
| 11 | ASH3 | IR light, high acoustic (100%) | 5 | 3916 | 3917 |
| 11 | ASH3 | IR light | NA | 3917 | 3918 |
| 11 | ASH3 | IR light, high acoustic (100%) | 6 | 3918 | 3919 |
| 11 | ASH3 | IR light | NA | 3919 | 3920 |
| 11 | ASH3 | IR light, high acoustic (100%) | 7 | 3920 | 3921 |
| 11 | ASH3 | IR light | NA | 3921 | 3922 |
| 11 | ASH3 | IR light, high acoustic (100%) | 8 | 3922 | 3923 |
| 11 | ASH3 | IR light | NA | 3923 | 3924 |
| 11 | ASH3 | IR light, high acoustic (100%) | 9 | 3924 | 3925 |
| 11 | ASH3 | IR light | NA | 3925 | 3926 |
| 11 | ASH3 | IR light, high acoustic (100%) | 10 | 3926 | 3927 |
| 11 | ASH3 | IR light | NA | 3927 | 3928 |
| 11 | ASH3 | IR light, high acoustic (100%) | 11 | 3928 | 3929 |
| 11 | ASH3 | IR light | NA | 3929 | 3930 |
| 11 | ASH3 | IR light, high acoustic (100%) | 12 | 3930 | 3931 |
| 11 | ASH3 | IR light | NA | 3931 | 3932 |
| 11 | ASH3 | IR light, high acoustic (100%) | 13 | 3932 | 3933 |
| 11 | ASH3 | IR light | NA | 3933 | 3934 |
| 11 | ASH3 | IR light, high acoustic (100%) | 14 | 3934 | 3935 |
| 11 | ASH3 | IR light | NA | 3935 | 3936 |
| 11 | ASH3 | IR light, high acoustic (100%) | 15 | 3936 | 3937 |
| 11 | ASH3 | IR light | NA | 3937 | 3938 |
| 11 | ASH3 | IR light, high acoustic (100%) | 16 | 3938 | 3939 |
| 11 | ASH3 | IR light | NA | 3939 | 3940 |
| 11 | ASH3 | IR light, high acoustic (100%) | 17 | 3940 | 3941 |
| 11 | ASH3 | IR light | NA | 3941 | 3942 |
| 11 | ASH3 | IR light, high acoustic (100%) | 18 | 3942 | 3943 |
| 11 | ASH3 | IR light | NA | 3943 | 3944 |
| 11 | ASH3 | IR light, high acoustic (100%) | 19 | 3944 | 3945 |
| 11 | ASH3 | IR light | NA | 3945 | 3946 |
| 11 | ASH3 | IR light, high acoustic (100%) | 20 | 3946 | 3947 |
| 11 | ASH3 | IR light | NA | 3947 | 3948 |
| 11 | ASH3 | IR light, high acoustic (100%) | 21 | 3948 | 3949 |
| 11 | ASH3 | IR light | NA | 3949 | 3950 |
| 11 | ASH3 | IR light, high acoustic (100%) | 22 | 3950 | 3951 |
| 11 | ASH3 | IR light | NA | 3951 | 3952 |
| 11 | ASH3 | IR light, high acoustic (100%) | 23 | 3952 | 3953 |
| 11 | ASH3 | IR light | NA | 3953 | 3954 |
| 11 | ASH3 | IR light, high acoustic (100%) | 24 | 3954 | 3955 |
| 11 | ASH3 | IR light | NA | 3955 | 3956 |
| 11 | ASH3 | IR light, high acoustic (100%) | 25 | 3956 | 3957 |
| 11 | ASH3 | IR light | NA | 3957 | 3958 |
| 11 | ASH3 | IR light, high acoustic (100%) | 26 | 3958 | 3959 |
| 11 | ASH3 | IR light | NA | 3959 | 3960 |
| 11 | ASH3 | IR light, high acoustic (100%) | 27 | 3960 | 3961 |
| 11 | ASH3 | IR light | NA | 3961 | 3962 |
| 11 | ASH3 | IR light, high acoustic (100%) | 28 | 3962 | 3963 |
| 11 | ASH3 | IR light | NA | 3963 | 3964 |
| 11 | ASH3 | IR light, high acoustic (100%) | 29 | 3964 | 3965 |
| 11 | ASH3 | IR light | NA | 3965 | 3966 |
| 11 | ASH3 | IR light, high acoustic (100%) | 30 | 3966 | 3967 |
| NA | NA | IR light | NA | 3967 | 4027 |
| 12 | ASH4 | IR light, high acoustic (100%) | 1 | 4027 | 4028 |
| 12 | ASH4 | IR light | NA | 4028 | 4029 |
| 12 | ASH4 | IR light, high acoustic (100%) | 2 | 4029 | 4030 |
| 12 | ASH4 | IR light | NA | 4030 | 4031 |
| 12 | ASH4 | IR light, high acoustic (100%) | 3 | 4031 | 4032 |
| 12 | ASH4 | IR light | NA | 4032 | 4033 |
| 12 | ASH4 | IR light, high acoustic (100%) | 4 | 4033 | 4034 |
| 12 | ASH4 | IR light | NA | 4034 | 4035 |
| 12 | ASH4 | IR light, high acoustic (100%) | 5 | 4035 | 4036 |
| 12 | ASH4 | IR light | NA | 4036 | 4037 |
| 12 | ASH4 | IR light, high acoustic (100%) | 6 | 4037 | 4038 |
| 12 | ASH4 | IR light | NA | 4038 | 4039 |
| 12 | ASH4 | IR light, high acoustic (100%) | 7 | 4039 | 4040 |
| 12 | ASH4 | IR light | NA | 4040 | 4041 |
| 12 | ASH4 | IR light, high acoustic (100%) | 8 | 4041 | 4042 |
| 12 | ASH4 | IR light | NA | 4042 | 4043 |
| 12 | ASH4 | IR light, high acoustic (100%) | 9 | 4043 | 4044 |
| 12 | ASH4 | IR light | NA | 4044 | 4045 |
| 12 | ASH4 | IR light, high acoustic (100%) | 10 | 4045 | 4046 |
| 12 | ASH4 | IR light | NA | 4046 | 4047 |
| 12 | ASH4 | IR light, high acoustic (100%) | 11 | 4047 | 4048 |
| 12 | ASH4 | IR light | NA | 4048 | 4049 |
| 12 | ASH4 | IR light, high acoustic (100%) | 12 | 4049 | 4050 |
| 12 | ASH4 | IR light | NA | 4050 | 4051 |
| 12 | ASH4 | IR light, high acoustic (100%) | 13 | 4051 | 4052 |
| 12 | ASH4 | IR light | NA | 4052 | 4053 |
| 12 | ASH4 | IR light, high acoustic (100%) | 14 | 4053 | 4054 |
| 12 | ASH4 | IR light | NA | 4054 | 4055 |
| 12 | ASH4 | IR light, high acoustic (100%) | 15 | 4055 | 4056 |
| 12 | ASH4 | IR light | NA | 4056 | 4057 |
| 12 | ASH4 | IR light, high acoustic (100%) | 16 | 4057 | 4058 |
| 12 | ASH4 | IR light | NA | 4058 | 4059 |
| 12 | ASH4 | IR light, high acoustic (100%) | 17 | 4059 | 4060 |
| 12 | ASH4 | IR light | NA | 4060 | 4061 |
| 12 | ASH4 | IR light, high acoustic (100%) | 18 | 4061 | 4062 |
| 12 | ASH4 | IR light | NA | 4062 | 4063 |
| 12 | ASH4 | IR light, high acoustic (100%) | 19 | 4063 | 4064 |
| 12 | ASH4 | IR light | NA | 4064 | 4065 |
| 12 | ASH4 | IR light, high acoustic (100%) | 20 | 4065 | 4066 |
| 12 | ASH4 | IR light | NA | 4066 | 4067 |
| 12 | ASH4 | IR light, high acoustic (100%) | 21 | 4067 | 4068 |
| 12 | ASH4 | IR light | NA | 4068 | 4069 |
| 12 | ASH4 | IR light, high acoustic (100%) | 22 | 4069 | 4070 |
| 12 | ASH4 | IR light | NA | 4070 | 4071 |
| 12 | ASH4 | IR light, high acoustic (100%) | 23 | 4071 | 4072 |
| 12 | ASH4 | IR light | NA | 4072 | 4073 |
| 12 | ASH4 | IR light, high acoustic (100%) | 24 | 4073 | 4074 |
| 12 | ASH4 | IR light | NA | 4074 | 4075 |
| 12 | ASH4 | IR light, high acoustic (100%) | 25 | 4075 | 4076 |
| 12 | ASH4 | IR light | NA | 4076 | 4077 |
| 12 | ASH4 | IR light, high acoustic (100%) | 26 | 4077 | 4078 |
| 12 | ASH4 | IR light | NA | 4078 | 4079 |
| 12 | ASH4 | IR light, high acoustic (100%) | 27 | 4079 | 4080 |
| 12 | ASH4 | IR light | NA | 4080 | 4081 |
| 12 | ASH4 | IR light, high acoustic (100%) | 28 | 4081 | 4082 |
| 12 | ASH4 | IR light | NA | 4082 | 4083 |
| 12 | ASH4 | IR light, high acoustic (100%) | 29 | 4083 | 4084 |
| 12 | ASH4 | IR light | NA | 4084 | 4085 |
| 12 | ASH4 | IR light, high acoustic (100%) | 30 | 4085 | 4086 |
| NA | NA | IR light | NA | 4086 | 4146 |
| 13 | ASH5 | IR light, high acoustic (100%) | 1 | 4146 | 4147 |
| 13 | ASH5 | IR light | NA | 4147 | 4148 |
| 13 | ASH5 | IR light, high acoustic (100%) | 2 | 4148 | 4149 |
| 13 | ASH5 | IR light | NA | 4149 | 4150 |
| 13 | ASH5 | IR light, high acoustic (100%) | 3 | 4150 | 4151 |
| 13 | ASH5 | IR light | NA | 4151 | 4152 |
| 13 | ASH5 | IR light, high acoustic (100%) | 4 | 4152 | 4153 |
| 13 | ASH5 | IR light | NA | 4153 | 4154 |
| 13 | ASH5 | IR light, high acoustic (100%) | 5 | 4154 | 4155 |
| 13 | ASH5 | IR light | NA | 4155 | 4156 |
| 13 | ASH5 | IR light, high acoustic (100%) | 6 | 4156 | 4157 |
| 13 | ASH5 | IR light | NA | 4157 | 4158 |
| 13 | ASH5 | IR light, high acoustic (100%) | 7 | 4158 | 4159 |
| 13 | ASH5 | IR light | NA | 4159 | 4160 |
| 13 | ASH5 | IR light, high acoustic (100%) | 8 | 4160 | 4161 |
| 13 | ASH5 | IR light | NA | 4161 | 4162 |
| 13 | ASH5 | IR light, high acoustic (100%) | 9 | 4162 | 4163 |
| 13 | ASH5 | IR light | NA | 4163 | 4164 |
| 13 | ASH5 | IR light, high acoustic (100%) | 10 | 4164 | 4165 |
| 13 | ASH5 | IR light | NA | 4165 | 4166 |
| 13 | ASH5 | IR light, high acoustic (100%) | 11 | 4166 | 4167 |
| 13 | ASH5 | IR light | NA | 4167 | 4168 |
| 13 | ASH5 | IR light, high acoustic (100%) | 12 | 4168 | 4169 |
| 13 | ASH5 | IR light | NA | 4169 | 4170 |
| 13 | ASH5 | IR light, high acoustic (100%) | 13 | 4170 | 4171 |
| 13 | ASH5 | IR light | NA | 4171 | 4172 |
| 13 | ASH5 | IR light, high acoustic (100%) | 14 | 4172 | 4173 |
| 13 | ASH5 | IR light | NA | 4173 | 4174 |
| 13 | ASH5 | IR light, high acoustic (100%) | 15 | 4174 | 4175 |
| 13 | ASH5 | IR light | NA | 4175 | 4176 |
| 13 | ASH5 | IR light, high acoustic (100%) | 16 | 4176 | 4177 |
| 13 | ASH5 | IR light | NA | 4177 | 4178 |
| 13 | ASH5 | IR light, high acoustic (100%) | 17 | 4178 | 4179 |
| 13 | ASH5 | IR light | NA | 4179 | 4180 |
| 13 | ASH5 | IR light, high acoustic (100%) | 18 | 4180 | 4181 |
| 13 | ASH5 | IR light | NA | 4181 | 4182 |
| 13 | ASH5 | IR light, high acoustic (100%) | 19 | 4182 | 4183 |
| 13 | ASH5 | IR light | NA | 4183 | 4184 |
| 13 | ASH5 | IR light, high acoustic (100%) | 20 | 4184 | 4185 |
| 13 | ASH5 | IR light | NA | 4185 | 4186 |
| 13 | ASH5 | IR light, high acoustic (100%) | 21 | 4186 | 4187 |
| 13 | ASH5 | IR light | NA | 4187 | 4188 |
| 13 | ASH5 | IR light, high acoustic (100%) | 22 | 4188 | 4189 |
| 13 | ASH5 | IR light | NA | 4189 | 4190 |
| 13 | ASH5 | IR light, high acoustic (100%) | 23 | 4190 | 4191 |
| 13 | ASH5 | IR light | NA | 4191 | 4192 |
| 13 | ASH5 | IR light, high acoustic (100%) | 24 | 4192 | 4193 |
| 13 | ASH5 | IR light | NA | 4193 | 4194 |
| 13 | ASH5 | IR light, high acoustic (100%) | 25 | 4194 | 4195 |
| 13 | ASH5 | IR light | NA | 4195 | 4196 |
| 13 | ASH5 | IR light, high acoustic (100%) | 26 | 4196 | 4197 |
| 13 | ASH5 | IR light | NA | 4197 | 4198 |
| 13 | ASH5 | IR light, high acoustic (100%) | 27 | 4198 | 4199 |
| 13 | ASH5 | IR light | NA | 4199 | 4200 |
| 13 | ASH5 | IR light, high acoustic (100%) | 28 | 4200 | 4201 |
| 13 | ASH5 | IR light | NA | 4201 | 4202 |
| 13 | ASH5 | IR light, high acoustic (100%) | 29 | 4202 | 4203 |
| 13 | ASH5 | IR light | NA | 4203 | 4204 |
| 13 | ASH5 | IR light, high acoustic (100%) | 30 | 4204 | 4205 |
| NA | NA | IR light | NA | 4205 | 4385 |
| 14 | ASR3 | IR light, high acoustic (100%) | 1 | 4385 | 4386 |
| 14 | ASR3 | IR light | NA | 4386 | 4446 |
| 14 | ASR3 | IR light, high acoustic (100%) | 2 | 4446 | 4447 |
| 14 | ASR3 | IR light | NA | 4447 | 4507 |
| 14 | ASR3 | IR light, high acoustic (100%) | 3 | 4507 | 4508 |
| 14 | ASR3 | IR light | NA | 4508 | 4568 |
| 14 | ASR3 | IR light, high acoustic (100%) | 4 | 4568 | 4569 |
| 14 | ASR3 | IR light | NA | 4569 | 4629 |
| 14 | ASR3 | IR light, high acoustic (100%) | 5 | 4629 | 4630 |
| NA | NA | IR light | NA | 4630 | 4633 |

**Table S4. Detailed assay information for the assay components NVS_LGIC_rGluNMDA_Agonist and NVS_LGIC_rGluNMDA_MK801_Agonist.** Data gathered from the EPA CompTox Chemicals Dashboard (https://comptox.epa.gov/dashboard/).

| **Property** | **NVS_LGIC_rGluNMDA_Agonist** | **NVS_LGIC_rGluNMDA_MK801_Agonist** |
| --- | --- | --- |
| Aeid | 702 | 703 |
| Assay Component Endpoint Name | NVS_LGIC_rGluNMDA_Agonist | NVS_LGIC_rGluNMDA_MK801_Agonist |
| Assay Component Endpoint Desc | Data from the assay component NVS_LGIC_rGluNMDA_Agonist was analyzed into 1 assay endpoint. This assay endpoint, NVS_LGIC_rGluNMDA_Agonist, was analyzed in the positive fitting direction relative to DMSO as the negative control and baseline of activity. Using a type of binding reporter, loss-of-signal activity can be used to understand changes in the binding as they relate to the gene Grin1.  Furthermore, this assay endpoint can be referred to as a primary readout, because the performed assay has only produced 1 assay endpoint. To generalize the intended target to other relatable targets, this assay endpoint is annotated to the ""ion channel"" intended target family, where the subfamily is "ligand-gated ion channel". | Data from the assay component NVS_LGIC_rGluNMDA_MK801_Agonist was analyzed into 1 assay endpoint. This assay endpoint, NVS_LGIC_rGluNMDA_MK801_Agonist, was analyzed in the positive fitting direction relative to DMSO as the negative control and baseline of activity. Using a type of binding reporter, loss-of-signal activity can be used to understand changes in the binding as they relate to the gene Grin1. Furthermore, this assay endpoint can be referred to as a primary readout, because the performed assay has only produced 1 assay endpoint. To generalize the intended target to other relatable targets, this assay endpoint is annotated to the "ion channel" intended target family, where the subfamily is "ligand-gated ion channel". |
| Assay Function Type | binding | binding |
| Normalized Data Type | percent_activity | percent_activity |
| Analysis Direction | positive | positive |
| Burst Assay | 0 | 0 |
| Key Positive Control | N-methyl-D-aspartate (NMDA) | (+)-MK-801 |
| Signal Direction | loss | loss |
| Intended Target Type | protein | protein |
| Intended Target Type Sub | receptor | receptor |
| Intended Target Family | ion channel | ion channel |
| Intended Target Family Sub | ligand-gated ion channel | ligand-gated ion channel |
| Assay Component Name | NVS_LGIC_rGluNMDA_Agonist | NVS_LGIC_rGluNMDA_MK801_Agonist |
| Assay Component Desc | NVS_LGIC_rGluNMDA_Agonist, is one of one assay component(s) measured or calculated from the NVS_LGIC_rGluNMDA_Agonist assay. It is designed to make measurements of radioligand binding, a form of binding reporter, as detected with scintillation counting signals by Filter-based radiodetection technology. | NVS_LGIC_rGluNMDA_MK801_Agonist, is one of one assay component(s) measured or calculated from the NVS_LGIC_rGluNMDA_MK801_Agonist assay. It is designed to make measurements of radioligand binding, a form of binding reporter, as detected with scintillation counting signals by Filter-based radiodetection technology. |
| Assay Component Target Desc | Changes to scintillation counting signals produced from the receptor-ligand binding of the key ligand [[3H]-CGP 39653] are indicative of a change in receptor agonist activity for the Norway rat glutamate receptor, ionotropic, N-methyl D-aspartate 1 [GeneSymbol:Grin1 \| GeneID:24408 \| Uniprot_SwissProt_Accession:P35439]. | Changes to scintillation counting signals produced from the receptor-ligand binding of the key ligand [[3H]-MK-801] are indicative of a change in receptor agonist activity for the Norway rat glutamate receptor, ionotropic, N-methyl D-aspartate 1 [GeneSymbol:Grin1 \| GeneID:24408 \| Uniprot_SwissProt_Accession:P35439]. |
| Parameter Readout Type | single | single |
| Assay Design Type | binding reporter | binding reporter |
| Assay Design Type Sub | radioligand binding | radioligand binding |
| Biological Process Target | receptor binding | receptor binding |
| Detection Technology Type | Filter-based radiodetection | Filter-based radiodetection |
| Detection Technology Type Sub | Scintillation counting | Scintillation counting |
| Detection Technology | Radiometry | Radiometry |
| Signal Direction Type | loss | loss |
| Key Assay Reagent Type | ligand | ligand |
| Key Assay Reagent | [3H]-CGP 39653 | [3H]-MK-801 |
| Technological Target Type | protein | protein |
| Technological Target Type Sub | receptor | receptor |
| Assay Name | NVS_LGIC_rGluNMDA_Agonist | NVS_LGIC_rGluNMDA_MK801_Agonist |
| Assay Desc | NVS_LGIC_rGluNMDA_Agonist is a biochemical, single-readout assay that uses extracted gene-proteins from Rat forebrain membranes in a tissue-based cell-free assay. Measurements were taken 1 hour after chemical dosing in a 48-well plate. | NVS_LGIC_rGluNMDA_MK801_Agonist is a biochemical, single-readout assay that uses extracted gene-proteins from Rat forebrain membranes in a tissue-based cell-free assay. Measurements were taken 1.5 hours after chemical dosing in a 48-well plate. |
| Timepoint Hr | 1 | 1.5 |
| Organism | rat | rat |
| Tissue | brain | brain |
| Cell Format | tissue-based cell-free | tissue-based cell-free |
| Cell Free Component Source | Rat forebrain membranes | Rat forebrain membranes |
| Cell Short Name | NA | NA |
| Cell Growth Mode | NA | NA |
| Assay Footprint | microplate: 48-well plate | microplate: 48-well plate |
| Assay Format Type | biochemical | biochemical |
| Assay Format Type Sub | protein single format | protein single format |
| Content Readout Type | single | single |
| Dilution Solvent | DMSO | DMSO |
| Dilution Solvent Percent Max | 0 | 0 |
| Assay Source Name | NVS | NVS |
| Assay Source Desc |  |  |
| Entrez Gene Id | 24408 | 24408 |
| Gene Name | glutamate receptor, ionotropic, N-methyl D-aspartate 1 | glutamate receptor, ionotropic, N-methyl D-aspartate 1 |
| Gene Symbol | Grin1 | Grin1 |

**Table S5. Active and inactive chemicals determined for the assay components NVS_LGIC_rGluNMDA_Agonist and NVS_LGIC_rGluNMDA_MK801_Agonist.** Data gathered from the EPA CompTox Chemicals Dashboard (https://comptox.epa.gov/dashboard/).

| **Chemical** | **DTXSID** | **NVS_LGIC_rGluNMDA_Agonist** | | **NVS_LGIC_rGluNMDA_MK801_Agonist** | |
| --- | --- | --- | --- | --- | --- |
|  |  | **Hit call** | **AC50 (µM)** | **Hit call** | **AC50 (µM)** |
| (2S,3S)-3-Methyl-2-(3-oxo-1,2-benzothiazol-2(3H)-yl)pentanoic acid | DTXSID3047261 | 0 | NA | NA | NA |
| 1,3,5-Trimethylbenzene | DTXSID6026797 | NA | NA | 0 | NA |
| 1,3-Diphenylguanidine | DTXSID3025178 | 1 | 4.45 | 0 | NA |
| 1-Benzylquinolinium chloride | DTXSID8044593 | NA | NA | 0 | 26.57 |
| 2,4,6-Trimethylphenol | DTXSID7022049 | 0 | 0.49 | NA | NA |
| 4,4',4''-Ethane-1,1,1-triyltriphenol | DTXSID2037712 | 0 | NA | NA | NA |
| 4,4'-Sulfonylbis[2-(prop-2-en-1-yl)phenol] | DTXSID9047598 | NA | NA | 0 | NA |
| AVE5638 | DTXSID0047371 | NA | NA | 1 | 35.48 |
| Abamectin | DTXSID8023892 | NA | NA | 0 | 7.56 |
| Alachlor | DTXSID1022265 | 0 | NA | NA | NA |
| Aldicarb oxime | DTXSID9024431 | 0 | NA | NA | NA |
| Aminopterin | DTXSID3022588 | 0 | NA | NA | NA |
| Azoxystrobin | DTXSID0032520 | NA | NA | 0 | 20.94 |
| Benodanil | DTXSID7041623 | 0 | NA | NA | NA |
| Bentazone | DTXSID0023901 | 0 | NA | NA | NA |
| Besonprodil | DTXSID2047270 | 1 | 0.06 | 0 | 0.3 |
| Bisphenol A | DTXSID7020182 | 1 | 17.49 | NA | NA |
| Bisphenol AF | DTXSID7037717 | 1 | 19.92 | 1 | 25.1 |
| Bisphenol B | DTXSID4022442 | 1 | 17.99 | NA | NA |
| Butralin | DTXSID3032337 | 0 | NA | NA | NA |
| CP-114271 | DTXSID2047274 | 0 | NA | NA | NA |
| CP-283097 | DTXSID8047264 | 0 | NA | 0 | NA |
| Cariporide mesylate | DTXSID3047344 | NA | NA | 0 | 10.63 |
| Chlorpromazine hydrochloride | DTXSID7024827 | 1 | 22.18 | 1 | 35.21 |
| Clofibrate | DTXSID3020336 | NA | NA | 0 | NA |
| Clorophene | DTXSID5020154 | 1 | 7.59 | NA | NA |
| Cycloate | DTXSID6032356 | 0 | NA | NA | NA |
| Cyclohexylamine | DTXSID1023996 | NA | NA | 0 | NA |
| Cypermethrin | DTXSID1023998 | 0 | NA | NA | NA |
| Decane | DTXSID6024913 | 0 | NA | NA | NA |
| Di(2-ethylhexyl) phthalate | DTXSID5020607 | 0 | NA | NA | NA |
| Difpas-pyrazole | DTXSID6048175 | 0 | 5.16 | NA | NA |
| Diphenhydramine hydrochloride | DTXSID4020537 | NA | NA | 1 | 15.47 |
| Docusate sodium | DTXSID8022959 | 1 | 31.87 | 0 | NA |
| Dodecylbenzene sulfonate triethanolamine(1:1) | DTXSID5027932 | 0 | NA | NA | NA |
| Dodecylbenzenesulfonic acid | DTXSID6027923 | 1 | 31.37 | NA | NA |
| Emamectin benzoate | DTXSID0034566 | 1 | 19.56 | NA | NA |
| Ethanolamine | DTXSID6022000 | 0 | NA | NA | NA |
| Ethephon | DTXSID7024085 | NA | NA | 0 | NA |
| Ethylene thiourea | DTXSID5020601 | 0 | NA | 0 | NA |
| HMR1171 | DTXSID6047367 | NA | NA | 0 | NA |
| Haloperidol | DTXSID4034150 | NA | NA | 0 | 2.7 |
| Hexachlorocyclopentadiene | DTXSID2020688 | NA | NA | 0 | NA |
| Hexaconazole | DTXSID4034653 | NA | NA | 0 | 19.68 |
| Imazalil | DTXSID8024151 | NA | NA | 1 | 19.41 |
| Isoxaben | DTXSID8024159 | 0 | NA | NA | NA |
| MEHP | DTXSID2025680 | 0 | NA | NA | NA |
| MK-968 | DTXSID9047334 | 1 | 0.6 | 0 | NA |
| Mancozeb | DTXSID0034695 | 1 | 9.5 | 1 | 28.65 |
| Maneb | DTXSID9020794 | 1 | 18.39 | 1 | 41.01 |
| Metam-sodium hydrate | DTXSID2040361 | 0 | NA | 0 | NA |
| Methadone hydrochloride | DTXSID2020501 | NA | NA | 1 | 8.4 |
| Methyl isothiocyanate | DTXSID2027204 | 0 | 0.4 | NA | NA |
| Methyl salicylate | DTXSID5025659 | 0 | NA | NA | NA |
| Metiram | DTXSID9034737 | 0 | NA | 1 | 18.3 |
| Metolachlor | DTXSID4022448 | NA | NA | 0 | NA |
| Milbemectin (mixture of 70% Milbemcin A4, 30% Milbemycin A3) | DTXSID8034742 | 1 | 25.68 | 1 | 23.22 |
| N-Nitrosodiphenylamine | DTXSID6021030 | NA | NA | 0 | 1.63 |
| Octanal | DTXSID3021643 | 0 | NA | 0 | NA |
| Octhilinone | DTXSID1025805 | 0 | NA | NA | NA |
| Paclobutrazol | DTXSID2024242 | 0 | 2.31 | NA | NA |
| Pentamidine isethionate | DTXSID5023796 | NA | NA | 1 | 24.34 |
| PharmaGSID_47263 | DTXSID3047263 | 0 | NA | NA | NA |
| PharmaGSID_47315 | DTXSID6047315 | NA | NA | 0 | NA |
| PharmaGSID_48511 | DTXSID4048511 | 1 | 15.07 | 1 | 20.63 |
| Potassium perfluorooctanesulfonate | DTXSID8037706 | 0 | NA | NA | NA |
| Propargite | DTXSID4024276 | 0 | NA | NA | NA |
| Prosulfuron | DTXSID9034868 | 0 | NA | NA | NA |
| SB236057A | DTXSID5047320 | NA | NA | 1 | 40.98 |
| SSR 103800 | DTXSID1047364 | NA | NA | 0 | NA |
| SSR 240612 | DTXSID2047351 | NA | NA | 0 | NA |
| SSR 241586 HCl | DTXSID2047353 | 1 | 31.55 | 0 | NA |
| Sodium L-ascorbate | DTXSID0020105 | 0 | 4.57 | NA | NA |
| Sodium dodecyl sulfate | DTXSID1026031 | NA | NA | 0 | NA |
| Sodium dodecylbenzenesulfonate | DTXSID7025219 | 1 | 158.11 | 0 | NA |
| Spironolactone | DTXSID6034186 | 1 | 14.35 | NA | NA |
| Tamoxifen citrate | DTXSID8021301 | NA | NA | 0 | NA |
| UK-337312 | DTXSID6047288 | 1 | 27.06 | NA | NA |
| Urea | DTXSID4021426 | 0 | NA | NA | NA |
| Zamifenacin | DTXSID9047257 | NA | NA | 0 | NA |
| Zenarestat | DTXSID0047296 | 0 | 33.87 | NA | NA |

**Table S6. Cytotoxicity limits and estimated acute toxicity to fish.** The cytotoxicity limit represents the lower bound of cytotoxicity, based on data from the EPA CompTox database, as described in the Materials and Methods section. LC50 represents the predicted 96-h LC_50_ in fish (ECOSAR v2.0, U.S. EPA).

| **Chemical** | **Cytotoxicity limit (µM)** | **LC50_1 (µM)** | **Organic Module Result_1** | **LC50_2 (µM)** | **Organic Module Result_2** |
| --- | --- | --- | --- | --- | --- |
| 1,3-Diphenylguanidine | 4.27 | 49.7 | Aliphatic Amines | NA | NA |
| AVE5638 | 11.65 | 7.4 | Aliphatic Amines | 5.84 | Amides |
| Besonprodil | 9.3 | 7.83 | Aliphatic Amines | 4.5 | Carbamate Esters, Phenyl |
| Bisphenol A | 15.68 | 5.61 | Phenols, Poly | NA | NA |
| Bisphenol AF | 14.01 | 1.8 | Phenols, Poly | NA | NA |
| Bisphenol B | 13.56 | 2.87 | Phenols, Poly | NA | NA |
| Chlorpromazine hydrochloride | 10.57 | 25.08 | Neutral Organics | NA | NA |
| Clorophene | 10.92 | 4.32 | Phenols | NA | NA |
| Diphenhydramine hydrochloride | 12.47 | 1922.42 | Neutral Organics | NA | NA |
| Docusate sodium | 9.47 | 5.42 | Esters | NA | NA |
| Dodecylbenzenesulfonic acid | 11.72 | 25.94 | Neutral Organics | NA | NA |
| Emamectin benzoate | 2.8 | NA | NA | NA | NA |
| Imazalil | 12.5 | 8.34 | Vinyl/Allyl/Propargyl Ketones | 0.45 | Pyrroles/Diazoles |
| Mancozeb | 0.63 | 25.29 | Thiocarbamate,Di (Free Acid) | NA | NA |
| Maneb | 1.58 | 25.29 | Thiocarbamate,Di (Free Acid) | NA | NA |
| Methadone hydrochloride | 9.88 | 2847.48 | Neutral Organics | NA | NA |
| Metiram | 11.72 | 3.63 | Thiocarbamate,Di (Substituted) | 45.23 | Thiocarbamate,Di (Free Acid) |
| Milbemectin (mixture of 70% Milbemcin A4, 30% Milbemycin A3) | 8.72 | NA | NA | NA | NA |
| MK-968 | 12.78 | 17.11 | Aliphatic Amines | 0.7 | Hydrazines |
| Pentamidine isethionate | 6.3 | 698.21 | Aliphatic Amines | NA | NA |
| PharmaGSID_48511 | 7.36 | 8.22 | Aliphatic Amines | 6.62 | Amides |
| SB236057A | 9.8 | NA | NA | NA | NA |
| Sodium dodecylbenzenesulfonate | 10.84 | 25.94 | Neutral Organics | NA | NA |
| Spironolactone | 11.72 | 41.29 | Esters | 218.45 | Vinyl/Allyl/Propargyl Ketones |
| SSR 241586 HCl | 9.12 | NA | NA | NA | NA |
| UK-337312 | 8.48 | NA | NA | NA | NA |

**Table S7. Similarity Ensemble Approach**^5^ **target predictions for MK-801.** TC, Tanimoto coefficient (maximum pairwise similarities between query molecule and sets of ligands).

| **Target Key** | **Target Name** | **Description** | **P-Value** | **MaxTC** |
| --- | --- | --- | --- | --- |
| NMDZ1_RAT | Grin1 | Glutamate receptor ionotropic, NMDA 1 | 4.163E-14 | 1.00 |
| NMDE1_HUMAN | GRIN2A | Glutamate receptor ionotropic, NMDA 2A | 2.207E-06 | 1.00 |
| PCP_HUMAN | PRCP | Lysosomal Pro-X carboxypeptidase | 4.352E-06 | 1.00 |
| ACM5_HUMAN | CHRM5 | Muscarinic acetylcholine receptor M5 | 0.06101 | 1.00 |
| CP2D6_HUMAN | CYP2D6 | Cytochrome P450 2D6 | 0.4944 | 1.00 |
| GBRA2_RAT | Gabra2 | Gamma-aminobutyric acid receptor subunit alpha-2 | 7.524E-08 | 0.63 |
| MCHR2_HUMAN | MCHR2 | Melanin-concentrating hormone receptor 2 | 9.984E-07 | 0.28 |
| SGMR1_RAT | Sigmar1 | Sigma non-opioid intracellular receptor 1 | 3.919E-06 | 0.63 |
| ACHP_LYMST | NA | Acetylcholine-binding protein | 5.589E-06 | 0.30 |

**Table S8. Similarity Ensemble Approach**^5^ **target predictions for chlorophene.** TC, Tanimoto coefficient (maximum pairwise similarities between query molecule and set of ligands).

| **Target Key** | **Target Name** | **Description** | **P-Value** | **MaxTC** |
| --- | --- | --- | --- | --- |
| Q965D5_PLAFA | fabI | Enoyl-acyl-carrier protein reductase | 4.74E-62 | 0.49 |
| LAT1_HUMAN | SLC7A5 | Large neutral amino acids transporter small subunit 1 | 1.48E-46 | 0.50 |
| MGRA_STAAU | mgrA | HTH-type transcriptional regulator MgrA | 6.68E-38 | 0.32 |
| C5ISA2_SHEEP | TUBA4A | Tubulin alpha chain | 2.85e-39 | 0.31 |
| HNMT_HUMAN | HNMT | Histamine N-methyltransferase | 5.06e-39 | 0.31 |
| ERCC1_HUMAN | ERCC1 | DNA excision repair protein ERCC-1 | 3.23e-37 | 0.36 |
| XPF_HUMAN | ERCC4 | DNA repair endonuclease XPF | 3.23e-37 | 0.36 |
| KCMA1_HUMAN | KCNMA1 | Calcium-activated potassium channel subunit alpha-1 | 4.65e-36 | 0.57 |
| TCMO_HELTU | CYP73A1 | Trans-cinnamate 4-monooxygenase | 1.10E-32 | 0.28 |
| Q6UCJ9_TOXGO | ENR | Enoyl-acyl carrier reductase | 2.46E-27 | 0.41 |
| PE2R1_HUMAN | PTGER1 | Prostaglandin E2 receptor EP1 subtype | 1.92E-25 | 0.43 |
| IBP5_HUMAN | IGFBP5 | Insulin-like growth factor-binding protein 5 | 5.13e-27 | 0.31 |
| SYUA_HUMAN | SNCA | Alpha-synuclein | 3.73E-22 | 0.37 |
| ARP19_RAT | Arpp19 | cAMP-regulated phosphoprotein 19 | 7.91E-22 | 0.31 |
| CTBP2_HUMAN | CTBP2 | C-terminal-binding protein 2 | 5.13E-20 | 0.38 |
| LOX5_RAT | Alox5 | Arachidonate 5-lipoxygenase | 1.13E-18 | 0.56 |
| EGFR_MOUSE | Egfr | Epidermal growth factor receptor | 1.90E-17 | 0.31 |
| Q9BJJ9_PLAFA | FabI | Enoyl-ACP reductase | 7.56E-17 | 0.32 |
| M9TGV3_MYCTX | inhA | Enoyl-[acyl-carrier-protein] reductase [NADH] | 1.25E-15 | 0.49 |
| PERM_HUMAN | MPO | Myeloperoxidase | 4.01E-14 | 0.38 |
| FTSZ_ECOLI | ftsZ | Cell division protein FtsZ | 2.22e-16 | 0.31 |
| UPP1_MOUSE | Upp1 | Uridine phosphorylase 1 | 2.22e-16 | 0.37 |
| Q8Z9J9_SALTI | folA | Dihydrofolate reductase | 5.55E-13 | 0.34 |
| Q49PX0_9POXV | N1L | N1L | 7.77E-13 | 0.59 |
| PGH2_RAT | Ptgs2 | Prostaglandin G/H synthase 2 | 1.22E-12 | 0.33 |
| Q3I4V7_CRYNV | CAN2 | Carbonic anhydrase | 4.33e-15 | 0.40 |
| HCAR2_RAT | Hcar2 | Hydroxycarboxylic acid receptor 2 | 1.88E-11 | 0.35 |
| PGH1_RAT | Ptgs1 | Prostaglandin G/H synthase 1 | 2.74E-11 | 0.31 |
| FABI_BACSU | fabI | Enoyl-[acyl-carrier-protein] reductase [NADH] FabI | 3.42E-11 | 0.32 |
| THB_RAT | Thrb | Thyroid hormone receptor beta | 4.23E-10 | 0.41 |
| Q5NGQ3_FRATT | fabI | Enoyl-[acyl-carrier-protein] reductase [NADH] | 6.86E-10 | 0.32 |
| B2LA1_HUMAN | BCL2A1 | Bcl-2-related protein A1 | 1.06E-09 | 0.34 |
| TTHY_HUMAN | TTR | Transthyretin | 1.15E-09 | 0.33 |
| OXDA_HUMAN | DAO | D-amino-acid oxidase | 1.39E-09 | 0.32 |
| S6A12_HUMAN | SLC6A12 | Sodium- and chloride-dependent betaine transporter | 1.4e-12 | 0.31 |
| PD2R2_MOUSE | Ptgdr2 | Prostaglandin D2 receptor 2 | 1.42E-09 | 0.30 |
| Q7ZJM1_9HIV1 | pol | Integrase | 7.35E-09 | 0.36 |
| GBRA3_RAT | Gabra3 | Gamma-aminobutyric acid receptor subunit alpha-3 | 8.66E-09 | 0.30 |
| THA_HUMAN | THRA | Thyroid hormone receptor alpha | 1.01E-08 | 0.34 |
| POLG_CXB3N | NA | Genome polyprotein | 2.55E-08 | 0.33 |
| LDHB_HUMAN | LDHB | L-lactate dehydrogenase B chain | 3.90E-08 | 0.35 |
| CBS_HUMAN | CBS | Cystathionine beta-synthase | 8.58E-08 | 0.29 |
| PPO1_AGABI | PPO1 | Polyphenol oxidase 1 | 1.44E-07 | 0.33 |
| FEN1_HUMAN | FEN1 | Flap endonuclease 1 | 1.75E-07 | 0.29 |
| S100B_HUMAN | S100B | Protein S100-B | 2.66E-07 | 0.30 |
| THB_HUMAN | THRB | Thyroid hormone receptor beta | 3.10E-07 | 0.34 |
| RNH1_HUMAN | RNASEH1 | Ribonuclease H1 | 9.46e-10 | 0.28 |
| POLG_HCV1 | NA | Genome polyprotein | 1.63E-06 | 0.33 |
| PDS_SYNE7 | pds | 15-cis-phytoene desaturase | 1.63E-06 | 0.30 |
| TAAR1_HUMAN | TAAR1 | Trace amine-associated receptor 1 | 1.76E-06 | 0.46 |
| CAN_CANAL | NCE103 | Carbonic anhydrase | 1.77E-06 | 0.40 |
| LX15B_RAT | Alox15b | Arachidonate 15-lipoxygenase B | 2.90E-06 | 0.29 |
| ERCC5_HUMAN | ERCC5 | DNA repair protein complementing XP-G cells | 3.02E-06 | 0.29 |
| AMD_HUMAN | PAM | Peptidyl-glycine alpha-amidating monooxygenase | 4.61E-06 | 0.28 |
| LYAM1_HUMAN | SELL | L-selectin | 7.08e-09 | 0.29 |
| TYTR_TRYBB | TPR | Trypanothione reductase | 1.00E-05 | 0.30 |
| O77078_PLAFA | fabH | Beta-ketoacyl-acyl carrier protein synthase III | 1.08E-05 | 0.29 |
| RGS8_HUMAN | RGS8 | Regulator of G-protein signaling 8 | 1.08E-05 | 0.29 |
| I23O1_MOUSE | Ido1 | Indoleamine 2,3-dioxygenase 1 | 1.12E-05 | 0.35 |
| DOPO_HUMAN | DBH | Dopamine beta-hydroxylase | 1.31E-05 | 0.32 |
| TBB2B_BOVIN | TUBB2B | Tubulin beta-2B chain | 1.44E-05 | 0.31 |
| LOX15_RAT | Alox15 | Arachidonate 15-lipoxygenase | 1.54e-08 | 0.29 |
| CBR1_HUMAN | CBR1 | Carbonyl reductase [NADPH] 1 | 1.91E-05 | 0.32 |
| NMDE2_HUMAN | GRIN2B | Glutamate receptor ionotropic, NMDA 2B | 2.26E-05 | 0.37 |
| LOX15_HUMAN | ALOX15 | Arachidonate 15-lipoxygenase | 2.85E-05 | 0.39 |
| AK1BA_HUMAN | AKR1B10 | Aldo-keto reductase family 1 member B10 | 3.22E-05 | 0.30 |
| ACPS_BACSU | acpS | Holo-[acyl-carrier-protein] synthase | 6.21E-05 | 0.33 |
| G6PC_HUMAN | G6PC | Glucose-6-phosphatase | 6.67E-05 | 0.31 |
| POLG_DEN2P | NA | Genome polyprotein | 1.45E-04 | 0.32 |
| GBRA2_RAT | Gabra2 | Gamma-aminobutyric acid receptor subunit alpha-2 | 1.91E-04 | 0.30 |
| CISY2_YEAST | CIT2 | Citrate synthase, peroxisomal | 1.91e-07 | 0.30 |
| LUXP_VIBHA | luxP | Autoinducer 2-binding periplasmic protein LuxP | 2.19E-04 | 0.32 |
| LYAM3_HUMAN | SELP | P-selectin | 2.23E-04 | 0.30 |
| RGS4_HUMAN | RGS4 | Regulator of G-protein signaling 4 | 2.46E-04 | 0.30 |
| A0A0C7ACN7_PSEAI | pqsD | 3-oxoacyl-ACP synthase | 2.75E-04 | 0.31 |
| Q8MTZ4_9TRYP | PPase1 | Vacuolar-type proton translocating pyrophosphatase 1 | 2.97E-04 | 0.31 |
| KCNQ2_MOUSE | Kcnq2 | Potassium voltage-gated channel subfamily KQT member 2 | 4.00E-04 | 0.32 |
| PD2R2_HUMAN | PTGDR2 | Prostaglandin D2 receptor 2 | 4.36E-04 | 0.35 |
| AK1C4_HUMAN | AKR1C4 | Aldo-keto reductase family 1 member C4 | 4.91E-04 | 0.29 |
| CAH3_BOVIN | CA3 | Carbonic anhydrase 3 | 5.63E-04 | 0.30 |
| VKOR1_RAT | Vkorc1 | Vitamin K epoxide reductase complex subunit 1 | 6.32E-04 | 0.28 |
| KDM4E_HUMAN | KDM4E | Lysine-specific demethylase 4E | 6.33e-07 | 0.31 |
| GBRA5_RAT | Gabra5 | Gamma-aminobutyric acid receptor subunit alpha-5 | 8.26E-04 | 0.30 |
| GBRB3_RAT | Gabrb3 | Gamma-aminobutyric acid receptor subunit beta-3 | 1.02E-03 | 0.30 |
| SHBG_HUMAN | SHBG | Sex hormone-binding globulin | 1.06E-03 | 0.32 |
| C3TDZ2_ECOLX | fabH | 3-oxoacyl-[acyl-carrier-protein] synthase 3 | 1.13E-03 | 0.35 |
| GPR18_HUMAN | GPR18 | N-arachidonyl glycine receptor | 1.29E-03 | 0.30 |
| SGMR2_HUMAN | TMEM97 | Sigma intracellular receptor 2 | 1.68E-03 | 0.29 |
| CA2D1_MOUSE | Cacna2d1 | Voltage-dependent calcium channel subunit alpha-2/delta-1 | 1.89E-03 | 0.29 |
| COMT_RAT | Comt | Catechol O-methyltransferase | 1.98E-03 | 0.33 |
| KDM5B_HUMAN | KDM5B | Lysine-specific demethylase 5B | 2.07e-06 | 0.31 |
| AL1A1_HUMAN | ALDH1A1 | Retinal dehydrogenase 1 | 2.09E-03 | 0.35 |
| FABH_MYCTU | fabH | 3-oxoacyl-[acyl-carrier-protein] synthase 3 | 2.47E-03 | 0.29 |
| PPBI_MOUSE | Iap | Intestinal-type alkaline phosphatase | 2.92E-03 | 0.33 |
| I23O1_HUMAN | IDO1 | Indoleamine 2,3-dioxygenase 1 | 3.14E-03 | 0.35 |
| NQO1_HUMAN | NQO1 | NAD(P)H dehydrogenase [quinone] 1 | 3.65e-06 | 0.33 |
| P96878_MYCTU | NA | Probable transmembrane carbonic anhydrase (Carbonate dehydratase) (Carbonic dehydratase) | 4.33E-03 | 0.40 |
| BLA2_BACCE | blm | Metallo-beta-lactamase type 2 | 4.4e-06 | 0.30 |
| PYGL_HUMAN | PYGL | Glycogen phosphorylase, liver form | 4.80E-03 | 0.30 |
| PNPH_RAT | Pnp | Purine nucleoside phosphorylase | 5.25E-03 | 0.28 |
| KIF11_HUMAN | KIF11 | Kinesin-like protein KIF11 | 5.29E-03 | 0.32 |
| ENPL_CANLF | HSP90B1 | Endoplasmin | 6.14E-03 | 0.32 |
| LDHA_HUMAN | LDHA | L-lactate dehydrogenase A chain | 6.81E-03 | 0.35 |
| DYR_LACCA | folA | Dihydrofolate reductase | 7.69E-03 | 0.34 |
| Q76353_9HIV1 | NA | Integrase | 7.72E-03 | 0.36 |
| GP183_MOUSE | Gpr183 | G-protein coupled receptor 183 | 7.88E-03 | 0.31 |
| VEGFA_HUMAN | VEGFA | Vascular endothelial growth factor A | 9.43e-06 | 0.32 |

### 2 Supplemental Figures

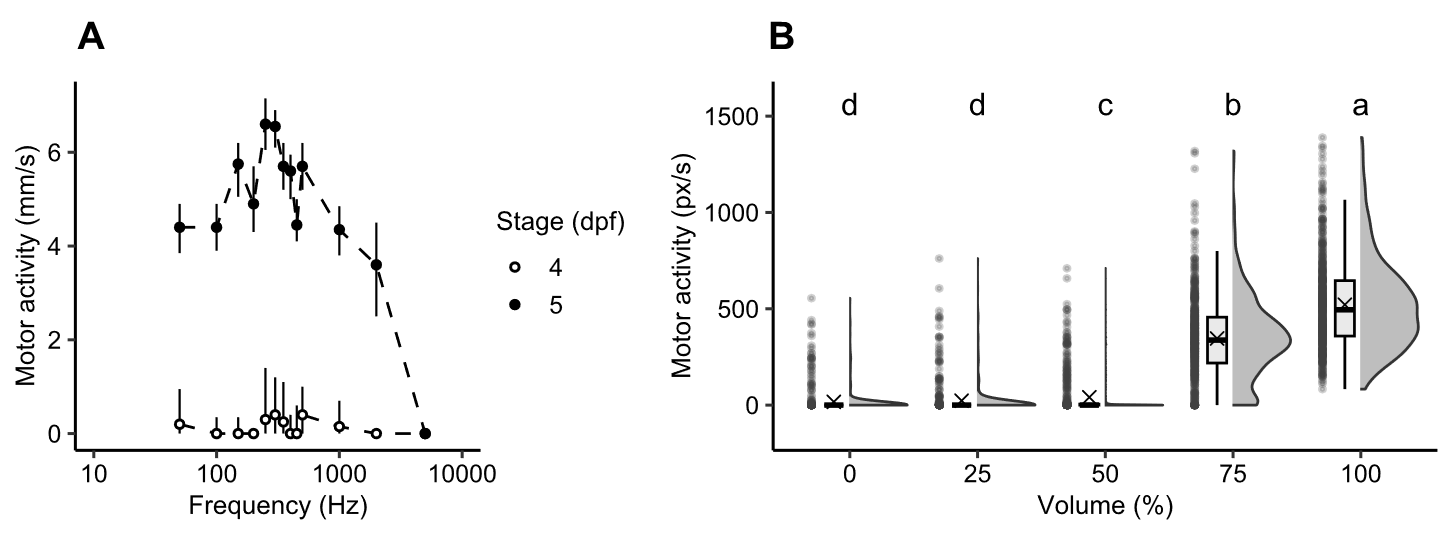

**Figure S1. Developmental stage-, frequency-, and volume-dependency of acoustic startle response and acoustic startle habituation in larval zebrafish.** Parameters considered for establishment of behavior assay battery components based on acoustic stimuli. (**A**) Comparison of responsiveness (y-axis) to 1-second acoustic stimuli (100% volume, 0% light intensity) across different frequencies (x-axis) in 4 and 5 d post-fertilization (dpf) zebrafish larvae. Data shows median ± 95% CI (n=192 larvae). (**B**) Acoustic startle response intensity (y-axis) at different nominal volume levels (x-axis) at 5 dpf. Larvae were dark-acclimated for 20 min prior to application of five consecutive acoustic stimuli (1 s, 300 Hz, inter stimulus interval (ISI) 60 s). Acoustic stimulation bouts per volume level were interspaced with inter bout intervals of 5 min. Data points represent average per volume level and individual across five trials. Compact letter display above rainclouds indicates significant differences between volume levels (p<0.05, Kruskal-Wallis rank sum test, n = 72 larvae). Summary data can be found in Excel Tables S32-S33.

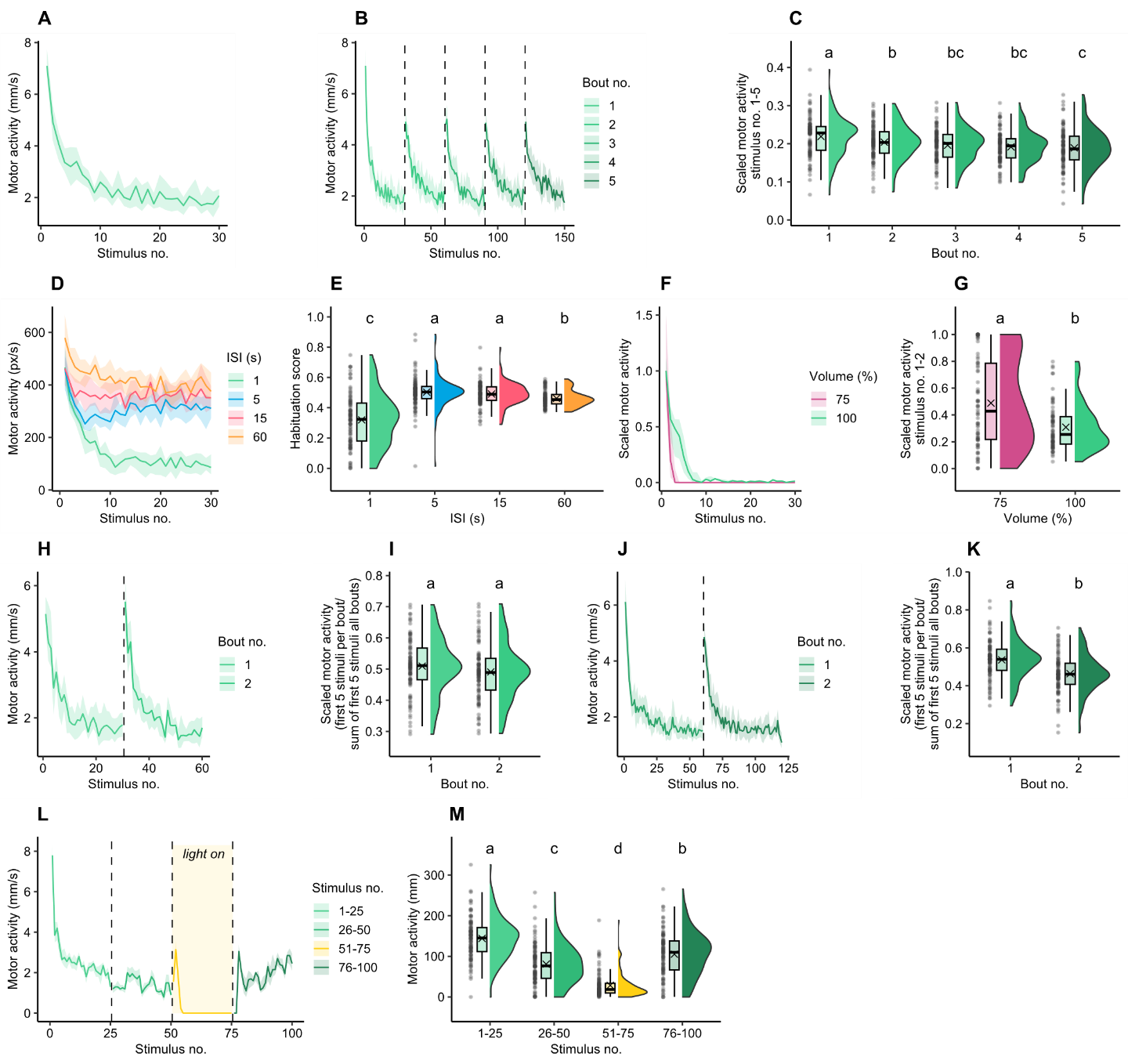

**Figure S2. Representative results from this study demonstrating characteristics of habituation as described by Rankin et al.**^1^**.** (**A**) Progressive response decrease to an asymptotic level. Motor activity (y-axis) of 5-dpf zebrafish larvae over a sequence of 30 consecutive 1-second acoustic stimuli (300 Hz, 100% volume, inter stimulus interval (ISI) 1 s). Data is median ± 95% CI (n = 96). (**B, C**) Spontaneous recovery of response after response decrement. Stimulus repetitions and spontaneous recoveries result in more rapid/pronounced habituation (potentiation of habituation). Line graph shows the motor activity (y-axis) of 5-dpf zebrafish larvae over a sequence of five habituation bouts, each consisting of 30 consecutive 1-second acoustic stimuli (x-axis) (300 Hz, 100% volume, ISI = 1 s). Dashed vertical lines indicate inter-bout intervals (IBIs) of 1 min. Data is median ± 95% CI (n = 96). Raincloud graph shows the cumulative motor activity during the first five stimuli per habituation bout scaled to the overall cumulative motor activity per individual across the first five stimuli (y-axis) of each habituation bout (x-axis) shown in B. Compact letter display above rainclouds indicates significant reduction in motor activity across habituation bouts (p<0.05, Kruskal-Wallis rank sum test). (**D, E**) More frequent stimulation results in more rapid/pronounced response decrement. Line graph shows the motor activity (y-axis) of 5-dpf zebrafish larvae over a sequence of 30 consecutive 1-second acoustic stimuli (300 Hz, 100% volume; x-axis) with different ISIs. Data is median ± 95% CI (n = 96). Raincloud graph shows the habituation scores (y-axis) for different ISIs (x-axis) shown in **D**. Habituation scores represent the cumulative motor activity per individual during stimuli 21-30 relative to the cumulative motor activity summed across stimuli 1-10 and 21-30. Compact letter display above rainclouds indicates significant differences between ISIs (p<0.05, Kruskal-Wallis rank sum test). (**F, G**) Less intense stimuli result in more rapid/pronounced response decrement. Line graph shows the motor activity (y-axis) of 5-dpf zebrafish larvae over a sequence of 30 consecutive 1-second acoustic stimuli (300 Hz, ISI = 1 s; x-axis) at different nominal volume levels. Motor activity was scaled to the median response to the first acoustic stimulus per volume group to account for differences in baseline startle activity. Individuals that did not react to any of the 30 acoustic stimuli, were excluded. Data is median ± 95% CI (n = 74). Raincloud graph shows the cumulative motor activity during the first two stimuli scaled to the overall activity across all 30 stimuli (y-axis) for two different volume levels (x-axis) shown in **F**. Compact letter display above rainclouds indicates significant differences between volume levels (p<0.05, Kruskal-Wallis rank sum test). (**H-K**) Accumulation of habituation beyond asymptotic level. Line graphs in **H** and **J** show the motor activity of 5-dpf zebrafish larvae (y-axis) over two subsequent habituation bouts, each consisting of a sequence of 30 and 60 consecutive 1-second acoustic stimuli (300 Hz, 100% volume, ISI = 1 s; x-axis), respectively. Data is median ± 95% CI (n = 96). Raincloud graphs in **I** and **K** show the cumulative motor activity during the first five stimuli per habituation bout scaled to the overall motor activity during the first five stimuli per bout (y-axis) for two subsequent bouts (x-axis), each comprising of 30 and 60 acoustic stimuli shown in **H** and **J**, respectively. Compact letter display above rainclouds indicates significant differences between bouts (p<0.05, Kruskal-Wallis rank sum test). (**L, M**) A second, novel stimulus changes the response to the habituated stimulus (stimulus generalization/specificity). A different stimulus increases the decremented response to the original stimulus (dishabituation). Line graph shows the motor activity (y-axis) of 5-dpf zebrafish larvae over a sequence of 100 consecutive 1-second acoustic stimuli (300 Hz, 100% volume, ISI = 1 s; x-axis). Light was turned on during stimuli 51-75. Data is median ± 95% CI (n = 96). Raincloud graph shows the cumulative motor activity per 25-stimuli bin (indicated by dashed vertical lines in **L**). Compact letter display above rainclouds indicates significant differences between bins (p<0.05, Kruskal-Wallis rank sum test). Summary data can be found in Excel Tables S34-S46.

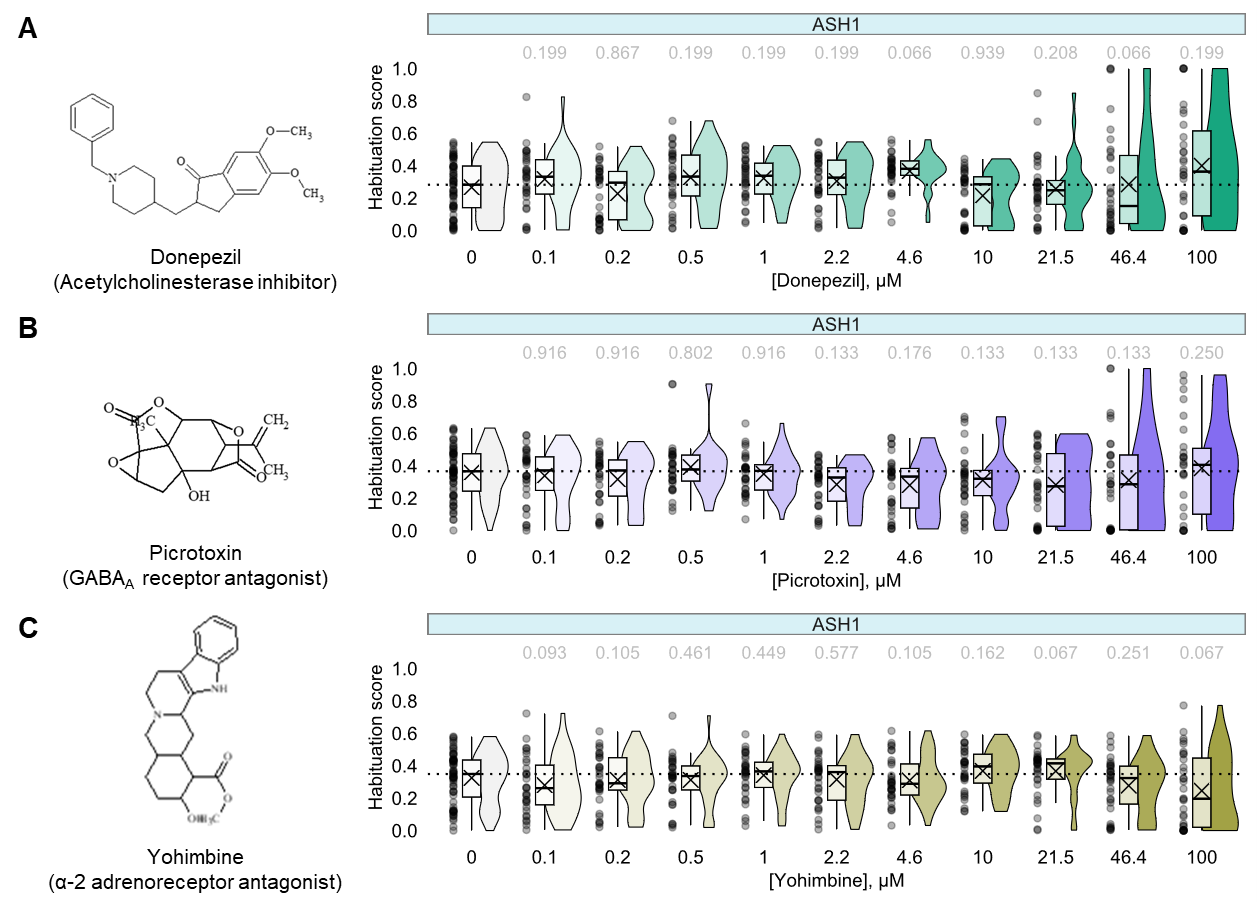

**Figure S3. Habituation scores for reported pharmacological modulators of habituation in zebrafish that did not provoke significant changes in this study.** Chemical structures and habituation scores for (**A**) donepezil, (**B**) yohimbine, and (**C**) picrotoxin. Habituation scores (y-axis) were calculated by dividing the cumulative motor activity during the final ten acoustic stimuli of a sequence of 30 stimuli (300 Hz, 100% volume, 1 s, inter-stimulus interval = 1 s) by the sum of cumulative motor activities during the initial and final ten stimuli per concentration (x-axis). Median habituation scores greater than negative control (grey, horizontal dotted lines) indicate reduced habituation whereas reduced values reflect enhanced habituation. Numbers above the rainclouds represent adjusted p-values (two-sample bootstrapping test, n = 32). Summary data can be found in Excel Tables S47-S49. ASH, acoustic startle habituation.

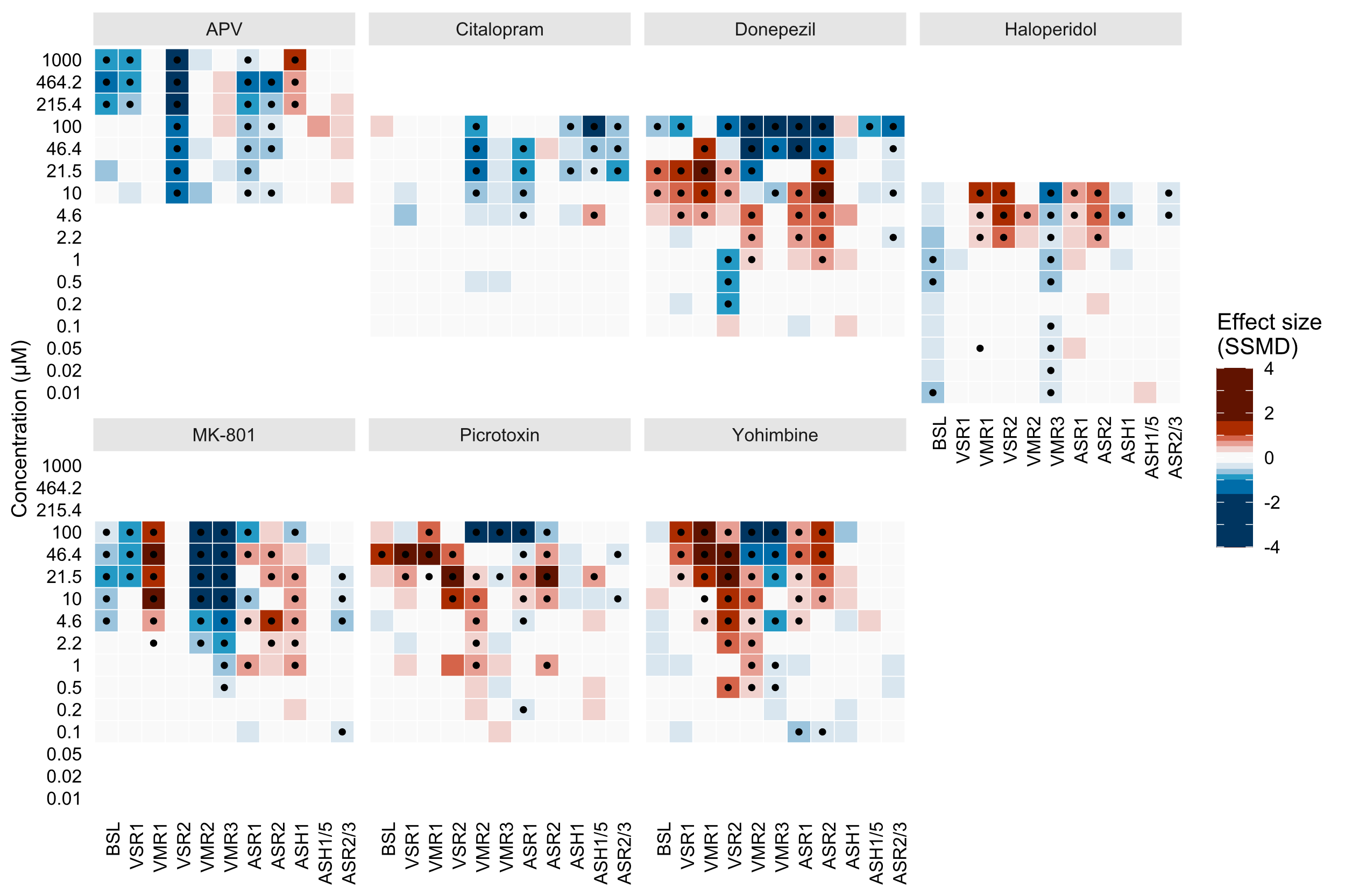

**Figure S4. Behavioral profiles of known pharmacological modulators of habituation in zebrafish.** All 11 behavioral metrics (x-axis) were normalized to vehicle control using strictly standardized median differences (SSMDs) across multiple concentrations (y-axis). Heatmap colors indicate deviation from the median control phenotype: red, more active; blue, less active. Statistically significant changes in behavioral phenotypes are marked with a dot (p<0.05, two-sample bootstrapping test). Summary data can be found in Excel Table S50. APV, (2R)-amino-5-phosphonovaleric acid; BSL, baseline motor activity; VSR, visual startle response; VMR, visual motor response; ASR, acoustic startle response; ASH, acoustic startle habituation.

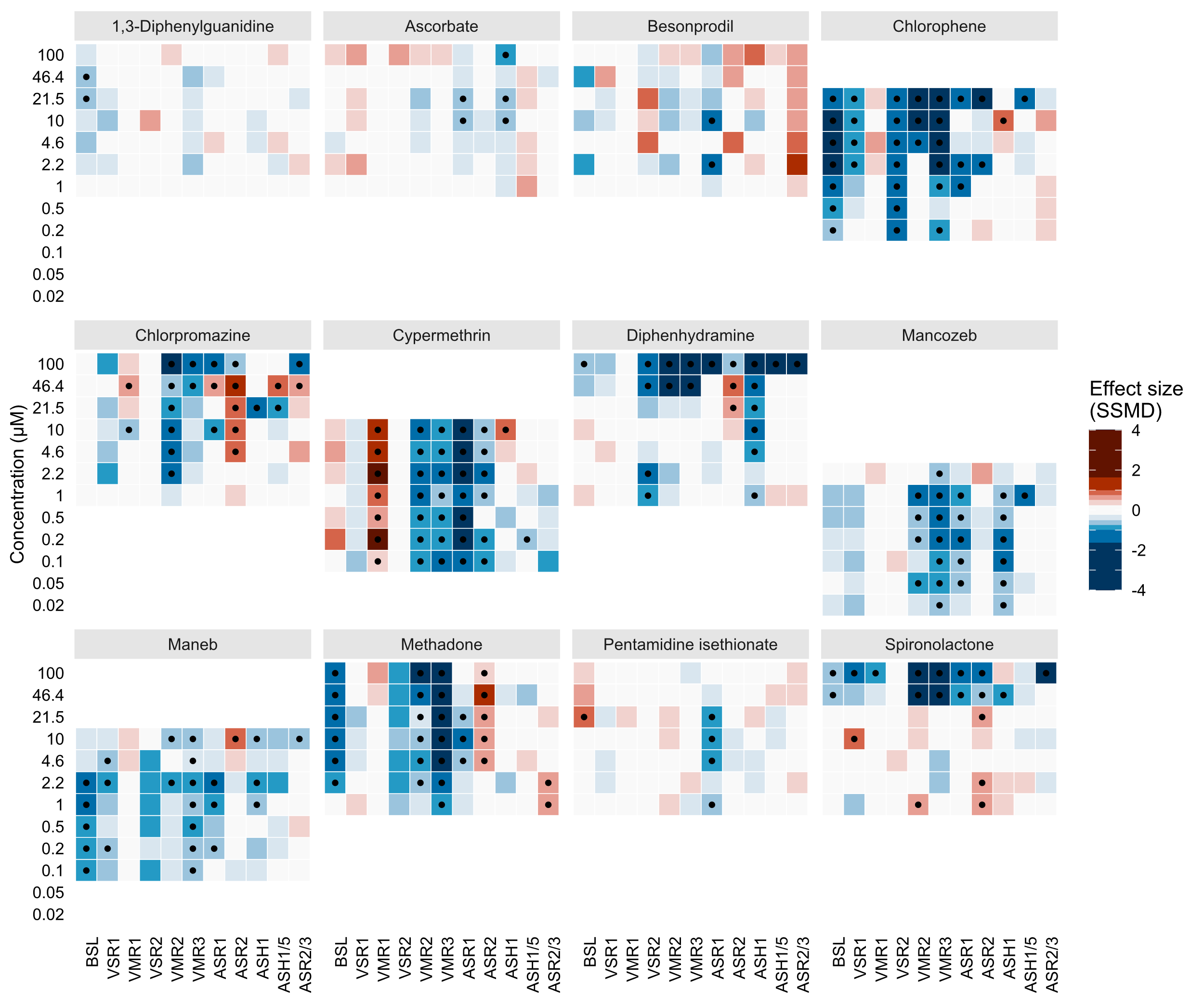

**Figure S5.** **Behavioral profiles of zebrafish treated with chemical compounds that modulate N-methyl-D-aspartate receptor activity in ex vivo rat brain forebrain membrane preparations.** All 11 behavioral metrics (x-axis) were normalized to vehicle control using strictly standardized median differences (SSMDs) across multiple concentrations (y-axis). Heatmap colors indicate deviation from the median control phenotype: red, more active; blue, less active. Statistically significant changes in behavioral phenotypes are marked with a dot (p<0.05, two-sample bootstrapping test). Summary data can be found in Excel Table S5. BSL, baseline motor activity; VSR, visual startle response; VMR, visual motor response; ASR, acoustic startle response; ASH, acoustic startle habituation.

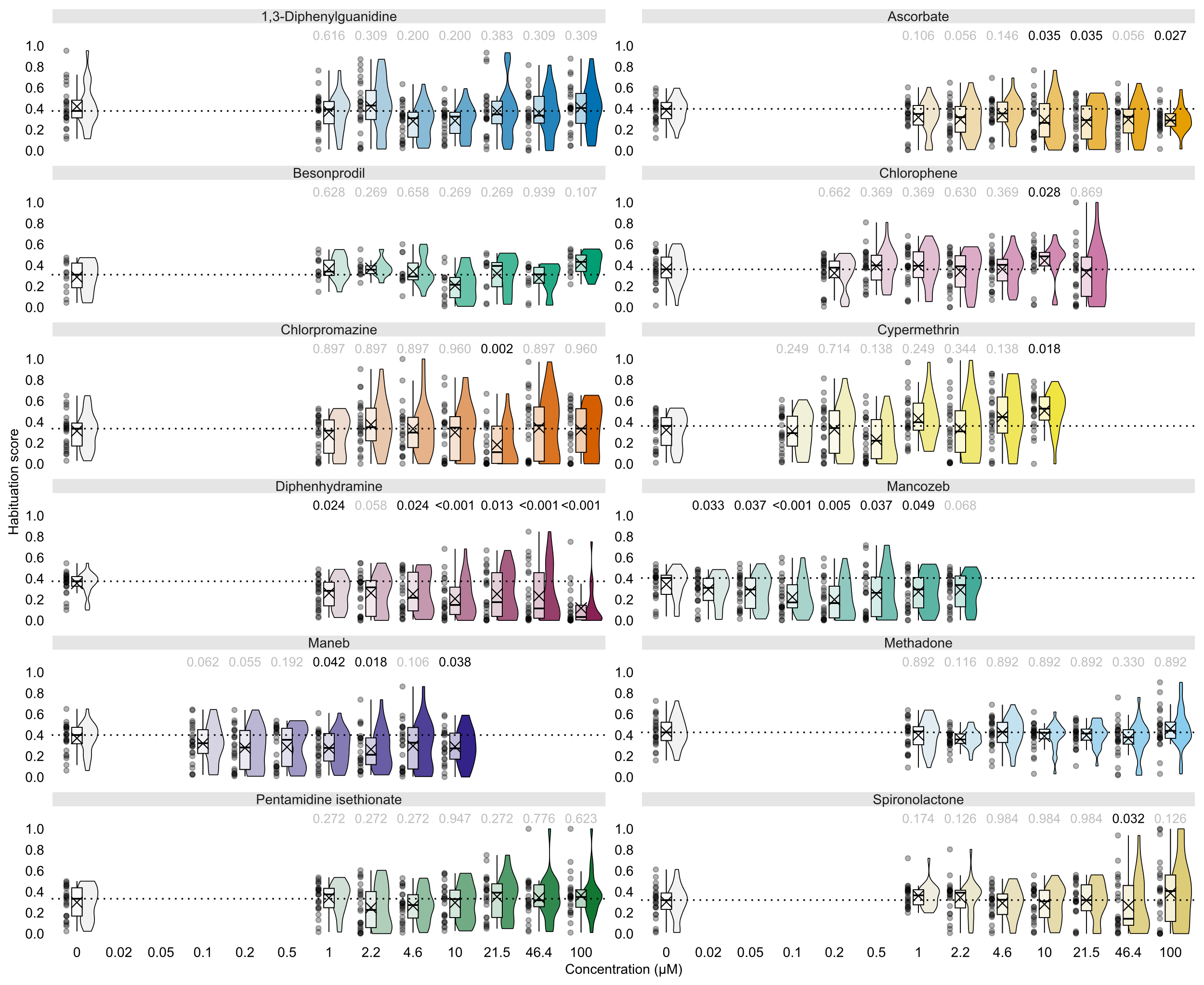

**Figure S6. Habituation scores for zebrafish treated with chemical compounds that modulate N-methyl-D-aspartate receptor activity in ex vivo rat forebrain membrane preparations.** Ascorbate and cypermethrin were selected as negative reference compounds because of their inactivity ex vivo. Habituation scores (y-axis) were calculated by dividing the cumulative motor activity during the final ten acoustic stimuli of a sequence of 30 stimuli (300 Hz, 100% volume, 1 s, inter-stimulus interval = 1 s) by the sum of cumulative motor activities during the initial and final ten stimuli. Chemical compounds and concentrations are indicated on the x-axis. Horizontal dotted lines indicate the median habituation score of untreated larvae. Habituation scores significantly greater than negative control indicate reduced habituation whereas reduced values reflect enhanced habituation. Numbers above the rainclouds represent adjusted p-values (grey: p ≥ 0.05, black: p < 0.05, two-sample bootstrapping test, n = 12-24). Summary data can be found in Excel Table S52. ASH, acoustic startle habituation.

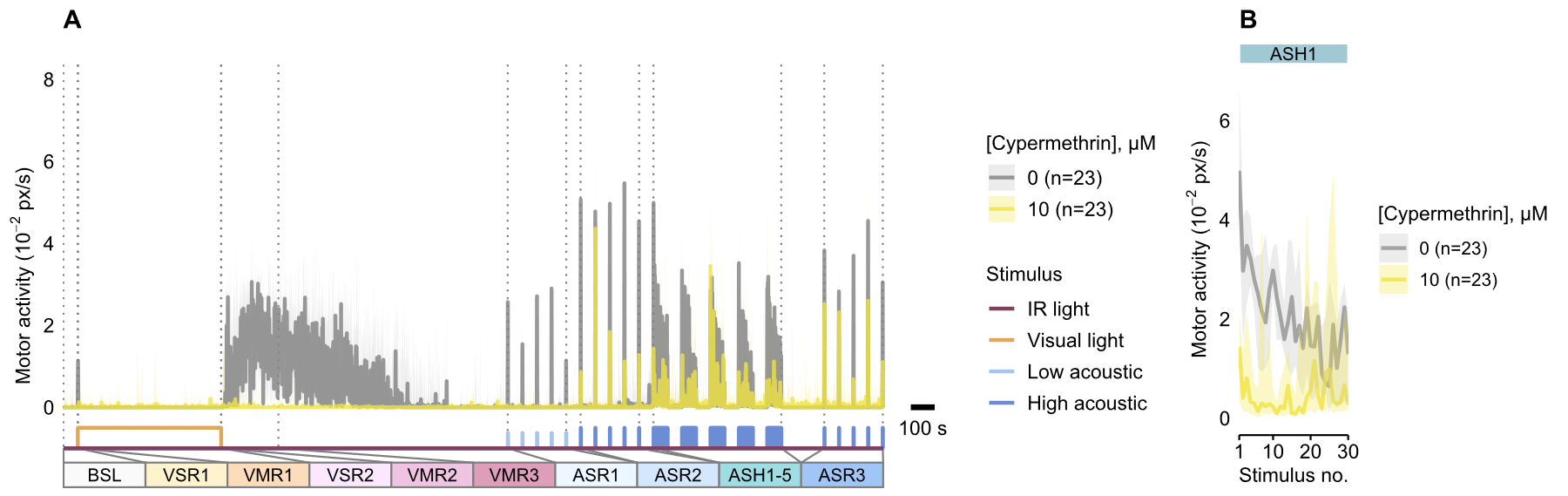

**Figure S7.** **Effects of acute cypermethrin exposure on visual and acoustic behaviors in zebrafish.** (**A**) Motor activity (y-axis) of zebrafish treated with cypermethrin (10 µM, yellow) or 0.01% DMSO (grey) over time and across various stimulus modalities (x-axis). (**B**) Median motor activity (y-axis) of vehicle control (0.01% DMSO, grey) and cypermethrin-exposed (10 µM, yellow) zebrafish larvae over 30 sequential (inter stimulus interval = 1 s) high-intensity acoustic stimuli (x-axis). Data represents median ± 95% CI. Summary data can be found in Excel Table S53. ASH, acoustic startle habituation; ASR, acoustic startle response; BSL, baseline motor activity; VMR, visual motor response; VSR, visual startle response.

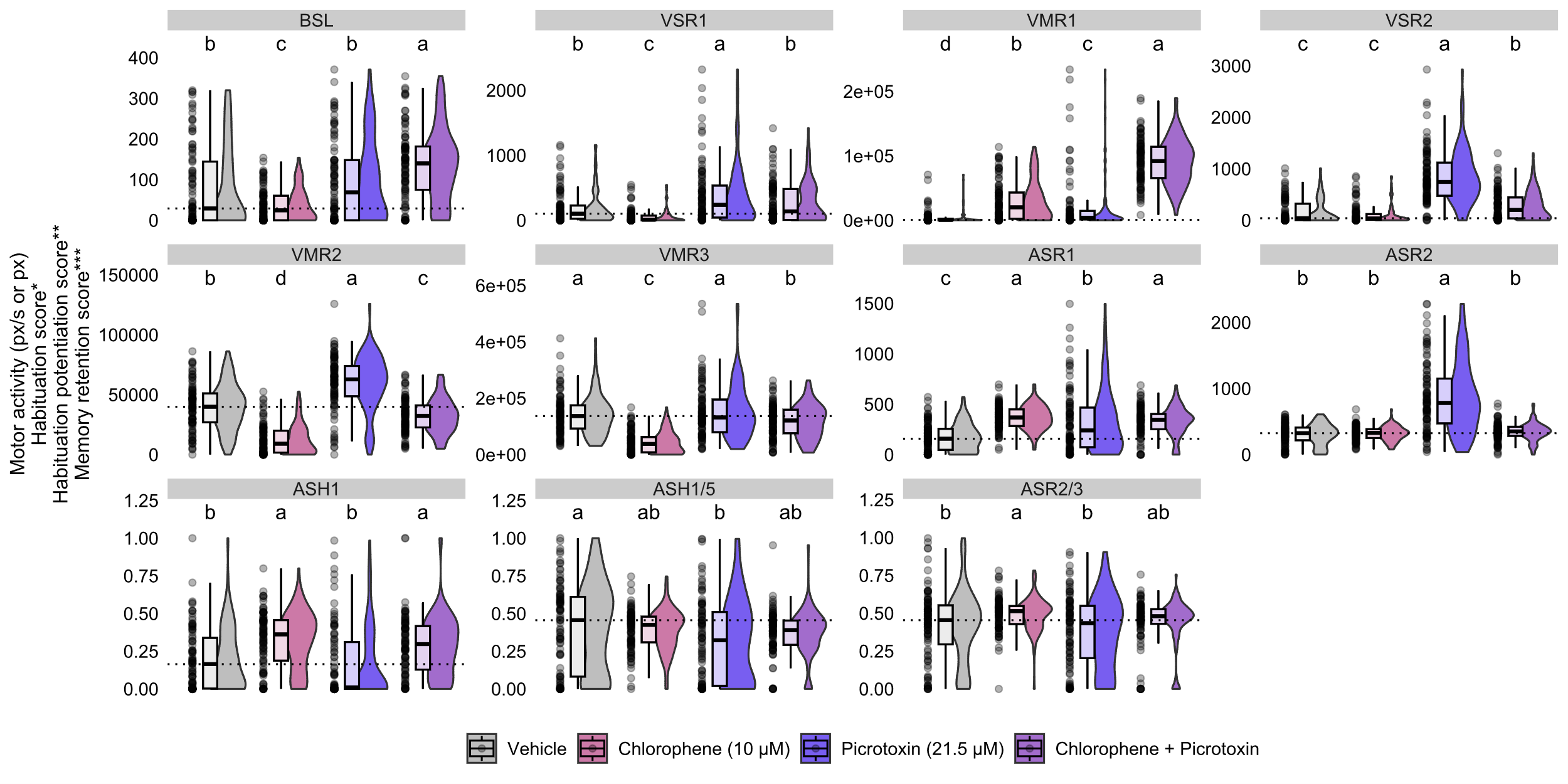

**Figure S8. Effect of the non-competitive GABA_A_ receptor antagonist picrotoxin on chlorophene-induced changes in visual and acoustic behaviors.** The raincloud plots show the locomotor responses (y-axis) to various visual and acoustic stimulus modalities (assay names indicated on top of each plot) in 5 dpf zebrafish treated with the indicated compounds (x-axis). The y-axis represents different response metrics depending on the assay: 'motor activity' for assays BSL-ASR2, 'habituation score' for assay ASH1, 'habituation potentiation score' for assay ASH1/5, and 'memory retention score' for assay ASR2/3. Compact letter displays above rainclouds indicate significant differences between treatments (p<0.05, Kruskal-Wallis rank sum test, n = 94-96 larvae per condition). Summary data can be found in Excel Table S54. BSL, baseline; VSR, visual startle response; VMR, visual motor response; ASR, acoustic startle response; ASH, acoustic startle habituation.

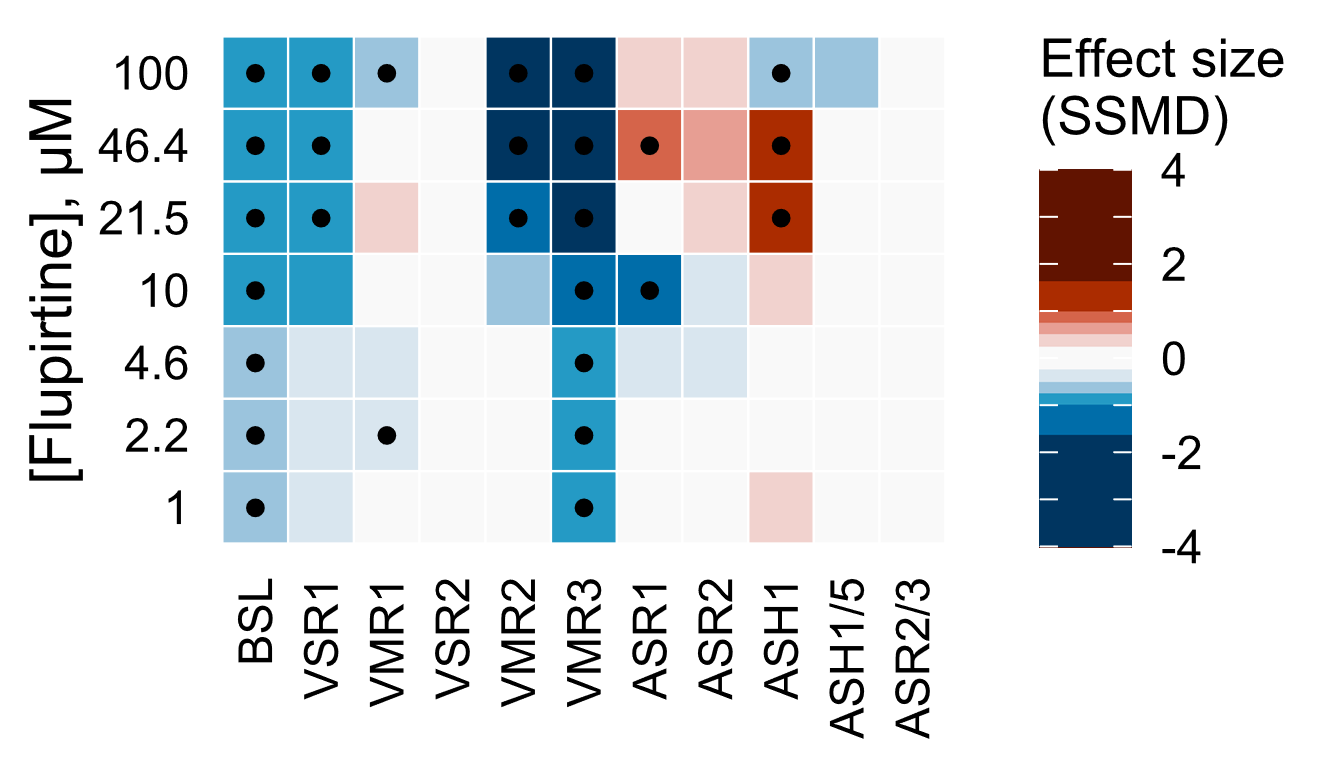

**Figure S9.** **Behavioral profiles of the M-current activator flupirtine.** All 11 behavioral metrics (x-axis) were normalized to vehicle control using strictly standardized median differences (SSMDs) across multiple concentrations (y-axis). Heatmap colors indicate deviation from the median control phenotype: red, more active; blue, less active. Statistically significant changes in behavioral phenotypes are marked with a dot (p<0.05, two-sample bootstrapping test). Summary data can be found in Excel Table S55. BSL, baseline motor activity; VSR, visual startle response; VMR, visual motor response; ASR, acoustic startle response; ASH, acoustic startle habituation.
